# Accurate Determination of Bacterial Abundances in Human Metagenomes Using Full-length 16S Sequencing Reads

**DOI:** 10.1101/228619

**Authors:** Fanny Perraudeau, Sandrine Dudoit, James H. Bullard

## Abstract

DNA sequencing of PCR-amplified marker genes, especially but not limited to the 16S rRNA gene, is perhaps the most common approach for profiling microbial communities. Due to technological constraints of commonly available DNA sequencing, these approaches usually take the form of short reads sequenced from a narrow, targeted variable region, with a corresponding loss of taxonomic resolution relative to the full length marker gene. We use Pacific Biosciences single-molecule, real-time circular consensus sequencing to sequence amplicons spanning the entire length of the 16S rRNA gene. However, this sequencing technology suffers from high sequencing error rate that needs to be addressed in order to take full advantage of the longer sequence. Here, we present a method to model the sequencing error process using a generalized pair hidden Markov chain model and estimate bacterial abundances in microbial samples. We demonstrate, with simulated and real data, that our model and its associated estimation procedure are able to give accurate estimates at the species (or subspecies) level, and is more flexible than existing methods like SImple Non-Bayesian TAXonomy (SINTAX).

## 1 Introduction

Our ability to rapidly and accurately sequence DNA has improved exponentially over the last 30 years. With this improvement, we are beginning to understand the diverse bacterial communities that surround us and how these ecosystems impact biological processes like energy harvest and vitamin synthesis. While we have appreciated bacteria’s capacity to cause infection and illness for over a century, with the introduction of very high-throughput DNA sequencing, we are beginning to understand how the host’s microbiome affects the course of viral and bacterial infections. Beyond these effects, the microbiome plays a key role in proper immune development and has been implicated in immune-mediated disorders like inflammatory bowel diseases and atopic dermatitis.

To study microbial ecosystems, by far, the most popular method is the sequencing of 16S rDNA derived from a polymerase chain reaction (PCR) amplification reaction targeting universal priming sites. Because this method has been used extensively to study a variety of communities, large databases exist with information connecting 16S sequences to taxonomic identifiers and subsequent annotation therein [39].

Beyond the identification of species and the estimation of their abundances in any given biological sample, we would additionally like to understand and characterize the entire genomic repertoire of a microbiome. While the 16S rRNA gene may provide clear taxonomic resolution for a sequence, it may not be able to predict the genomic repertoire of a sample because of issues like, horizontal gene transfer, database incompleteness and bias towards culturable organisms, primer mismatch and bias [7] [30], and inherent ambiguity in the mapping between 16S and a single genome. Despite these limitations, the identification of a strain from a marker gene sequence is a valuable method.

A first step to the construction of a reliable mapping between 16S sequences and genetic capabilities is to produce estimates of a species or strain’s abundance in a sample from 16S sequences. Here, our goal is to define and characterize the performance of a bacterial abundance estimation procedure. Importantly, we seek procedures that provide estimates of uncertainty when estimating the abundance. We want to ensure that uncertainty in assignment is propagated in downstream analyses.

### 1.1 16S rRNA Gene Profiling

#### 1.1.1 Motivation

**Figure 1:**
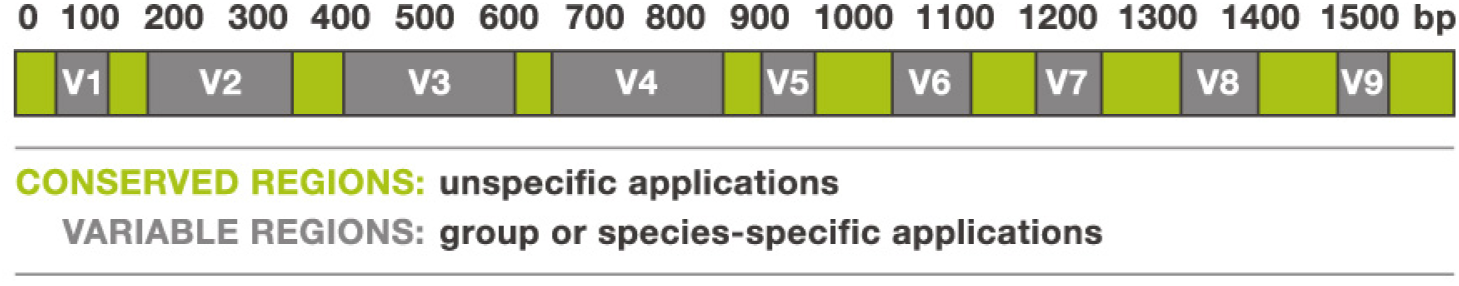
Mosaic structure of the 16S rRNA gene. Conserved regions of the gene are identical for all bacteria, while variable regions contain specific sites unique to individual bacteria. Variable regions enable taxonomic positioning and identification of bacteria, while conserved regions are used to unselectively amplify all bacterial DNA present in a sample [96].

The 16S rRNA gene is the most commonly used genetic marker for classifying bacteria, since it is conserved in all bacteria and has a length large enough (about 1, 500 base-pairs or bp) for discriminative purposes [47]. The distinctive feature of a 16S rRNA gene, which makes it a suitable genetic marker, is its mosaic structure, i.e., alternating conserved and variable regions. Conserved regions of the gene are nearly identical across the majority of characterized bacteria, while variable regions (V1–V9) contain specific sites unique to individual bacterial taxa. Variable regions enable taxonomic positioning and identification of bacteria, while conserved regions are used to non-specifically amplify bacterial DNA present in a sample [96] (see Figure 1). Additionally, large databases of known 16S rRNA gene sequences and primers exist (e.g., Ribosomal Database Project [19], SILVA [72], Greengenes [20]), currently comprising more than three million gene sequences [72].

#### 1.1.2 Issues

Biologists group organisms into taxa, which are assigned a taxonomic rank, whereby groups of a given rank can be aggregated to form a super group of higher rank, thus creating a taxonomic hierarchy. There are commonly eight ranks in biological taxonomy: Domain, kingdom, phylum or division, class, order, family, genus, and species (Figure 2). The ultimate goal of 16S rRNA classification is to identify bacteria at the lowest taxonomic level possible, e.g., species. Although 16S rRNA gene sequencing has been effective at bacterial classification at higher taxonomic rank, e.g., phylum or family, it can perform poorly when attempting to classify at the species level. The degradation in performance for finer grained distinctions occurs for the following three principal reasons:

- **Inter-species similarity.** Two bacteria from different species or even genera can have very similar 16S rRNA genes. For example, Yassin et al. [93] reported that *Nocardia brevicatena* and *Nocardia paucivorans* show 99.6% 16S rRNA gene sequence similarity. However, the results of DNA-DNA hybridization studies^1^ and phenotypic testing indicate that they are distinct species. This is not a novel finding, as Fox [33] remarked in their 1992 publication that 16S rRNA sequence identity may not be sufficient to guarantee species identity, especially for recently diverged species.
- **Intra-species heterogeneity.** One bacterium can have between 1 and 15 different 16S genes, i.e., loci with different 16S gene sequences or multiple copies of a given gene sequence. For instance, for the University of Michigan Ribosomal RNA Database, the median and mode for the number of 16S genes for the bacteria in the database are respectively 4 and 2 [84]. Even if there is a strong pressure to maintain a high level of conservation for the 16S rRNA gene, intra-genomic heterogeneity exists, where the average number of nucleotides that differ between any pair of 16S rRNA genes within a genome is 2.91 (standard deviation 4.78) and the corresponding average similarity is 99.81% (standard deviation 0. 31) [18]. This intra-species heterogeneity is especially problematic for bacterial classification, as the various sequences of the 16S rRNA gene in a single organism can be more different from each other than those in different organisms.
- **Genotype doesn’t correspond to phenotype.** Phylogenetic classification using 16S rRNA gene sequences largely relies on the assumption that 16S rRNA genes are only vertically inherited from parent to offspring and thus belong exclusively to a given species. However, microbial genomic analysis has revealed that some bacteria contain 16S rRNA genes that are mosaics of sequences from multiple species (this phenomenon is called horizontal gene transfer), suggesting that 16S rRNA can be transferred between different species [52]. Assuming that these mosaics are not in fact chimeras (i.e. artifacts made during the PCR process), classification of bacteria based on the 16S rRNA gene should be carefully interpreted. Additionally, virulence or phenotypically relevant genes are probably horizontally transferred so if these horizontally transferred genes are the drivers of phenotype, then using a vertically inherited gene would be problematic.

### 1.2 16S rRNA Gene Sequencing

#### 1.2.1 Sequencing technologies

In early bacterial community studies, near full-length 16S rRNA genes were sequenced using the Sanger technology. This approach, though informative, was time-consuming, expensive, and provided a limited depth of sequencing, which was insufficient to uncover the complete bacterial diversity present in a complex sample [69] [17] (Table 1). Moreover, the Sanger technology is not capable of accurately sequencing mixtures, but rather demands an isogenic population of molecules, which is incompatible with 16S rRNA studies.

**Figure 2:**
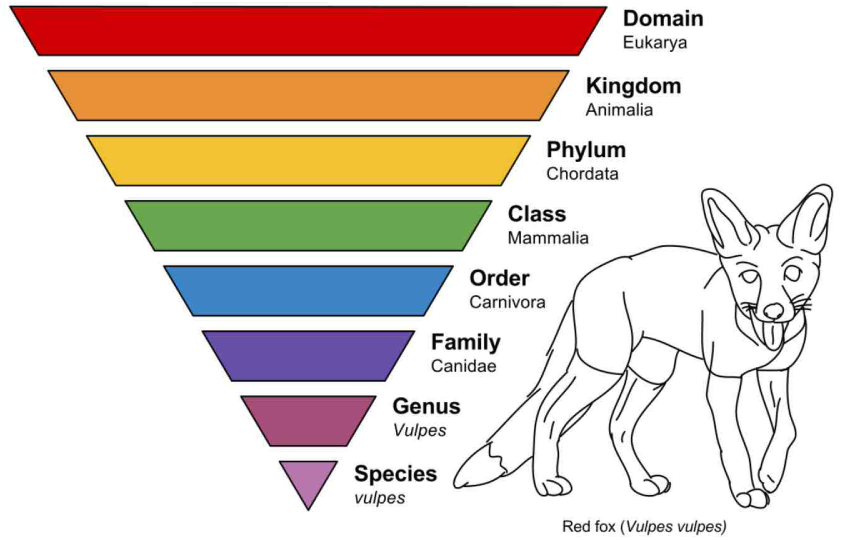
Biological classification. Main taxonomic ranks: Domain, kingdom, phylum, class, order, family, genus, and species. In this example, taxonomic ranking is used to classify animals and earlier life forms related to the red fox, *Vulpes vulpes* [94].

In contrast to Sanger sequencing, second-generation sequencing technologies, e.g., Illumina, provide much higher sequencing depth and allow sequencing of complex mixtures [22]. The latter have been widely used to assess the composition of microbial communities, enabling the completion of high-profile microbiome projects, such as the Human Microbiome Project (HMP) [46] [63]. However, the higher throughput comes at the expense of read length; current technologies produce only partial sequences of 16S rRNA genes, with length varying from 250 (Illumina MiSeq) to 500 base-pairs (Roche 454). Therefore, most studies using second-generation sequencing technologies have to select the most informative variable regions (V1–V9, see Figure 1) as phylogenetically informative markers to identify taxa. Such partial 16S rRNA gene sequencing can bias estimates of diversity, since nucleotide differences are not evenly distributed along 16S rRNA genes [50].

Thus, the recent development of third-generation sequencing technologies offers a promising approach for high-resolution analysis of microbial communities (Table 1). By virtue of sequencing single contiguous molecules, third-generation technologies are capable of producing long reads. Unfortunately, with current technology, gains in length can come at the expense of accuracy. In particular, the basecall error rate for the current PacBio RS sequencer is around 10%, whereas second-generation technologies like Illumina produce basecalls with error rates around 0.02%. Since our aim is to distinguish between species with sequence similarity around 99%, the high sequencing error rate is a problem. To solve this problem, we use high-coverage single-molecule consensus reads as input to our classification algorithm. This classifier is trained off-line on a training set where reads have been annotated. It means that for each read, we known from which bacteria it has been sequenced from.

#### 1.2.2 Pacific Biosciences’ single-molecule, real-time circular consensus sequencing

The first step in Pacific Biosciences’ (PacBio) single-molecule, real-time (SMRT) circular consensus sequencing (CCS) process is to amplify full-length 16S rRNA genes from metagenomic DNA samples using polymerase chain reaction, with 16S rRNA gene-specific primers. The resulting PCR products, called amplicons, are then converted into a circular form, called SMRTbell, by ligation of hairpin adaptors to both ends of the double-stranded linear amplicons. A hairpin adaptor contains a sequence complementary to a primer, where a DNA polymerase can bind forming a sequencing-productive complex [87].

**Table 1:**
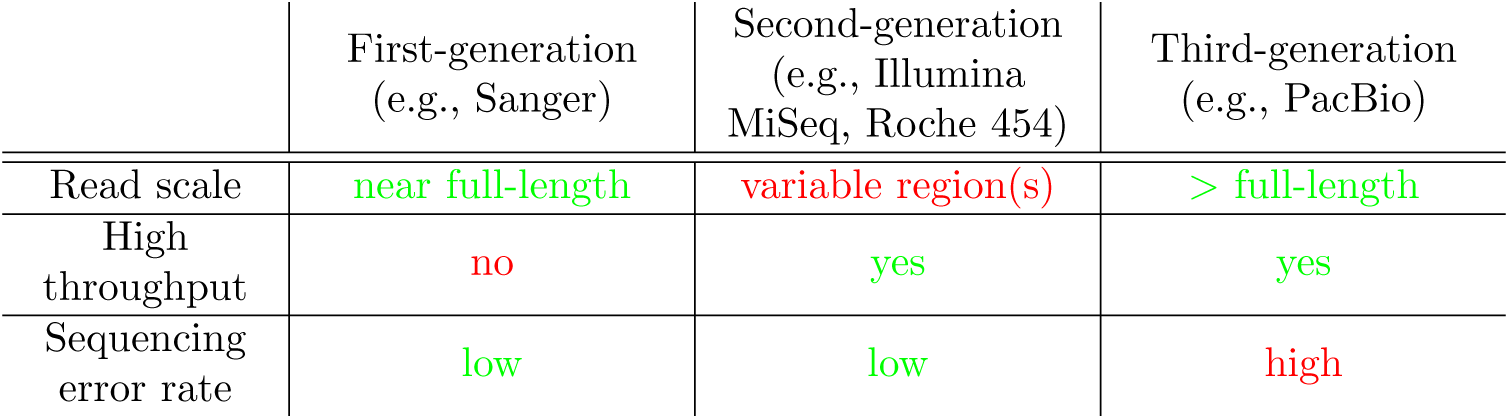
Overview of the three generations of sequencing technologies in the context of 16S rRNA gene sequencing. First-generation sequencing allows sequencing of near full-length 16S rRNA gene sequences. However, low throughput limits power to analyze microbial communities. Second-generation sequencing is currently the most used technology, as it has high throughput and low sequencing error rate. However, the short length of the sequencing reads makes it necessary to select variable regions as informative markers to identify taxa, limiting phylogenetic power. Third-generation sequencing seems promising for increasing phylogenetic resolution, as reads spanning the entire length of 16S rRNA genes can be sequenced. However, third-generation technologies have high sequencing error rates.

|  | First-generation<br>(e.g., Sanger) | Second-generation<br>(e.g., Illumina<br>MiSeq, Roche 454) | Third-generation<br>(e.g., PacBio) |
| --- | --- | --- | --- |
| Read scale | near full-length | variable region(s) | > full-length |
| High throughput | no | yes | yes |
| Sequencing error rate | low | low | high |

Essentially, the sequencing platform takes a video of the polymerase adding fluorescent nucleotides to the template. Each nucleotide {*A, C, G, T*} is associated with a different fluorescent color, allowing the optical system to distinguish between different nucleotides. The sequencing process stops when the circular template either falls off or is killed as a side-effect of the fluorescent excitation, or when the polymerase dies. If the amplicon is short enough, the polymerase will have time to go around the hairpin multiple times, producing a sequencing read with multiple observations for each base. These multiple observations can then be used to generate high-accuracy consensus sequences for each 16S rRNA gene.

Circular consensus sequencing (CCS), enabling a consensus sequence to be obtained from multiple passes on a single template, overcomes the high single-pass error rate of the PacBio SMRT technology. While the overall single-pass sequencing error rate is estimated at about 15% [54], in the circular consensus sequencing mode, the error rate depends on the number of passes of the polymerase on the template (i.e., the number of sequenced nucleotides for each unique base in the template). With the PacBio RS II sequencer and the P4/C2 chemistry, the mean length of the sequencing reads is of about 6, 900 bp. As 16S rRNA genes are about 1, 500 bp long, most of the raw long reads are sequenced from at least 3 passes of the polymerase on the template. This results in an overall sequencing error rate of about 2% for a typical dataset for 16S rRNA genes.

**Figure 3:**
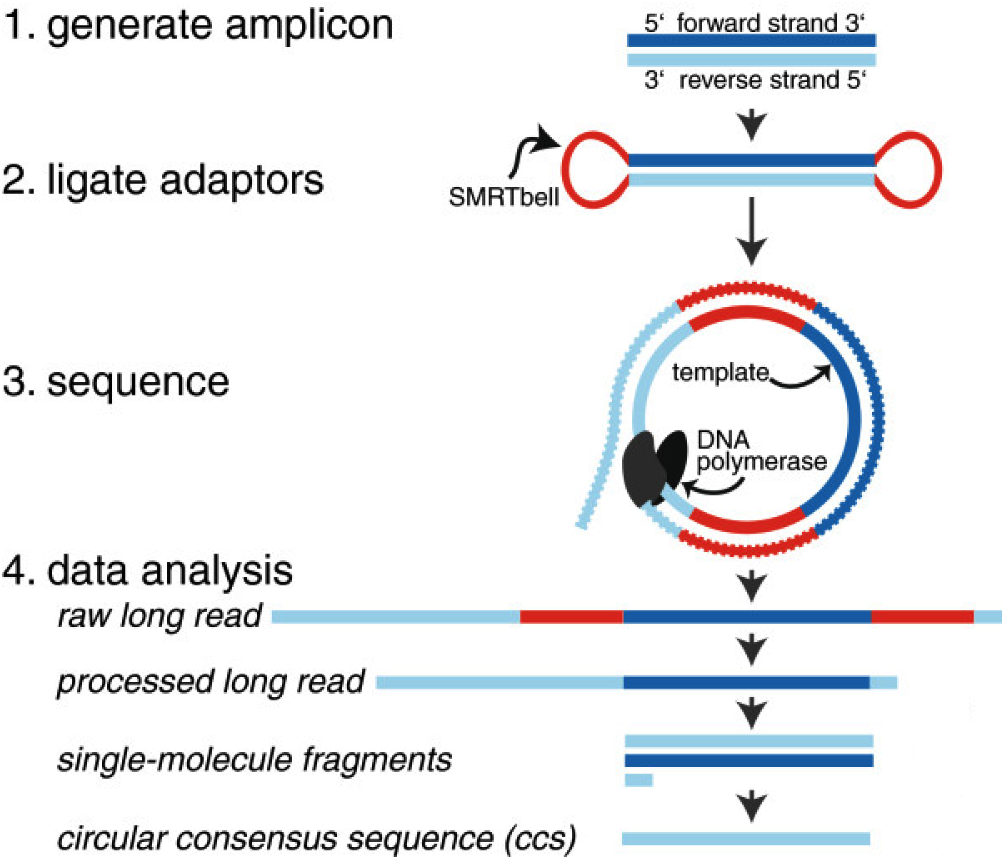
Illustration of PacBio’s single-molecule, real-time circular consensus sequencing. Hairpin adaptors are ligated to double-stranded DNA, e.g., PCR amplicons or shotgun library. This construct is known as a SMRTbell and a library of these will be loaded onto a single chip. The PacBio RS II observes each polymerase as it sequences copies of the SMRTbell. By circularizing the DNA when constructing the library, the instrument is able to collect many measurements of each base in its forward and reverse complement context. After combining these individual base-pair measurements, the RS II software produces a consensus sequence, i.e., best estimate of the original double-stranded DNA molecule [32].

### 1.3 Review of Current Statistical Methods

#### 1.3.1 Cluster-based methods

Most current statistical methods used to analyze microbial communities are based on clustering and comprise two steps.

##### Step 1. Clustering 16S rRNA gene sequencing reads

###### OTUs

In the first step, the reads sequenced from a 16S rRNA gene are clustered according to their sequence similarity. The idea is to compare all pairs of sequences and group similar sequences together. As the number of reads is large, performing all pairwise comparisons is intractable, thus greedy algorithms are used instead. Specifically, a sequence is selected to constitute the seed of the first cluster using some criterion (e.g., the longest sequence of the dataset is selected). Next, remaining sequences are compared with the seed. If its similarity with the seed meets a predefined cutoff (often 97%), a sequence is grouped into that cluster; otherwise, it becomes the seed of a new cluster. The process is repeated until all reads are clustered. Examples of commonly-used algorithms include UPARSE [27], CD-HIT [36], UCLUST [24], and DNACLUST [38].

The resulting clusters, called operational taxonomic units (OTU), collapse the complete set of reads into a smaller collection of representative sequences (one for each OTU), where reads within an OTU have a similarity higher than a fixed threshold. The commonly-used threshold is 97% and would correspond to clustering at the species level. However, it remains debatable how well this method recapitulates a true microbial phylogeny, as it both overestimates diversity when there are more sequencing errors than the OTU-defining threshold and cannot resolve real diversity at a scale finer than that threshold [74].

###### Exact-sequence methods

Recently, new methods have been developed that resolve amplicon sequence variants (ASVs) from amplicon data without imposing the arbitrary similarity threshold that define OTUs. These methods are also referred as exact-sequence methods. ASVs are inferred by a *de novo* process in which biological sequences are discriminated from errors on the basis of, in part, the expectation that biological sequences are more likely to be repeatedly observed than are error-containing sequences [13]. The most commonly used tools for exact-sequence methods are DADA2 [14] (offered in QIIME2 [95]), UNOISE2 [26] (offered in USEARCH [24]), and Deblur [4] (offered in QIIME2 [95]).

##### Step 2. Labeling clusters using a reference database

In the second step, a sequence per cluster (e.g., the seed used during the clustering step) can be defined to be the cluster representative and corresponding abundances are computed based on the number of reads falling within each cluster. Then, the sequence of the cluster representative is compared to all the 16S rRNA sequences present in a reference database [17], such as the Ribosomal Database Project, Greengenes, or SILVA, where each sequence in these reference databases is annotated with its taxonomic rank. Finally, each cluster representative is assigned the same taxonomic rank as its most similar sequence in the reference database. The most commonly used tools are the RDP-Classifier, which uses a naive Bayesian classifier [90], UTAX [28], and PyNAST [15].

###### Pipelines

Several pipelines can be used to perform the above two steps [70]. Commonly-used pipelines include QIIME [16], Mothur [80], and MG-RAST [64]. For example, QIIME uses UCLUST as the default algorithm to cluster reads into OTUs, then the most abundant read in each OTU is selected as the representative sequence, and the script `Assign_taxonomy.py` is used for the classification of each of the representative sequences.

#### 1.3.2 Closed-reference methods

The methods developed to label cluster representatives using a reference database can also be used to label each read in a dataset. The benefit of such an approach is that there is no need to select an arbitrary similarity cutoff (often 97%) to build OTUs. However, such a computation can be time-consuming and require a lot of memory, especially when the method has not been designed to handle long reads. For example, to build a reference database using our own reference sequences (about 80, 000 sequences, about 1, 500bp-long each) using the software UTAX, with the command *usearch* with argument *makeudb_tax*, requires more than 4Gb of memory. Unfortunately, the free 32-bit version of UTAX cannot handle more than 4Gb of memory; a 64-bit version is available with a paid license. Thus, it was impossible for us to use the free version of this tool to classify long reads.

**Table 2:** Comparison between current commonly-used statistical methods and our method. Most studies of microbial communities based on 16S rRNA genes use second-generation sequencing to generate short reads spanning only portions of a 16S rRNA gene and consist of two steps. In the first step, similar reads are grouped into clusters, called Operational Taxonomic Units (OTUs), while in the second step each cluster is labeled with taxonomic rank. In our method, reads spanning the entire length of a 16S rRNA gene are sequenced using PacBio’s single-molecule, real-time circular consensus sequencing. Each read is then assigned a taxonomic rank by comparison with annotated sequences in a pre-filtered reference database.

|  | Cluster-based | Closed<br>reference-based | Proposed method |
| --- | --- | --- | --- |
| Read length | short | short/long | long |
| Initial step | pre-clustering of<br>the reads into<br>OTUs | none | pre-filtering of<br>reference sequences |
| Labeling | cluster<br>representatives | reads | reads |

Recently though, a few tools have been designed for long reads. For example, oneCodex [65] and Simple Non-Bayesian TAXonomy (SINTAX) [29] were released, respectively, in 2015 and 2016. OneCodex is fast and easy to use. However, we could not apply this method to our data, as it does not allow the use of a reference database different from either the NCBI RefSeq database or their own in-house reference database. On the other hand, with SINTAX it is easy to provide a reference database and there is no need to train the algorithm (while training is required when using UTAX or RDP-Classifier). However, SINTAX has a low sensitivity, that is, some bacteria known to be present in the sample are not detected (see Section 4).

### 1.4 Our Approach

Most studies using the 16S rRNA gene to analyze microbial communities rely on second-generation sequencing. However, the short reads yielded by these technologies only allow consideration of a few variable regions as phylogenetically informative markers to identify taxa. Such partial 16S rRNA gene sequencing can bias estimates of diversity, since nucleotide differences are not evenly distributed along the 16S rRNA gene. Instead, we use third-generation sequencing, more specifically PacBio’s single-molecule, real-time circular consensus sequencing, to sequence reads spanning the entire length of the 16S rRNA gene (Table 2). As standard tools were designed for short reads, there is a need to develop new statistical methods for long reads.

Sequencing full-length 16S rRNA genes has the potential to provide a higher phylogenetic resolution than short-read sequencing (at a finer level than the 97% threshold commonly used to define OTU), as it is no longer necessary to target specific variable regions. To take full advantage of long reads, our method does not group reads into OTUs; instead, each read is assigned the same taxonomic level as its most similar sequence in a reference database. As the number of sequences in a reference database is typically on the order of millions and the number of sequencing reads on the order of twenty thousand, such a computation is intractable. To reduce the computation, reference sequences with a small probability of having generated the reads in a dataset are eliminated in a pre-filtering step. Then, the probability that each read could have been sequenced from the remaining sequences in the reduced reference database is computed. Finally, the read is assigned the same taxonomic rank as the reference sequence with which it has the greatest probability. This probabilistic framework, using sequencing reads spanning the entire length of 16S rRNA genes, allows us to determine accurately bacterial relative abundances at the species (or subspecies) level.

### 1.5 Statistical Inference Framework

#### 1.5.1 Population and parameters of interest

Consider a population of *M* (*M* ⋍ 10^12^) bacteria (e.g., a patient’s gut microbiome [82]), where the bacteria are of *K* different types, 𝓑 = {*b_k_*: *k* = 1,…, *K*}. Let π = (π_*k*_ : *k* = 1,…, *K*) denote the population frequencies for each of the *K* bacteria in 𝓑. Our goal is to estimate the parameter π = (π_*k*_ : *k* = 1,…, *K*). In some cases, we may also wish to estimate a function of π or test hypotheses about π (e.g., test for each *k* whether π_*k*_ > *є*, where *є* > 0 represents the detection limit of the system).

Although beyond the scope of this report, a problem of great interest is the comparison of two bacterial populations, i.e., the identification of bacteria *k* which are present at different frequencies in the two populations (cf. differential expression in high-throughput microarray and sequencing assays). This involves testing for each *k* the null hypothesis that 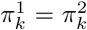, where π^1^ and π^2^ denote, respectively, the bacterial frequencies in the first and second population.

#### 1.5.2 Reference database

We rely on an in-house annotated reference database

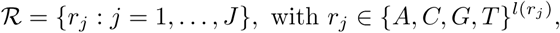

where *l* is the function mapping a sequence of nucleotides to its length (see Section 2.2). The *J* (*J* ⋍ 1.4 × 10^6^) reference sequences in 𝓡 are all the 16S genes extracted from the bacteria in 𝓑, i.e., it is assumed that all bacteria in 𝓑 are represented in 𝓡. Note that one bacterium can have 16S genes at different loci in its genome, with either identical sequences or slightly different sequences (at only a handful of bases). In other words, a given bacterium can have, at different loci, either multiple identical *copies* of a 16S gene or multiple *variants* of a 16S gene. Moreover, two different bacteria can have the same 16S gene. Given this setting, we define a *K* × *J* matrix *C*, where *C_kj_* designates the number of copies of reference sequence *r_j_* in bacterium *b_k_*. In particular, we let the mapping *b*(*r_j_*) denote the set of all bacteria containing at least one copy of *r_j_*, i.e.,

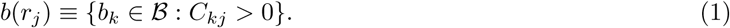

#### 1.5.3 Data generation model

In order to infer the bacterial population frequencies π = (π_*k*_ : *k* = 1,…, *K*) (or functions of these frequencies), we adopt the following three-step data generation model.

- **Step 1. Sampling bacteria**. Sample *m* (a few thousand) bacteria at random (without replacement) from the population of interest and denote by *Ƶ* = {*Z_i_* : *Z_i_* ∈ **𝓑**; *i* = 1; …*m*} the resulting set of bacteria (Figure 4). For simplicity, we ignore the presence of eukaryotic and viral cells and we assume that these cells have no effect on subsequent steps. For large *M* and small m compared to *M*, sampling without replacement can be treated as sampling with replacement, so that the sample frequencies for each of the *K* bacteria in 𝓑 follow a multinomial distribution, that is,

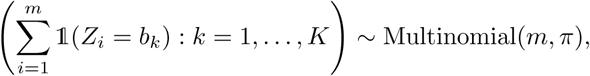

where 𝟙(·) is the indicator function, equal to one if its argument is true and zero otherwise. In particular, Pr(*Z_i_* = *b_k_*) = *π_k_*.

**Figure 4:**
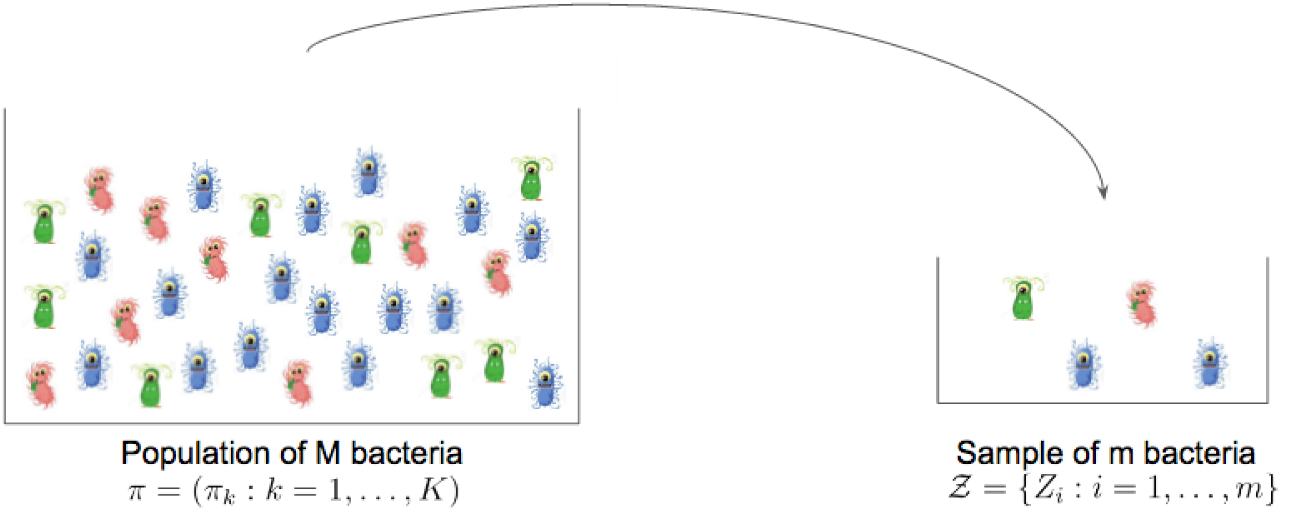
Step 1. Sampling bacteria. Simple random sample of m bacteria from a population of M bacteria. For simplicity, we ignore the presence of eukaryotic and viral cells in the figure. We assume that these cells have no effect on subsequent steps.

- **Step 2. Sampling 16S amplicons**. For each sampled bacterium in *Ƶ*, extract all of its 16S genes and amplify them using polymerase chain reaction (PCR). In what follows, we refer to the amplified 16S gene sequences as *amplicons.* Select at random (without replacement) n of these amplicons (pooled from all m sampled bacteria), 𝓧 = {*X_i_* : *X_i_* ∈ 𝓡, *i* = 1,…, *n*} (Figure 5). As cell lysis performance varies from one bacterium to another, we define *e*(*b_k_*) as the chance that DNA is extracted from bacterium *b_k_*. Additionally, as different sequences are amplified with varying efficiencies, not all 16S genes have the same chance of being sampled. For a sequence *r_j_* with PCR efficiency *e*(*r_j_*), *c* PCR cycles yield approximately (2*e*(*r_j_*))^*c*^ copies of the sequence. Hence, the probability of selecting amplicon *X* = *r_j_* for bacterium *Z* = *b_k_* is

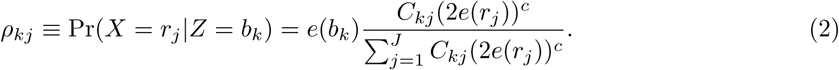 Here, we will assume that PCR makes no errors. Without this assumption, the amplicons may not belong to 𝓡. Additionally, we assume that each sampled bacterium contains all copies of its 16S genes represented in 𝓡, i.e., all *C_kj_* copies of *r_j_* ∈ 𝓡 when *Z* = *b_k_*. DNA extraction and PCR efficiencies will be estimated in an upcoming experiment at Whole Biome.

**Figure 5:**
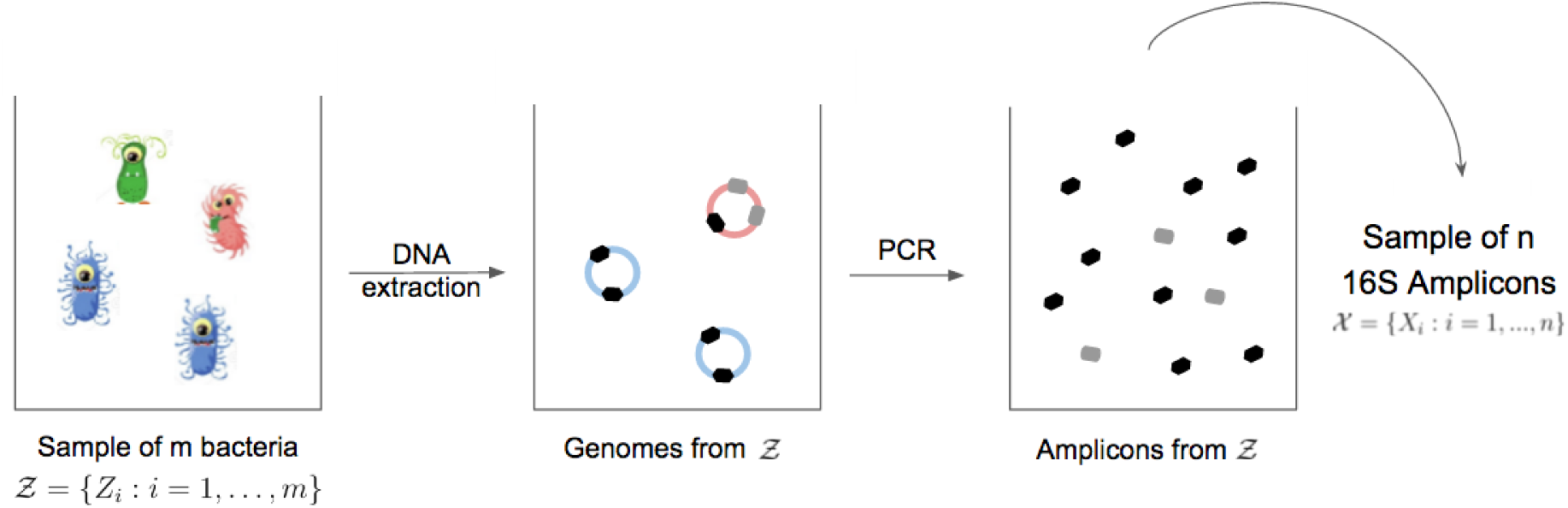
Step 2. Sampling 16S amplicons. Sample *n* amplicons at random (without replacement) from the amplicons of the bacteria in *Ƶ*. Here, four bacteria were sampled (two blue, one green, and one pink). DNA extraction did not work for the green bacterium. The blue bacterium has two identical copies of its 16S gene (black), while the pink bacterium has two different variants of its 16S gene (black and gray), with one variant (gray) having two identical copies. Hence, for the pink bacterium, there are three distinct loci where amplification of the 16S gene can occur. The blue and pink bacteria share the same 16S gene sequence (black variant). One cycle of PCR is represented and PCR worked only partially for the gray variant.

- **Step 3. Sequencing 16S amplicons.** Sequence each of the 16S amplicons in 𝓧, using the PacBio SMRT platform, to obtain a set of *n* reads

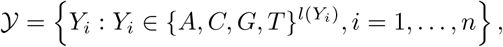

where *Y_i_* is the sequencing read corresponding to amplicon *X_i_* and *l*(*Y_i_*) ⋍ 1; 500 bp. If amplicons in 𝓧 were sequenced without error, each read in 𝒚 could be matched to a reference sequence in 𝓡. However, errors occur during the sequencing process, introducing noise in the data. Essentially, the sequencing platform takes a video of a polymerase adding uorescent nucleotides to a template. Each nucleotide {*A, C, G, T*} is associated with a different uorescent color, allowing the optical system to distinguish between different nucleotides. As this process happens quickly – in real time, actually – the imaging system might randomly make mistakes (i.e., mismatches), skip, or add a nucleotide. Each sequence of nucleotides *Y* we get to observe can then be different from the true sequence of nucleotides *X*. Viewing the error generation as stochastic, we can consider the probability Pr(*Y* = *y*|*X* = *x*) that a noisy read *Y* was generated from a true sequence *X*. We will estimate such probabilities using the generalized pair hidden Markov model (GPHMM) presented in Section 3.3.1 and we further assume that errors are introduced independently between reads [73].

Our three-step data generation model is equivalent to the graphical model in Figure 6, whereby, for each *i* = 1, …, *n*,

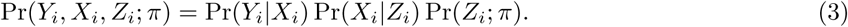

However, only the reads *Y* are observed and the probability of a read *Y_i_* is

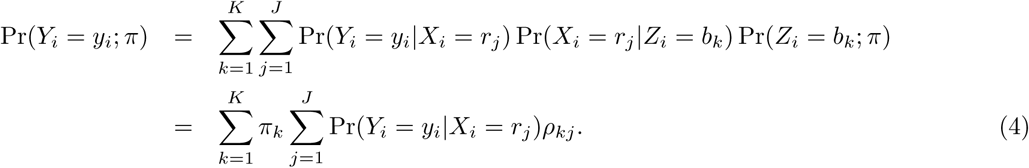

**Figure 6:**
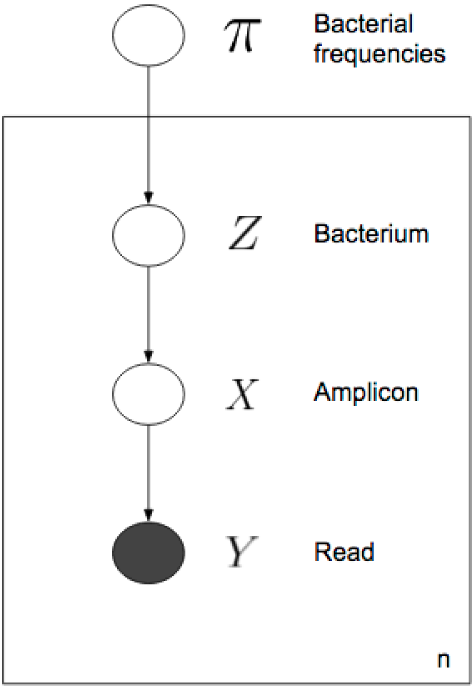
Data generation model. Graphical model representation of generative model for read dataset 𝒚. To generate each read *Y* ∈ 𝒚, sample a bacterium *Z* at random from a microbiomic population of interest with bacterial frequencies π. For each sampled bacterium *Z*, extract all of its 16S genes, amplify them using PCR, and select at random one of these 16S amplicons, *X*. Finally, sequence amplicon X to generate read Y. The process is repeated independently for each of the *n* reads in 𝒚. Only the shaded node is observed.

Given that each read *Y_i_* in 𝒚 is generated independently and from Equation (4), the log-likelihood of the data 𝒚 is

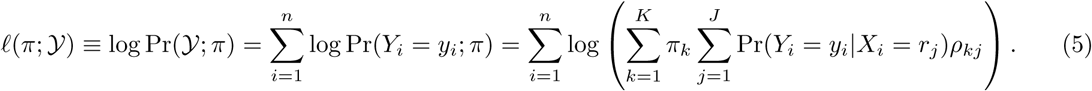

A natural estimator of the bacterial frequencies π is the maximum likelihood estimator (MLE), defined as

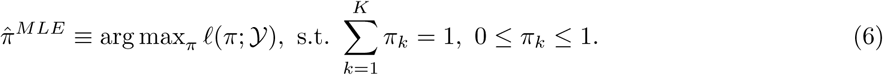

The naive way to compute the log-likelihood 𝓁(π; 𝒚) would be to compute Pr(*Y_i_* = *y_i_*|*X_i_* = *r_j_*) for each of the *n* reads *Y_i_* in 𝒚 and each of the *J* reference sequences *r_j_* in 𝓡. The number of reference sequences in 𝓡 being on the order of 1.4 million and the number of reads in 𝒚 on the order of the thousands, such a computation is intractable. Thus, our reference database 𝓡 is first reduced to a smaller database of a few thousand sequences (the exact number depending on the dataset 𝒚), by eliminating the reference sequences with a small probability of having generated the reads in 𝒚. This step is performed using the alignment tool Bowtie2 [56] (Section 3.2). Secondly, the probabilities Pr(*Y_i_* = *y_i_*|*X_i_* = *r_j_*), for a reduced version of the database 𝓡, are computed using the Viterbi algorithm for a generalized pair hidden Markov model (Section 3.3). Given estimates of Pr(*Y_i_* = *y_i_*|*X_i_* = *r_j_*) and *ρ_kj_*, Section 3.4 describes various approaches for estimating π. This process is summarized in Figure 7 and, in greater detail, in the workflow of Figure 15.

**Figure 7:**
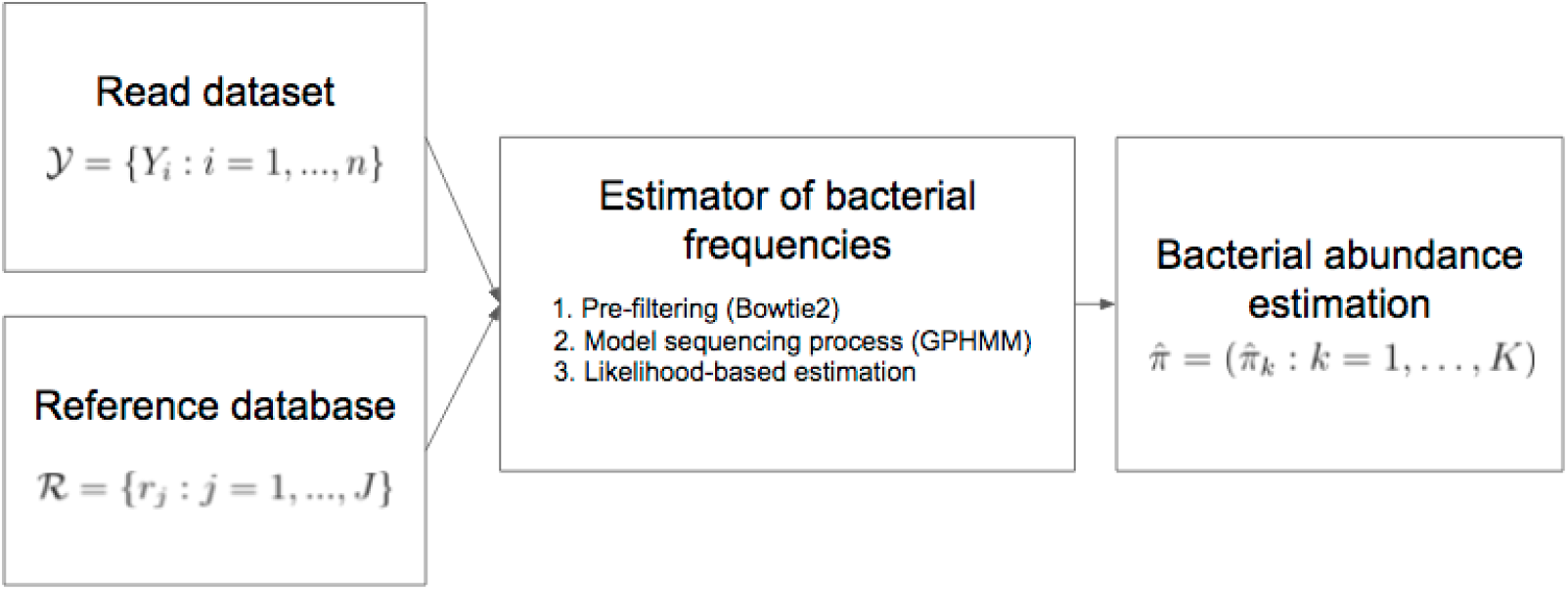
Bacterial composition estimation procedure. The read dataset 𝒚 and the reference database 𝓡 are used to estimate the bacterial composition of the population 𝒚 was derived from.

**Figure 8:**
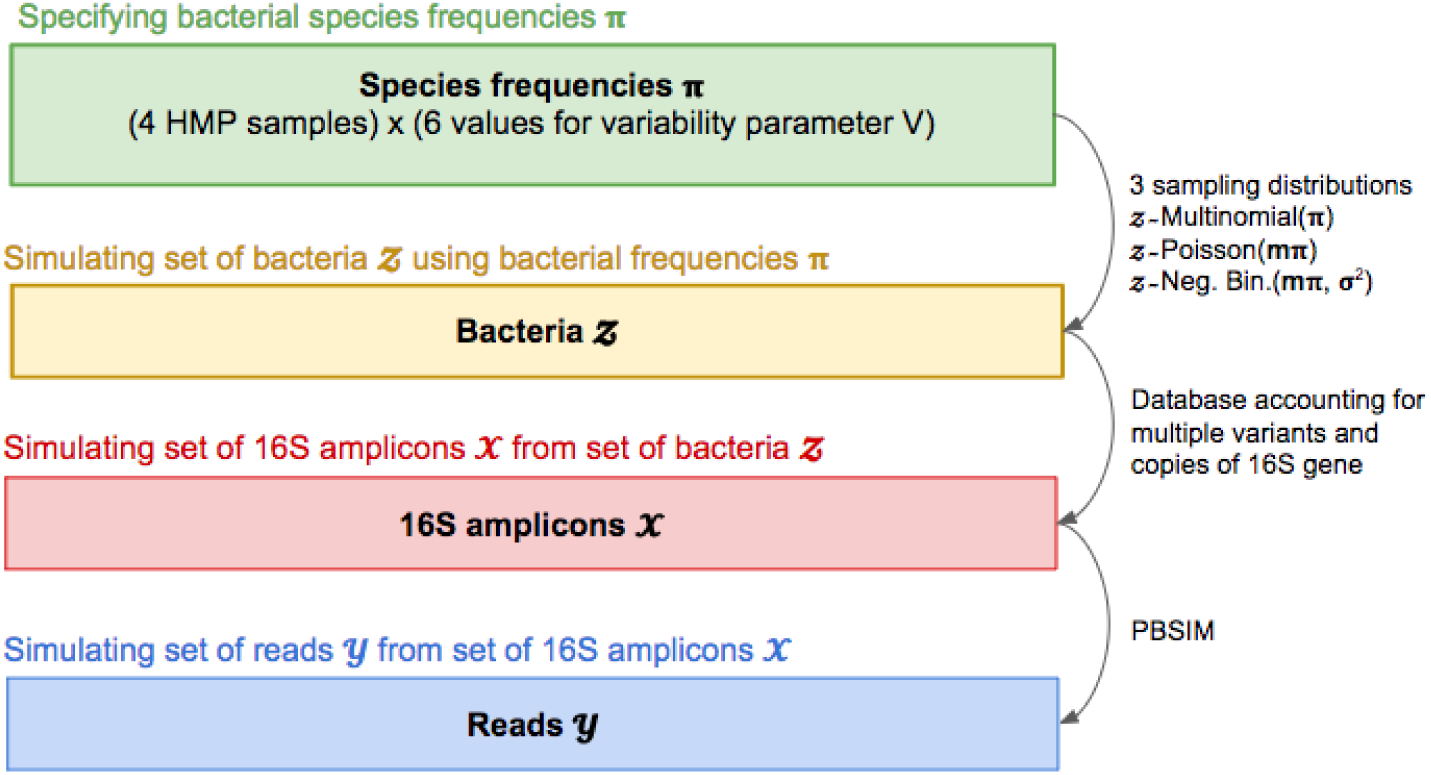
Simulation process.

## 2 Data

### 2.1 Sequencing Reads

#### 2.1.1 Simulated dataset

The goal is to simulate realistic datasets of 16S rRNA reads. In this section, we focus primarily on specifying bacterial frequencies π = (π_*k*_ : *k* = 1,…,*K*); given π, we then use the software package `PBSIM` [67] to generate read datasets 𝒚. The simulation process is summarized in Figure 8.

Different levels of richness (i.e., number of distinct species) and evenness (i.e., uniformity of species frequencies) can be observed in microbial communities. Reduced richness and/or imbalances in the gut microbiome have been associated with a variety of diseases, including obesity [88], inflammatory bowel diseases [53], type

II diabetes [71], and fatty liver disease [6]. As detailed in the remainder of this section, to span the vast range of possible bacterial compositions that have been reported in the literature, we use gut and vaginal samples from the Human Microbiome Project (HMP) [45] to generate bacterial frequencies π. Specifically, to reflect different levels of richness and evenness, we start from four HMP microbial communities (two gut and two vaginal) and, for each such community, simulate bacterial frequencies using six different variability parameters. Additionally, to test the robustness of our method, bacterial communities *Ƶ* are simulated using three different sampling distributions: the multinomial used in our data generation model and two other distributions, a Poisson distribution and a negative binomial distribution allowing over-dispersion. Overall, this yielded 72 simulated read datasets 𝒚, corresponding to four microbial communities (two gut and two vaginal), six variability parameters, and three sampling distributions.

On average, about *n* = 3, 200 reads from *m* = 1,000 bacteria were simulated for each of the 72 datasets. Figure 9 displays an overview of the 24 sets of generative bacterial frequencies π = (π_*k*_ : *k* = 1,…, *K*) for the 72 simulated microbial communities: Number of distinct species Σ_*k*_ 𝟙(π_*k*_ > 0), Shannon diversity —Σ_*k*_ π_*k*_ log π_*k*_ summarizing richness and evenness, and minimum, median, and maximum of the bacterial frequencies π_*k*_. Figure 11 represents the distributions of read lengths and quality scores for each of the 72 simulated datasets. Figure 12 displays the bacterial frequencies π_*k*_ for simulated community *Gut 2* for the two most extreme variability parameters (*V* = 0.1 and *V* = 10, 000).

##### Specifying bacterial species frequencies π

The first step of the simulation is concerned with obtaining the bacterial species frequencies π used to simulate bacterial communities *Ƶ*. The results of an analysis performed as part of the Human Microbiome Project [45] [44] were downloaded from the HMP website (http://hmpdacc.org/, dataset `hmp1.v13.hq`). In this analysis, 16S variable regions 1–3 of about 4, 000 bacterial samples from five different body sites (oral, airways, skin, gut, vagina) were sequenced using the Roche-454 FLX Titanium platform. The software package `Mothur` was used to classify the sequencing reads at the genus level. See details of the analysis in [81]. From these 4,000 samples, we randomly selected four samples, two from the gut body site and two from the vaginal body site, yielding four bacterial communities: *Gut 1, Gut 2, Vaginal 1,* and *Vaginal 2.* Then, for each community, we computed the proportion of reads assigned to each genus using files `hmp1.v13.hq.phylotype.counts` and `hmp1.v13.hq.phylotype.lookup` downloaded from the HMP website. The bacterial genera frequencies for each simulated community were set to be equal to the HMP genera frequencies.

However, as we want to evaluate the ability of our method to estimate bacterial frequencies at the species/sub-species level and the HMP dataset only provides information at the genus level, we use our reference database 𝓡 to identify species and their associated 16S sequences within each genus. Specifically, for each genus, the number of species κ included in the simulation is the number of distinct species found in 𝓡 for this genus. For each species, a reference sequence is then randomly sampled from 𝓡 and its frequency (for simulation purposes) is sampled from a normal distribution with mean 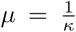 and variance *μV*, where *V* ∈ {0.1,1,10,100,1, 000,10,000}. A small variability parameter *V* yields a balanced microbial community, whereas a high variability parameter corresponds to a less diverse community. Sampled species frequencies greater than 1 or smaller than 0 are set to 0, resulting in a less rich microbial community, especially when the variability parameter is large. Finally, for each genus, species frequencies are scaled to sum to the HMP genus frequency.

This sampling process yields frequencies π = (π_*k*_ : *k* = 1,…, *K*) for each of K bacterial species. For example, for the simulation scenario of the left panel of Figure 12, the genus *Alistides*, with frequency of 0.09 in the HMP dataset, has six distinct species, each with a single reference sequence, according to the reference database 𝓡. This genus is represented in the simulation with six unique sequences, with different frequencies (0.0162, 0.0141, 0.0164, 0.0187, 0.0136, 0.011).

##### Simulating set of bacteria Ƶ using bacterial frequencies π

Following Step 1 of our data generation model (see Figure 4), a set of *m* = 1, 000 bacteria *Ƶ* = {*Z_1_* : *Z_»_* ∈ 𝓑, *i* = 1,…,*m*} is simulated from the Multinomial(*m*, π) distribution. To test the robustness of our method, we additionally used two other distributions to simulate *Ƶ*: A Poisson distribution, where, for each bacterium *b_k_*, *Z_k_* is sampled from Poisson(*mπ_k_*), and a negative binomial distribution, where *Z*_*k*_ is sampled from Negative Binomial(*μ* = *mπ*_*k*_, *σ*^2^ = *μ* + *ϕμ*^2^) with *ϕ* =1/2 (here, *μ* is the mean, *σ*^2^ the variance, and *ϕ* the dispersion parameter, corresponding to the inverse of the `size` argument of the R function `rnbinom`).

##### Simulating set of 16S amplicons 𝓧 from set of bacteria *Ƶ*

For Step 2 of our data generation model (see Figure 5), DNA extraction and PCR efficiencies are set to one (i.e., *e*(*b*_*k*_) = 1 ∀*k* ∈ {1,…,*K*} and *e*(*r_j_*) = 1 ∀*j* ∈ {1,…, *J*}) and the number of PCR cycles is set to *c* =1.

Ideally, our referene database 𝓡 would provide all possible variants and copies of a 16S rRNA gene, i.e., the number *C*_*kj*_ of copies of variant *r_j_* for bacterium *b*_*k*_ would be known ∀*k* ∈ {1,…, *K*} and ∀*k* ∈ {1,…, *J*}). Then, an amplicon dataset 𝓧 could be simulated using the sequences of each variant and the expected number of amplicons for variant with reference sequence *r_j_* from bacterium *b*_*k*_ would be *mπ_k_C_*kj*_*.

However, not all variants of a 16S gene are usually available in our database 𝓡 and *C*_*kj*_ is unknown. Still, to account for the possible multiple variants and copies of a 16S rRNA gene, we estimated the total number of loci at which a 16S gene could be found in the genome of each bacterium 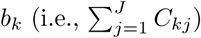 using database rrnDB [85]. Bacteria for which the number of 16S genes is not available at the species level are assigned the number of 16S genes of their lowest taxonomic rank for which the number of 16S genes is available. If bacteria at this taxonomic level have different numbers of 16S genes, the median is used as an approximation for 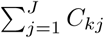. Then, for each bacterium *b*_*k*_, one variant of its 16S gene with reference sequence *r_j_* is randomly sampled from our reference database 𝓡. The number of amplicons for bacterium *b*_*k*_ is therefore expected to be 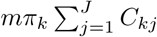.

##### Simulating set of reads 𝒚 from set of 16S amplicons 𝓧

A simulated read dataset 𝒚 is obtained by subjecting each amplicon in the simulated dataset 𝓧 to a sequencing error process. Following the PacBio CCS error model, deletions, insertions, and mismatches are simulated in the amplicon sequences using the `PBSIM` software [67].

Specifically, errors are introduced independently at each position of a sequence according to the following deletion, insertion, and mismatch probabilities, *P_D_*, *P_I_*, and *P_Mis_*, respectively. The deletion probability *P_D_* is assumed constant at each position of a simulated read and defined as

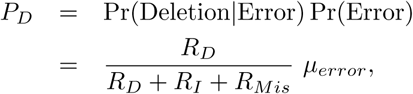

where *μ*_*error*_ is a user-supplied error probability for the read set and *R_D_*, *R_I_*, and *R_Mis_* are, respectively, user-supplied ratios of deletions, insertions, and mismatches. The insertion and mismatch probabilities are computed for each position of a simulated read from the quality score *Q* of the nucleotide at that position:

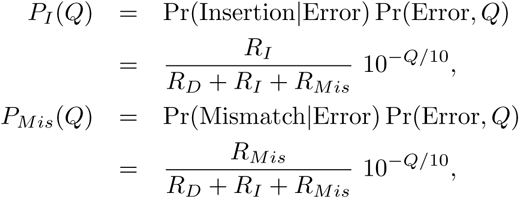

where Pr(Error, *Q*) = 10^−*Q*/10^ is the error probability of nucleotides with quality score *Q* and *Q* is sampled from an in-house FASTQ file containing previously sequenced PacBio CCS reads for 16S genes. Moreover, mismatches are simulated by using a uniform distribution over the four nucleotides {*A, C, G, T*}, while half of inserted nucleotides are chosen to be the same as their following nucleotide and the other half selected from a uniform distribution over {*A, C, G, T*}. For our simulation, default parameters of `PBSIM` are used, that is, *μ_error_* = 0.02, *R_D_* = 0.73, *R_I_* = 0.21, and *R_Mis_* = 0.06.

#### 2.1.2 Microbial mock community

We estimate the performance of our methods on a mock community composed of an even distribution of genomic DNA from 21 bacterial strains: Acinetobacter baumannii ATCC 17978, Actinomyces odontolyticus ATCC 17982, Bacillus cereus ATCC 10987, Bacteroides vulgatus ATCC 8482, Clostridium beijerinckiiATCC 51743, Deinococcus radiodurans ATCC 13939, Enterococcus faecalis ATCC 47077, Escherichia coli ATCC 70096, Helicobacter pylori ATCC 700392, Lactobacillus gasseri ATCC33323, Listeria monocytogenes ATCC BAA-679, Neisseria meningitidis ATCC BAA-335, Porphyromonas gingivalis ATCC 33277, Propionibacterium acnes DSM 16379, Pseudomonasaeruginosa ATCC 47085, Rhodobacter sphaeroides ATCC 17023, Staphylococcus aureusATCC BAA-1718, Staphylococcus epidermidis ATCC 12228, Streptococcus agalactiae ATCCBAA-611, Streptococcus mutans ATCC 700610, and Streptococcus pneumoniae ATCC BAA-334. The data used here are a subset of the data used in [79] (v3.1, HM-278D) and were downloaded from the Sequence Read Archive at NCBI under accession SRP051686 associated with BioProject P*R_J_*NA271568.

Library generation, sequencing, and pre-processing to generate the downloaded reads are described in [79]. Briefly, we used reads sequenced from the V1-V9 variable region (full length 16S rRNA gene) sequenced by Pacific Biosciences using the P6-C4 chemistry on a PacBio RS II SMRT DNA Sequencing System with MagBead loading. We filtered out reads with length smaller than 1,300 and greater than 1,600 bp resulting in a dataset with 66,450 reads. See the distribution of the length of the reads in Figure 13. For this mock community, the individual DNA extracts were mixed based on the genome size and the number of different loci where 16S rDNA genes are in each genome to have equal-molar 16S rDNA copies for each species [41]. So, as opposed to section 1.5.2, we did not account for the multiple variants and copies of the 16S gene for the different strains.

**Figure 9:**
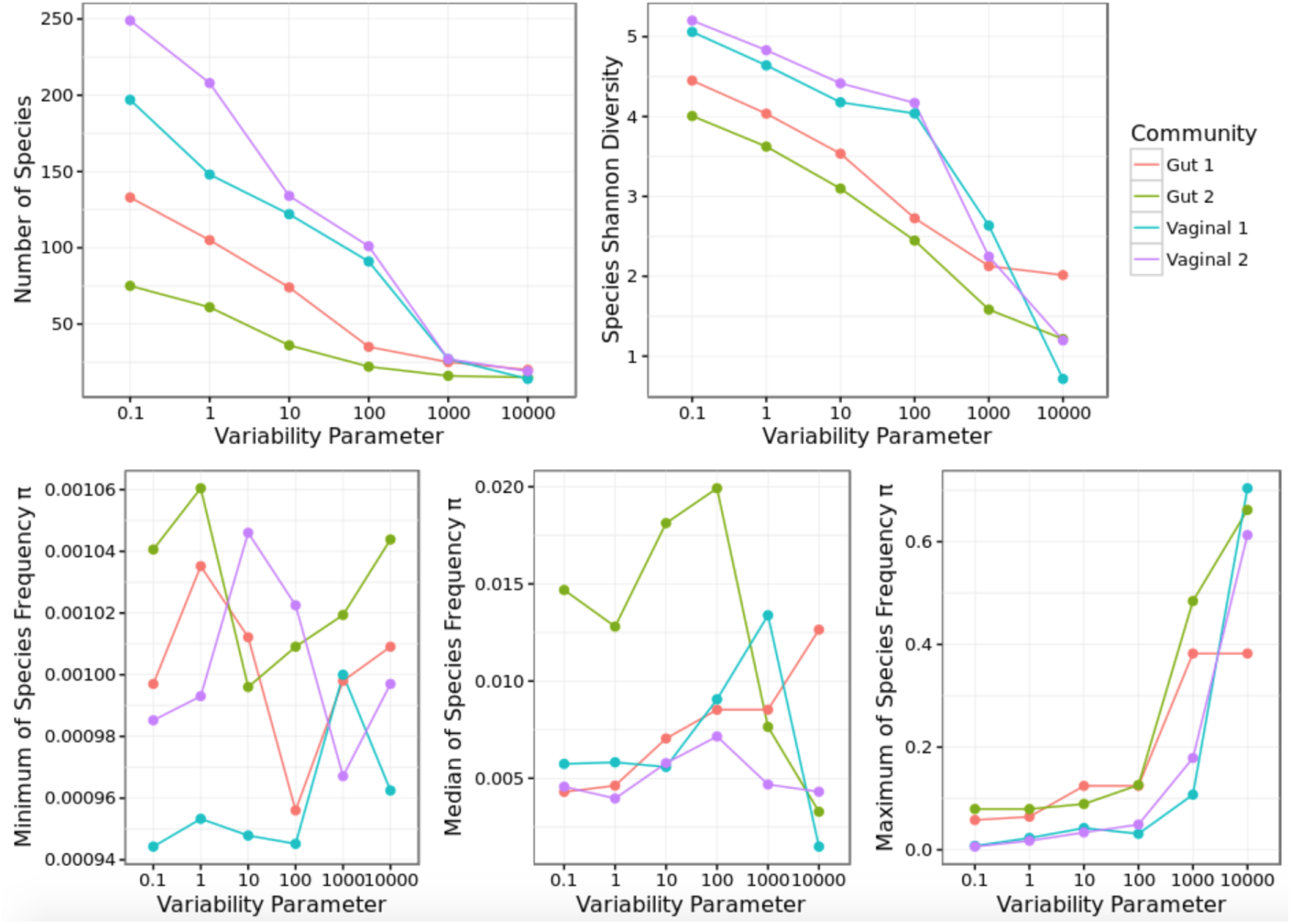
Overview of bacterial frequencies for the simulated datasets. Read datasets were simulated using 24 different sets of bacterial frequencies π = (π_*k*_ : *k* = 1,…,*K*) (at the species level). The frequencies π correspond to four HMP communities (two gut and two vaginal) and six different variability parameters (*V* ∈ {0.1,1,10,100,1, 000,10, 000}) for the sampling of species from each of these communities. Top-left panel: Number of distinct species Σ_*k*_ 𝟙(π_*k*_ > 0). Top-right panel: Shannon diversity of bacterial frequencies — Σ_*k*_ π_*k*_ log π_*k*_, providing a measure of richness and evenness of a microbial community. Bottom panels, from left to right: Minimum, median, and maximum of the bacterial frequencies π_*k*_. While minimum and median bacterial frequencies tend to be similar across simulated datasets, the maximum bacterial frequency is higher when the variability parameter is larger, introducing imbalance in the microbial community. Note that the y-axis scales are different for the graphs in the bottom panels.

**Figure 10:**
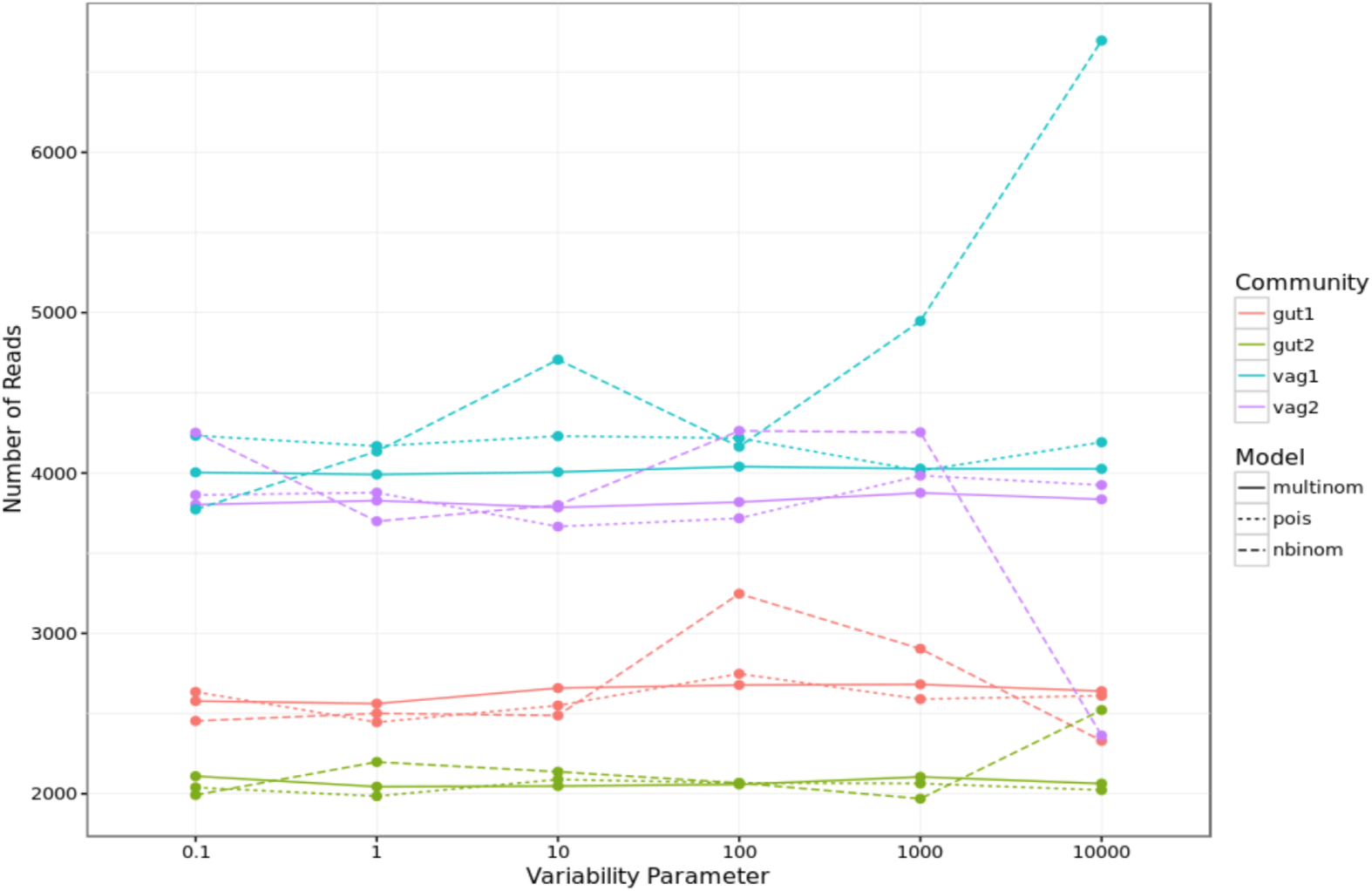
Number of reads in simulated datasets. The total number of bacteria in a simulated community *Ƶ* is *m* = 1, 000. If there was only one 16S gene per bacterium, the number of reads would be equal to the number of bacteria. In practice, however, there are multiples variants and copies of a 16S gene for a given bacterium, so the number of reads *n* is greater than *m*. The number of reads is bigger for the vaginal simulations than for the gut simulations because there are more 16S genes in the genome of the species picked for the vaginal simulations. Not surprisingly, the number of reads is more variable for the negative binomial model as the number of bacteria simulated for each species *b*_*k*_ is more variable. It is especially true when π_*k*_ is big. For example, for *Vaginal 1, V* = 10,000, and model negative binomial, the number of reads is big. In this simulated dataset, the two most abundant strains are from species *Lactobacillus vaccinostercus* (π = 0.70) and *Lactobacillus sanfranciscensis* (π = 0.27). Therefore, the means of the negative binomial were respectively 0.7 * 1000 = 700 and 0.27 * 1000 = 270, but because of the over-dispersion of the negative binomial, the number of bacteria simulated were respectively 991 and 644. Both of the strains have four 16S genes in their genome, explaining the about 6, 500 reads in this dataset.

**Figure 11:**
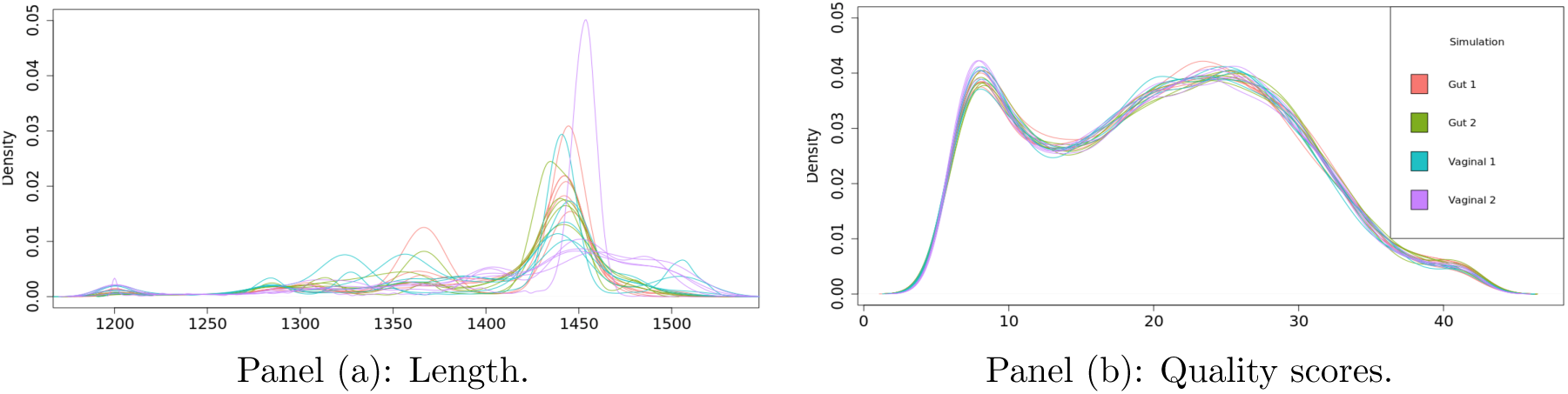
Panel (a): Distribution of read lengths for each of the 72 simulated datasets. Panel (b): Distribution of Phred read quality scores for each of the 72 simulated datasets (extracted from FASTQ file generated by software PBSIM).

**Figure 12:**
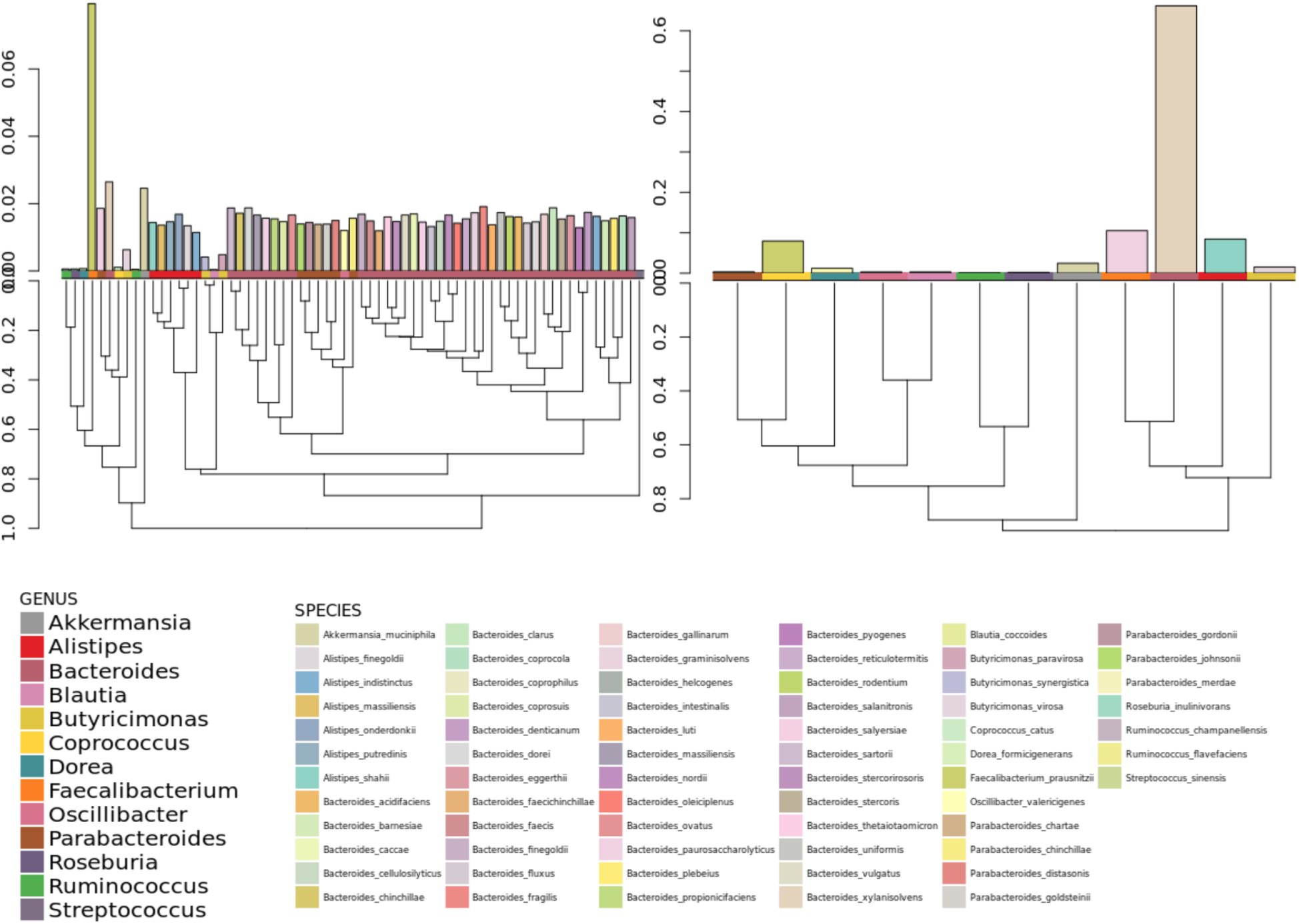
Phylogenetic tree and frequencies of bacterial species π in two simulation scenarios. Barplots in the top-left and top-right panels represent bacterial frequencies *πk* for community *Gut 2* with the two most extreme variability parameters, respectively, *V* = 0.1 and *V* = 10, 000. The smallest variability parameter yields a rich and balanced microbial community, whereas the greatest variability parameter results in a less rich and more imbalanced community. The phylogenetic trees in the bottom panels represent the similarity between the bacterial species, where the distance between two species is the Levenshtein distance between their consensus 16S gene sequences. The horizontal colored bars above the phylogenetic trees indicate the genus of each species.

**Figure 13:**
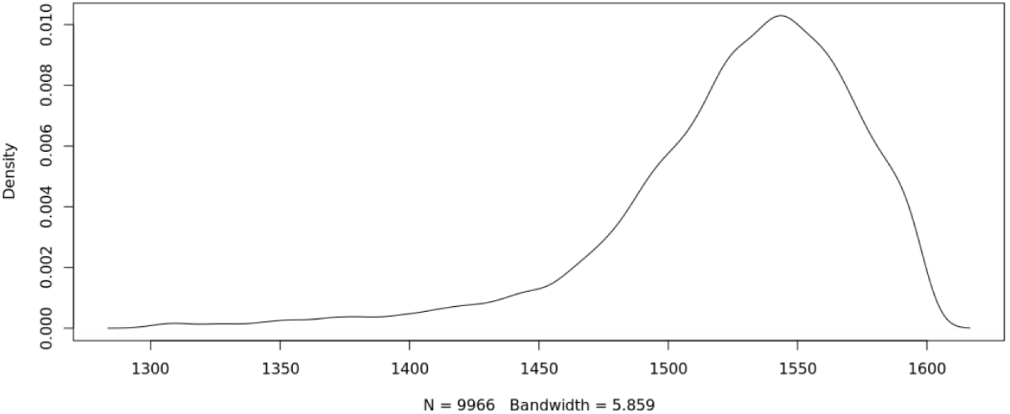
Distribution of the length of the reads in the filtered dataset for the mock community.

### 2.2 16S Database

It is essential to use a reference database with high confident taxonomic classifications. In our method, each read is assigned the same taxonomic level as its most similar sequence in a reference database. Then, if the database has annotation errors, a prediction could be wrong not because of incorrect assignment of the read to a reference sequence, but because of incorrect annotations of the reference sequences in the database. Among the different available databases (e.g., SILVA, Greengenes), we decided to use the full RDP database because it is exhaustive, i.e., it contains most of the sequences found in the other databases.

First, we downloaded the full RDP database, version 11.4, with 3,070,243 16S gene sequences. After filtering out sequences with length outside of the range 1,300bp to 1,600bp, sequences with ambiguous bases (i.e., nucleotides that are not A,C,G,T), sequences with identical DNA sequences (Bowtie2 does not allow duplicated sequences), and sequences with entirely redundant annotation (i.e., annotation where the only difference is the RDP identifier), 1,362,820 16S gene sequences remained. To allow the possibility that novel reference sequences could enter the database, we used 32-byte md5 hash values as the sequences identifiers. Then, when we want to add a sequence to the database, we compute its md5-hash value, determine if it exists in our dataset, if so, simply add the annotation to the corresponding reference sequence, otherwise construct a new entry in our database.

The last step is to add lineage when it is not specified in the filtered RDP database. The taxonomies in the full RDP database were predicted by the RDP Classifier which has a high rate of over-classification errors (i.e., novel taxa are incorrectly predicted to have known names) on full-length sequences [29]. To get a database with only well-annotated reference sequences, we filtered out the filtered RDP database to keep only reference sequences with taxonomic annotations at the species level, often added manually to the database, thus highly accurate annotations. It resulted in a species level annotated RDP database with 81,088 16S gene sequences. We then used this species level annotated database to aligned the entire set of sequences in the filtered RDP database. If alignments surpass 99% of similarity, we re-annotated the reference sequence with the best matching reference’s annotation.

**Figure 14:**
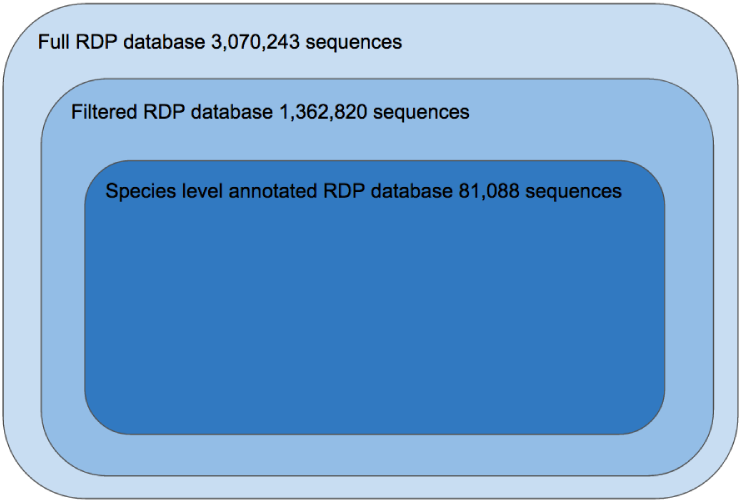
16S databases. The full RDP database, version 11.4, with 3,070,243 16S gene sequences was downloaded and filtered out to get a filtered RDP database with 1,362,820 16S gene sequences. To create a database with only well-annotated reference sequences, we filtered out the filtered RDP database to keep only reference sequences with taxonomic annotations at the species level, resulting in a reduced RDP database with 81,088 16S gene sequences. Both the filtered and species level annotated RDP databases were used to classify the reads.

Both filtered and species level annotated databases were used to classify reads. See Figure 14. The former is used to increase the sensitivity (i.e. decrease the number of false negatives, that is make sure we do not miss strains) at the risk of assigning reads to sequences with incorrect annotations. The later is used to increase the specificity (i.e. decrease the number of false positives, that is make sure we do not incorrectly call a strain present while it is absent) at the risk of missing strains that have been filtered out from the database. In this report, we only show results when the species level annotated RDP database is used.

## 3 Methods

### 3.1 Training and Validation of Bacterial Composition Estimation Procedure

In order to train our bacterial composition estimation procedure, i.e., select optimal Bowtie2 tuning parameters and estimate the parameters of the HMM, and evaluate its overall accuracy, we dispose of an annotated dataset 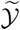 for which we know, for each read *Y_i_*, its corresponding true 16S gene sequence 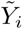,

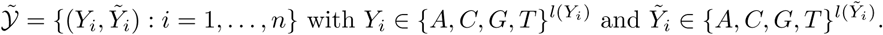

We divide the annotated learning dataset 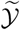 at random into two datasets:

- a training set 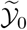, containing *n*_0_ reads (*n*_0_ = 5,000), used only to select optimal Bowtie2 tuning parameters and estimate the parameters of the HMM,
- a validation set 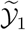, containing *n*_1_ = *n* — *n*_0_ reads (*n*_1_ = 5, 000), used only to assess the overall accuracy of the procedure trained using 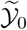.

By construction, reads in 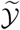 are generated from 16S gene sequences present in our database 𝓡, that is, 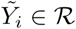 for each *i* = 1,…,*n*, so that one can identify for each pair 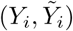 the bacteria of origin 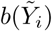.

**Figure 15:**
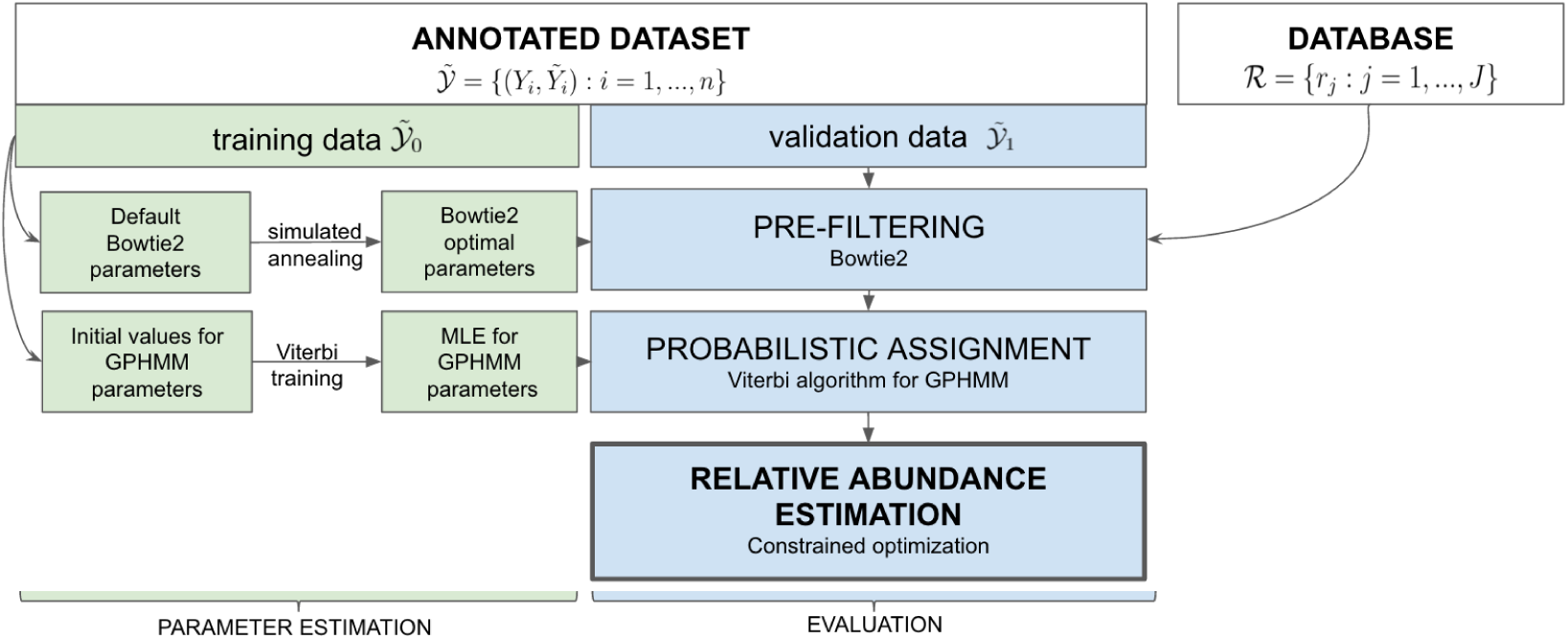
Pipeline for training and validation of bacterial composition estimation procedure.

### 3.2 Pre-filtering

#### 3.2.1 Bowtie2

To estimate the bacterial composition of a microbiomic sample, we need to compute the likelihood Pr(𝒚; π), meaning that we need to compute Pr(*Y_i_* = *y_i_*|*X_i_* = *r_j_*) for each read *y_i_* in 𝒚 and each reference sequence *r_j_* in 𝓡 (Section 1.5.3). The number of reference sequences in 𝓡 being on the order of 1.4 million and the number of reads in 𝒚 on the order of the thousands, such a computation is intractable. To reduce the number of computations, we use the alignment tool Bowtie2 to eliminate reference sequences with small probabilities of having generated the reads in 𝒚.

Bowtie2 first indexes the set of reference sequences in our database 𝓡 using a scheme based on the Burrows-Wheeler transform [12] and FM-index [31], which allows compression of the database while still permitting fast substring queries. Then, each read in 𝒚 is divided into substrings — called seeds — and only reference sequences matching almost perfectly the seed substrings are kept. Seed substrings are then extended to the entire length of the read and a score is given to each alignment between the read and the selected reference sequences. This alignment score (AS) quantifies how similar a read is to a reference sequence it is aligned to; the higher the score, the higher the chance that the read could have been generated from the reference sequence. The score is calculated by adding a “bonus” for each match and subtracting a “penalty” for each difference (mismatch, insertion, or deletion) between the read and the reference sequence. Only reference sequences with an AS higher than a minimum alignment score (min-score) are selected. That is, for each read Y, the set of selected reference sequences is given by the mapping

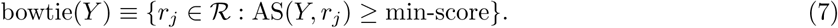

Unlike other index-based alignment tools, such as, Burrows-Wheeler Aligner (BWA) [59], BWA’s Smith-Waterman alignment (BWA-SW) [58], and short oligonucleotide alignment program 2 (SOAP2) [60], Bowtie2 has a rich set of tuning parameters, that have a large influence on the selection of the candidate reference sequences and are highly dependent on read length. As reads in our dataset are long reads, spanning the entire length of the 16S rRNA genes (⋍ 1,500 bp), and Bowtie2 default parameters are tuned for shorter reads, it is essential to select appropriate parameters for our setting:

- L, the length of the seed substrings,
- i, the interval between seed substrings,
- R, the maximum number of times Bowtie2 re-seeds reads with repetitive seeds,
- D, the number of consecutive seed extension attempts that can fail before Bowtie2 moves on,
- k, the maximum number of candidate reference sequences that can be selected during the seed search,
- min-score, the minimum alignment score needed for an alignment to be good enough to be selected,
- the parameters configuring the alignment scores: match bonus (ma), mismatch penalty (mp), deletion penalty (rdg), and insertion penalty (rfg).

For example, if for read *Y* the maximum number of candidates k allowed during the seed search is set low and there are many more candidate reference sequences with an AS higher than min-score, then Bowtie2 can select k candidate reference sequences matching the seeds in *Y* before finding the correct reference sequence. See the Bowtie2 manual for more details on tuning parameters.

Our goal is to find the optimal set of Bowtie2 tuning parameters, *θ* = {L, i, R, D, k, min-score, ma, mp, rdg, rfg}, so that the true sequence for read *Y* is among the candidate reference sequences found by Bowtie2. For our training set 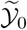 (Section 3.1), we know that each observed noisy read *Y_i_* was generated from a true sequence 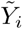. Then, we can define an objective function *L* to be maximized over the set of Bowtie2 tuning parameters *θ*:

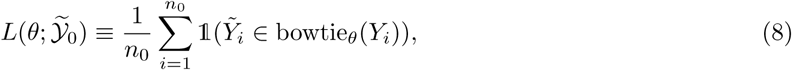

where bowtie_*θ*_ is the function mapping each read *Y_i_* to the candidate reference sequences selected by Bowtie2 with tuning parameters *θ*.

#### 3.2.2 Selecting optimal Bowtie2 parameters: simulated annealing

Simulated annealing (SA) [51] is used to maximize the objective function 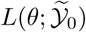. SA is a powerful technique for approximating global optimization in a large search space and is known to perform well when the search space is discrete – which is the case here. For each individual parameter in *θ*, a search domain with lower and upper bounds is specified and random initial estimates of the parameters are chosen within the search domains. At each iteration *i*, the SA heuristic considers some neighboring parameter *θ*_*i*+1_ of the current parameter *θ_i_*. The probability of making the transition from the current *θ_i_* to the candidate new parameter *θ*_*i*+1_ (acceptance probability) is then calculated. The system moves ultimately to sets of parameters with higher objective function, the process being repeated until a stopping criterion is reached.

More precisely, at each step *i*, reads in our training set 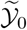 are aligned to our database 𝓡 using Bowtie2 with parameter *θ_i_* and an acceptance probability is calculated as follows

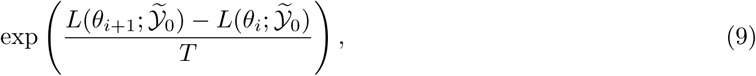

with *T* a global time-varying parameter called temperature. If the acceptance probability is larger than a value sampled uniformly between zero and one, then *θ* is set to the new value *θ*_*i*+1_, otherwise it stays as *θ_i_*. The search ends when an acceptable solution is found, that is, the objective function is higher than a threshold, or the maximum number of iterations is reached. The pseudocode corresponding to the description above is provided in Algorithm 1.

##### Algorithm 1: Simulated annealing

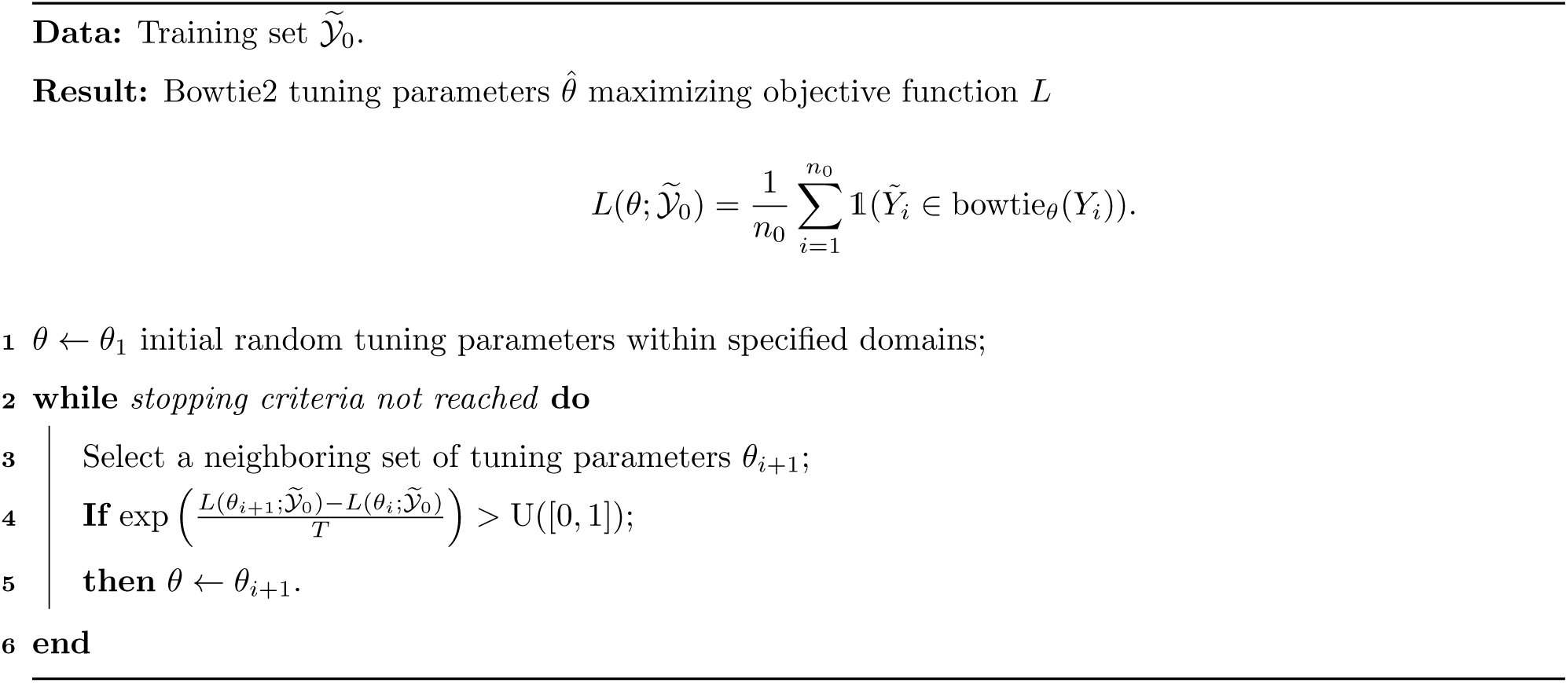

### 3.3 Modeling the Sequencing Process

We want to compute the probability Pr(*Y*|*X*) that an observed noisy read *Y* in our dataset 𝒚 was generated from an unobserved 16S gene sequence *X*. To do so, we need to consider the set 𝒬 of all possible alignments between *X* and *Y*, i.e., the ways *X* can be construed to have generated *Y*,

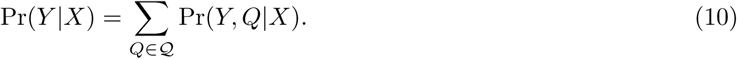

Specifically, three types of errors can occur in the sequencing of a given gene *X*:

- mismatches, which substitute a nucleotide in *X* with another nucleotide in *Y*,
- insertions, which add nucleotides in *Y* compared to *X*,
- deletions, which delete nucleotides in *Y* compared to *X*.

An alignment *Q* between two sequences *X* and *Y* therefore consists of a sequence of matches, mismatches, insertions, and deletions.

Using Equation (10), the probability Pr(*Y, Q*|*X*) would have to be computed for each alignment *Q* in 𝒬. The number of possible alignments between *X* and *Y* being the product of the lengths of *X* and *Y* (*l*(*X*)*l*(*Y*) ⋍ 1,500^2^ = 2.25 × 10^6^ for our problem), the computation is prohibitive. It has been shown that computing the probability of the optimal alignment between *X* and *Y*, instead of summing over the probabilities of all possible alignments, is substantially more computationally efficient, with no impact on accuracy [ref, PacBio suppl. paper, other ref]. Thus, we compute

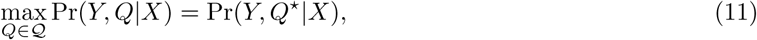

where *Q*^*^ arg max_*Q∈𝒬*_ Pr(*Y, Q*|*X*) is the most probable alignment between *X* and *Y*.

As detailed next, we model the sequencing process relating *Y* to *X* using a generalized pair hidden Markov model and use the Viterbi algorithm to infer the optimal alignment *Q*^*^. Although the true sequence *X* is unobserved, in what follows, *X* is treated as observed while the alignment *Q* is unobserved.

#### 3.3.1 Generalized pair hidden Markov model

Instead of observing a single sequence of random variables, as is usually the case for a hidden Markov model (HMM), here, we have a pair of sequences of random variables, namely, the unaligned nucleotide sequences for a read Y and the putative 16S gene *X* it was sequenced from. We therefore adopt a generalized pair hidden Markov model (GPHMM) to model the sequencing process, where each hidden state emits a pair of sequences. As seen below, our model is a special case of a generalized pair HMM, in the sense that the durations are deterministic given the hidden states.

Our GPHMM is defined as follows, where, for the rest of Section 3.3, we modify the notation adopted above for an observed read Y and the corresponding unobserved 16S gene sequence *X*.

1. **Hidden states and durations.** Let {*X_t_* : *t* = 1,…, *T*} denote the hidden states corresponding to a particular pairwise alignment between two sequences *Y̅*^1^ and *Y̅*^2^, representing, respectively, a 16S gene and its corresponding sequencing read. Three hidden states 𝒮 = {*M, Ins, Del*} are required to represent the sequence of matches/mismatches, insertions, and deletions corresponding to a particular pairwise alignment,

- *M*, the state for matches/mismatches,
- *Ins*, the state for insertions,
- *Del*, the state for deletions. To each hidden state *X_t_* ∈ 𝒮, we associate a pair of also hidden random durations 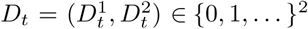, generated according to the conditional distribution *p_i_*(*d*) ≡ Pr(*D_t_* = *d*|*X_t_* = *i*), and denoting the number of nucleotides emitted for the 16S gene sequence and the read, respectively. In our application, we consider a special case where durations are either 0 or 1 and are deterministic given hidden states, 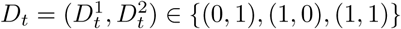 and

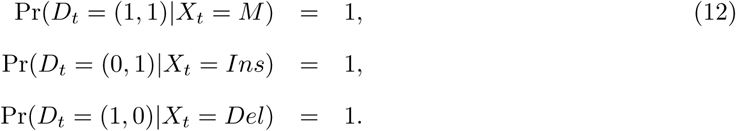 Let 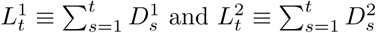 denote the cumulative durations up to time *t*.
2. **Emitted sequences.** Each hidden state *X_t_* emits a pair of sequences 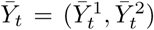, where 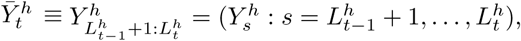*h*= 1,2. In our special case, 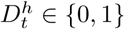 and

- if 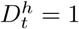, then one nucleotide 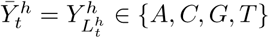 is emitted at location 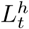,
- if 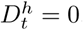, then no nucleotide is emitted and 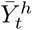 is the empty string ∅. Then, the unobserved emitted aligned pair of sequences is

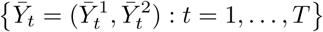

and yields an observed unaligned pair of sequences

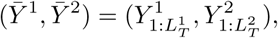

where 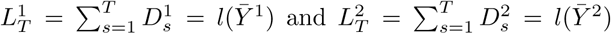 are the lengths of the true 16S gene sequence and the read, respectively. Note that, in this GPHMM, only 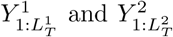 are observed, that is, *X_t_*, *D_t_*, *Y̅_t_*, and *T* are hidden.
3. **State transition probability distribution.** We denote the state transition probability matrix by *A* = (*A_ij_* : *i, j* ∈ 𝒮). To simplify our model, we assume that transitions from state *Ins* to state *Del* and from *Del* to *Ins* are not allowed. We also define the following probabilities

- *γ_I_* the transition probability from state *M* to state *Ins*,
- *γ_D_* the transition probability from state *M* to state *Del*,
- *є_I_* the probability of staying in state *Ins*,
- *є_D_* the probability of staying in state *Del*. The state transition probability matrix *A* can then be written as

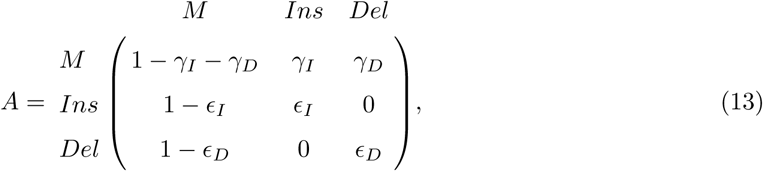

where the other entries of *A* result from noting that *A* is a stochastic matrix, i.e., its rows sum to one. For example, the probability to stay in state *M* is 1 — *γ_I_* — *γ_D_*.
4. **Initial state distribution.** The initial state distribution *a* = (*a* : *i* ∊ 𝒮)is the same as the first row of the transition matrix *A*, that is,

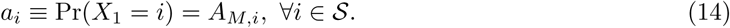
5. **Emission probability distribution.** Emitted sequences *Y̅_t_* are generated according to the conditional joint distribution

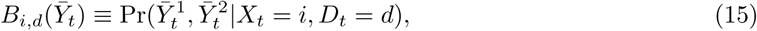

which depends only on the current hidden state *X_t_* = *i* ∈ 𝒮 and pair of durations *D_t_* = *d* ∈ {(0,1), (1, 0), (1,1)} and is represented in the matrix in Figure 16.

For convenience, we denote by λ ≡ (*A, B*) the entire parameter set of our model.

A pair of sequences can be generated as follows from our GPHMM, for a given value of the parameter λ.

**Figure 16:**
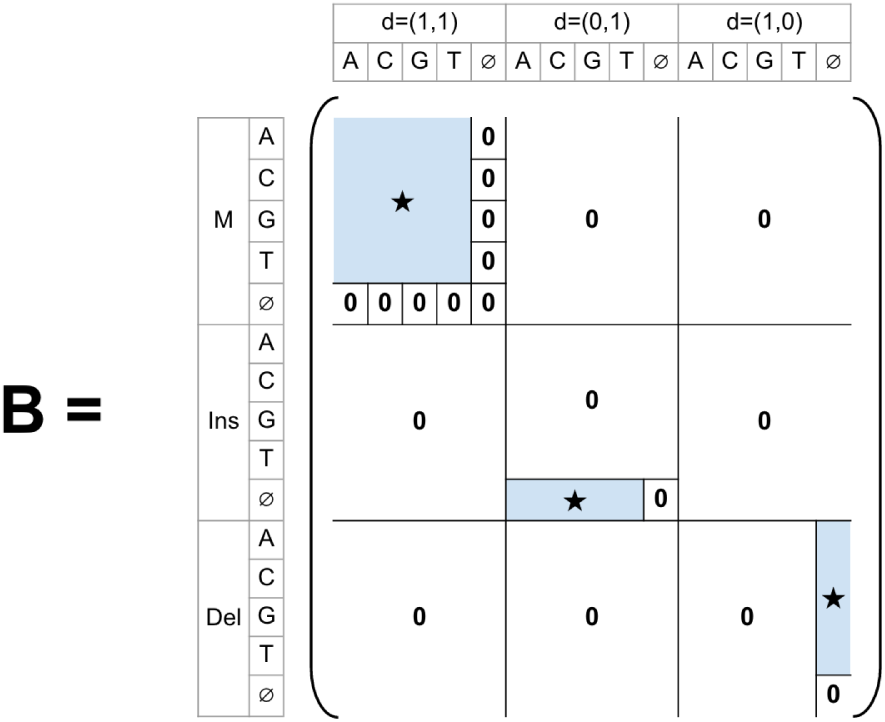
Emission probability matrix. Stars represent non-null entries.

1. Generate the first hidden state *X*_1_ ∈ 𝒮 according to initial state distribution *a* in Equation (14). Given state *X*_1_, generate durations 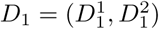 according to Equation (12). Given state *X*_1_ and durations *D*_1_, emit a pair of sequences 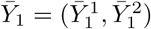 according to the emission distribution *B*. For example, in Figure 17, *X*_1_ = *M*, thus *p_M_*(1,1) = 1 and *D*_1_ = (1,1). The emitted pair of sequences is then chosen according to *B_M_*,(1,1) and is, for instance, 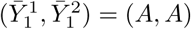.
2. Given *X*_1_, select the next hidden state *X*_2_ ∈ 𝒮 according to the transition probability matrix *A*. For example, in Figure 17, *X*_2_ = *Del*, thus *p_Del_*(1, 0) = 1 and *D*_2_ = (1, 0). The emitted pair of sequences is then chosen according to *B*_*Del*,(1,0)_ and is, for instance, 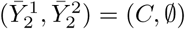.
3. Repeat Step 2 until 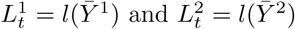.

#### 3.3.2 Most probable alignment: Viterbi algorithm

We want to find the sequence of hidden states *X*_*t*_ that describes best the fact that the read *Y̅*^2^ could have been sequenced from the 16S gene *Y̅*^1^, where our definition of best means maximizing the conditional probability of the sequences of hidden states *X̅* = (*X*_*t*_ : *t* = 1,…, *T*) and pairs of durations *D̅* = (*D*_*t*_ : *t* = 1,…, *T*) given the read and its corresponding true sequence. That is, we want to maximize Pr(*X̅*, *D̅*|*Y̅*^1^, *Y̅*^2^), which is equivalent to maximizing Pr(*X̅*, *D̅*, *Y̅*^1^, *Y̅*^2^), over {*T*, *X*, *D̅*}.

We use the dynamic programming method called the Viterbi algorithm, which involves computing the function *δ*(*i*, (*d*_1_, *d*_2_), (*l*_1_,*l*_2_)), defined as the highest probability of an emitted pair of sequences of lengths (*l*_1_,*l*_2_) and corresponding sequences of hidden states and durations, ending in hidden state *X*_*t*_ = *i* with durations *D_t_* = (*d*_1_, *d*_2_),

**Figure 17:**
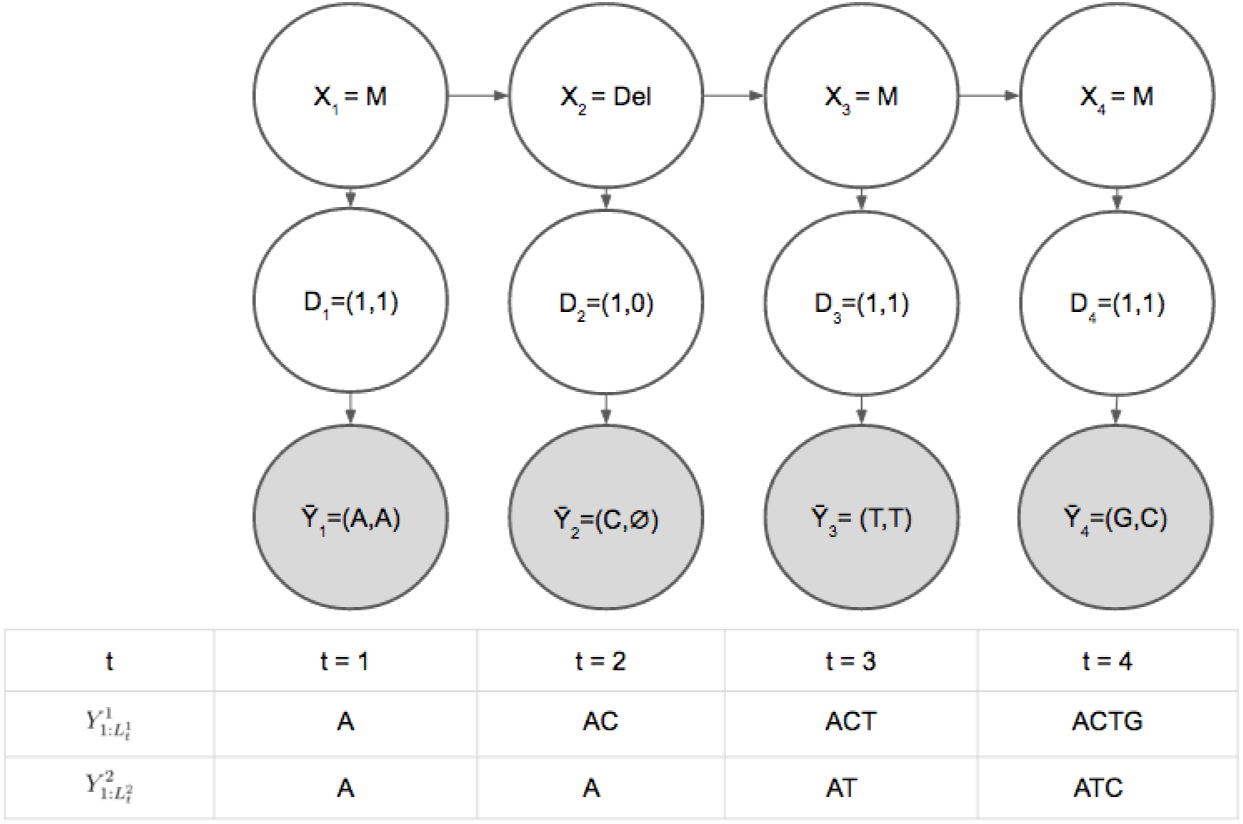
Generalized pair hidden Markov model. Generation of a 16S gene sequence and its corresponding sequencing read using our generalized pair HMM. Upper white circles represent hidden states, intermediate white circles durations, and lower grey circles pairs of nucleotide sequences emitted at each state. The table below the graph represents the unaligned pairs of sequences up to time *t* = 1,…, *T* = 4. Only the 16S gene 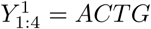 and the read 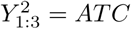 are observed.

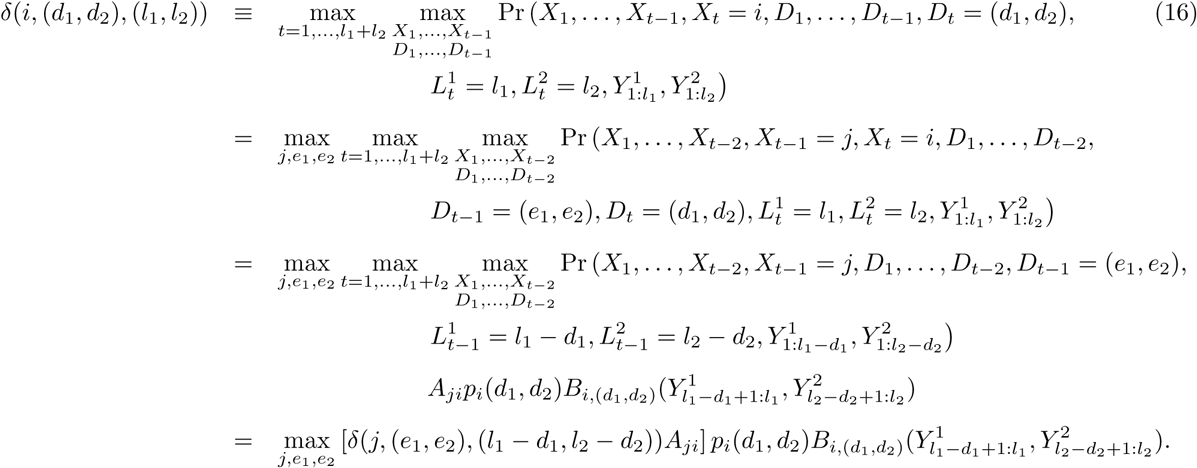

Additionally, to retrieve the optimal sequences of hidden states and durations for the alignment, we need to keep track of the arguments *i* and (*d*_1_,*d*_2_) that maximize *δ*(*i*, (*d*_1_,*d*_2_), (*l*_1_,*l*_2_)) for each pair of emitted sequence lengths (*l*_1_, *l*_2_). To do so, we use a function *ψ*(*i*, (*d*_1_,*d*_2_), (*l*_1_,*l*_2_)). Then, the steps of the Viterbi algorithm are as follows.

**Figure 18:**
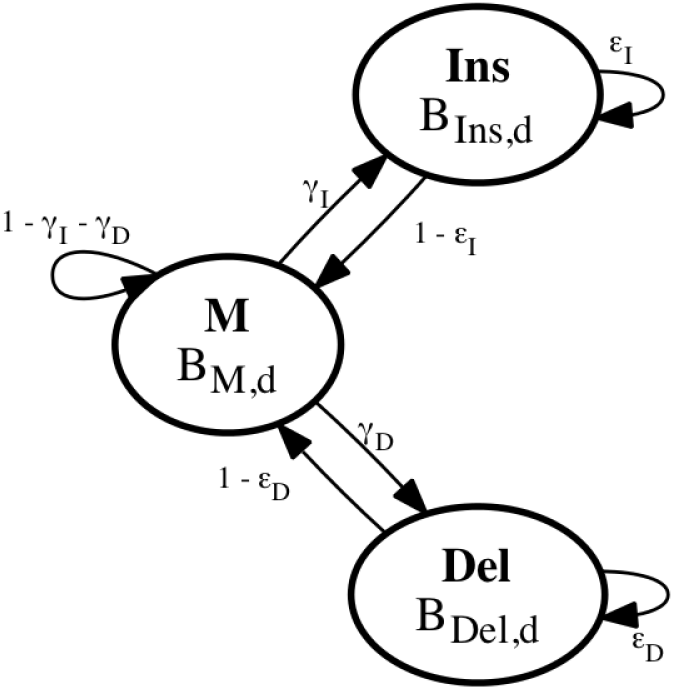
Generalized pair hidden Markov model. Hidden states and transition and emission probability distributions.

1. **Initialization.** Let

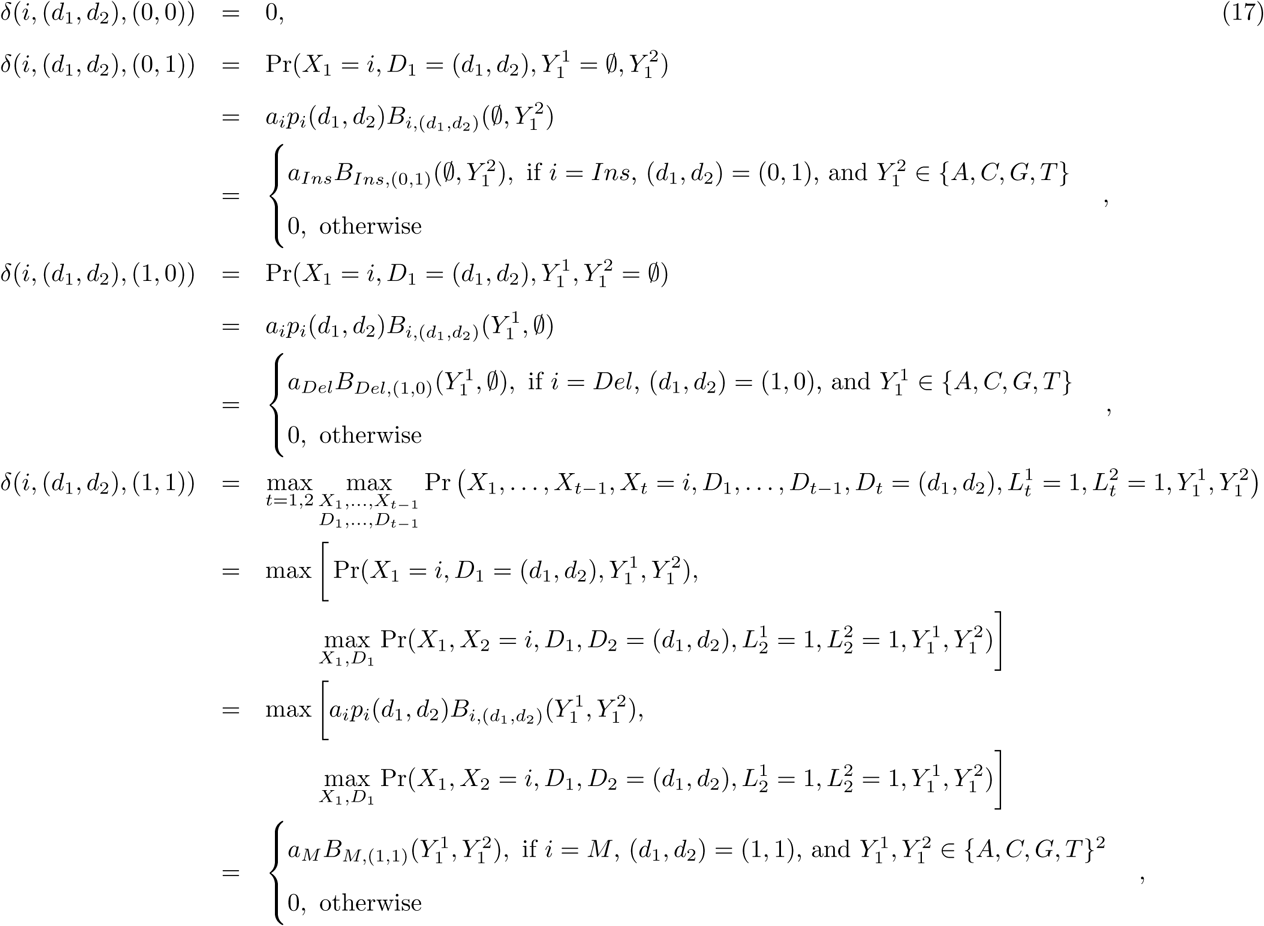

where the above simplification for *δ*(*i*, (*d*_1_,*d*_2_), (1,1)) follows by noting that *A_Ins,Del_* = *A_Del,Ins_* = 0, so that 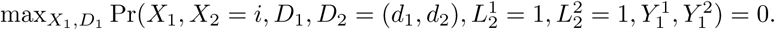 For *i* ∈ 𝒮 and (*d*_1_, *d*_2_) ∊ {(0,1), (1, 0), (1,1)}, also let

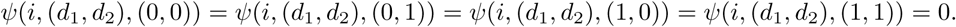
2. **Recursion.**

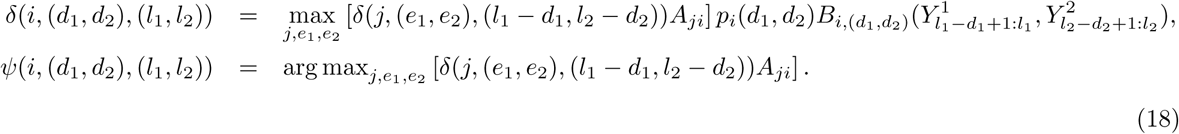
3. **Termination.**

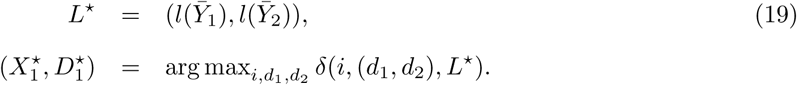
4. **Backtracking.** Set *t* = 1. While *L*^*^ ≠ (0, 0),

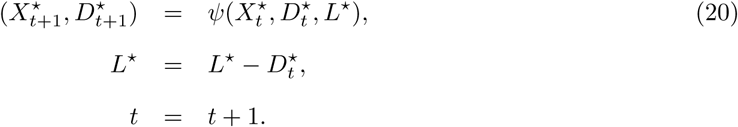 At the end of the loop, *t* = *T* − 1. The sequences of hidden states and durations can then be obtained by reversing the order of the elements of *X*^*^ and *D*^*^.

##### More efficient implementation of the Viterbi algorithm

The emission probability matrix in Figure 16 being sparse, it is possible to reduce the number of iterations in the recursion of Equation (18). Durbin et al. [23] propose a faster implementation of the Viterbi algorithm, where *Begin* and *End* states are added for, respectively, the initialization and termination of a pairwise alignment and where one does not make use of durations *D*_*t*_.

There are no emissions from the *Begin* and *End* states. The transition probabilities from any hidden state in {*M, Ins, Del*} to the *End* state are set to *τ* and the transition probabilities from the *Begin* state to any hidden state in {*M, Ins, Del*} are given by the first row of the transition probability matrix *A* in Equation (13). Transitions from and to the *Begin* and *End* states are designated by dashed lines in Figure 19. For simplicity, it is assumed that an alignment starts in the *M* state. With 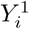 denoting the nucleotide at location *i* in the true sequence *Y̅*_1_ and 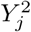 the nucleotide at location *j* in the read *Y̅*_2_, the algorithm is as follows.

1. **Initialization.** For 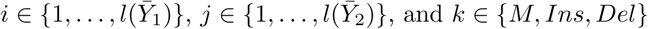, let

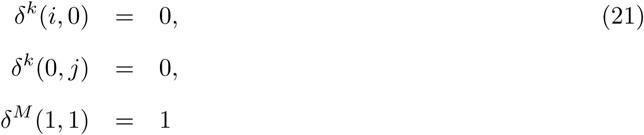

and

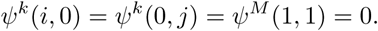
2. **Recursion.** For 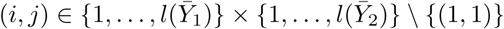,

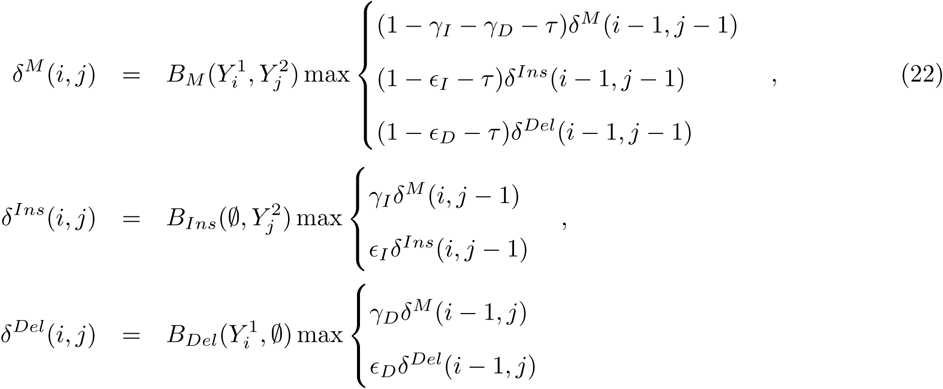

and

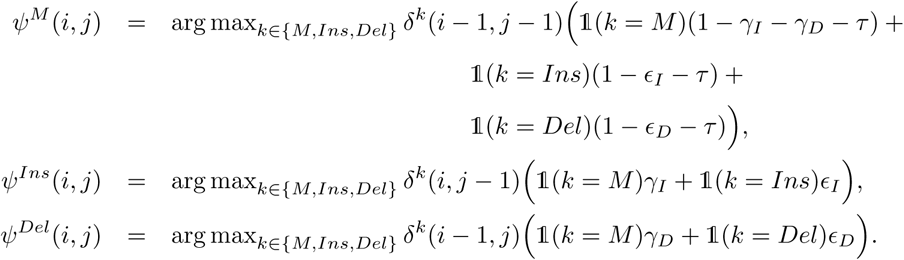
3. **Termination.**

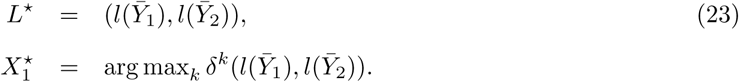
4. **Backtracking.** Set *t* = 1. While *L*^*^ ≠ (0, 0),

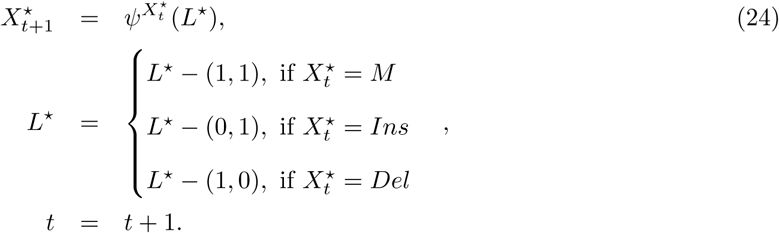 At the end of the loop, *t* = *T* − 1. The sequence of hidden states can then be obtained by reversing the order of the elements of *X*^*^.

The notation for the emission probabilities *B* and the *δ* and *ψ* functions has been simplified as a result of not using the durations *D*_*t*_.

**Figure 19:**
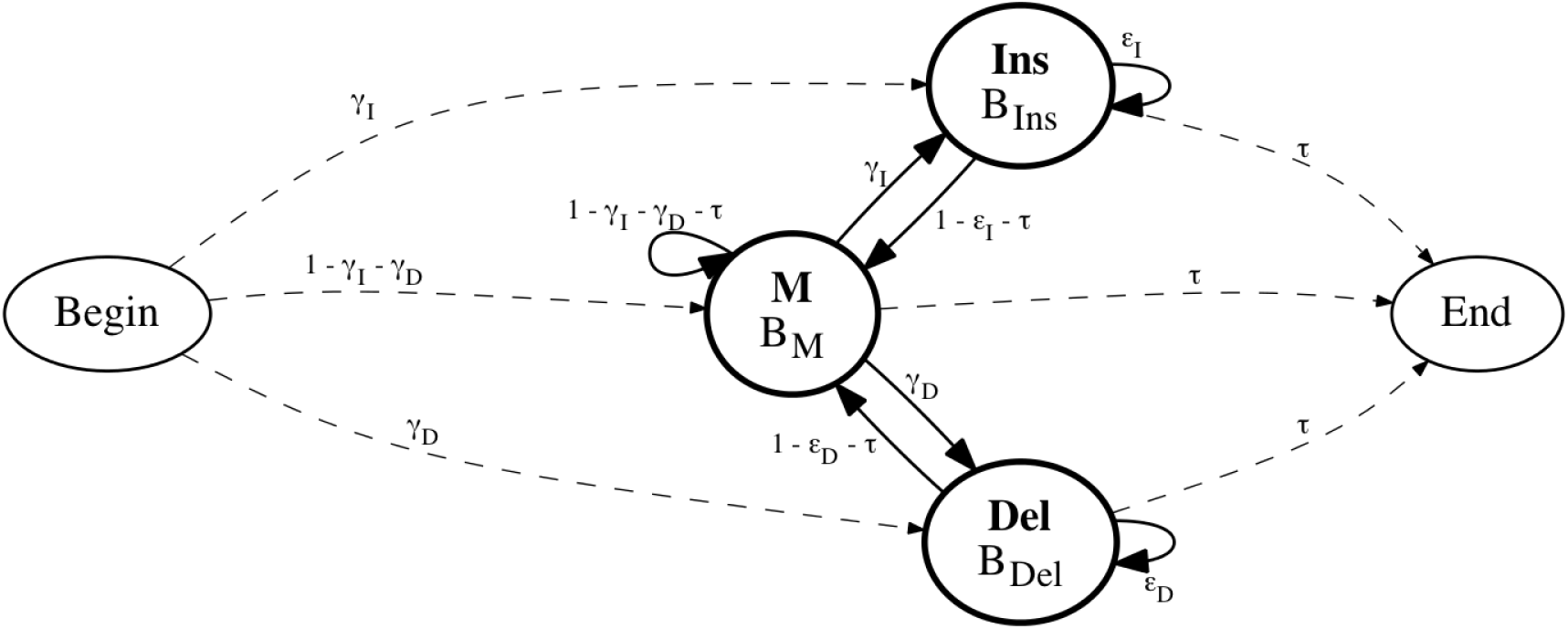
Generalized pair hidden Markov model with Begin and End states added. Hidden states and transition and emission probability distributions. Transitions from and to the *Begin* and *End* states are designated by dashed lines.

#### 3.3.3 Parameter estimation: Viterbi training algorithm

To compute the probability that an observed noisy read *Y* was generated from a true 16S gene sequence 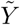 using the Viterbi algorithm, we need to estimate the parameter λ = (*A, B*). For this purpose, we rely on our training set 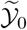 and maximize the conditional likelihood of the reads in 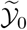 given their corresponding true sequences. Specifically, assuming that errors are introduced independently between reads, we need to compute

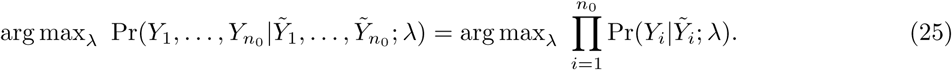

If the pairwise alignments between the reads and their corresponding true sequences were known, that is, if we knew a priori the sequence of matches/mismatches, insertions, and deletions that occurred during the sequencing, we could estimate the transition and emission probabilities by counting the occurrences of each particular event in the training set 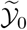. The maximum likelihood estimators of *A* and *B* would then be given by

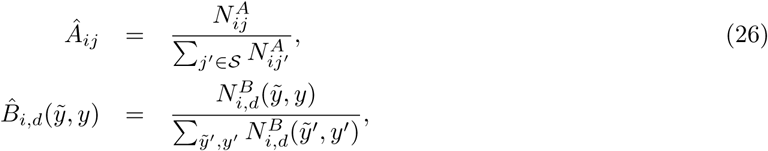

with 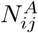 the number of transitions from hidden state *i* to state *j* and 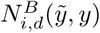 the number of emissions of the aligned pair of nucleotides 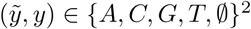 from hidden state *i* with pair of durations *d*.

However, alignments are not known a priori and an iterative procedure must be used. There are two standard methods: the Baum-Welch and the Viterbi training algorithms. The Baum-Welch algorithm is a special case of the Expectation-Maximization (EM) algorithm. The most probable alignment needs to be found for each read in 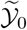 using initial estimates for the transition and emissions probabilities. Then, new values for these probabilities are calculated. The procedure is iterated until some stopping criterion is met. The Baum-Welch algorithm is computationnaly expensive because the forward and backward probabilities need to be computed at each nucleotide in the sequence and for each read in 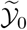 in order to find the most probable alignment. We use the Viterbi training algorithm instead. In this approach, the most probable alignment is computed for each read in our training set using the Viterbi algorithm, with initial values for the parameter λ. Then, Equation (26) is used to estimate the transition and emission probabilities.

Note that unlike the Baum-Welch algorithm, the Viterbi training algorithm does not maximize the conditional likelihood as in Equation (25). Instead, it seeks the value of λ that maximizes the contribution of the most probable alignments to the likelihood

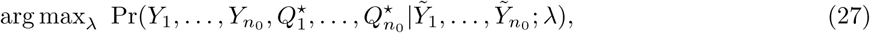

where 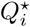 denotes the most probable alignment between a read *Y_i_* and a true 16S gene sequence 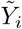. Probably for this reason, the Viterbi training algorithm is known to perform less well in general than the Baum-Welch algorithm. However, as we use the Viterbi algorithm to find the most probable alignment 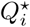 instead of computing the likelihood over all possible alignments (Equation (11)), it is satisfactory for our purpose to estimate the parameter λ with the Viterbi training algorithm instead of the Baum-Welch algorithm [ref].

##### Incorporating quality values in transition probabilities

Often, DNA-sequences obtained from high-throughput sequencing instruments come with base- and read-level quality values. These values are estimates of a base- or read-level error rate. In a PacBio CCS read, we make multiple observations of the same base, in both its forward and reverse-complement context. For each of these observations, an estimate of the base-error probability is available. A CCS-base error rate is computed by averaging the base-error estimates from each observation. The read-level error rate is an estimate of the reads accuracy when aligned to the true template that it was sequenced from. We use the CCS-base Phred quality values (QV) during estimation of transition probabilities where the Phred quality value is defined as — 10*log*_10_(CCS-base error rate).

We let the GPHMM transition probabilities from state *M* to state *Ins* and *Del*, *δ_I_* and *δ_D_*, respectively, depend on the QV of each read in 𝒚_0_. The procedure described below for *δ_I_* can also be used for *δ_D_*.

At each iteration of the standard Viterbi training algorithm, an estimator of *δ_I_* is computed as

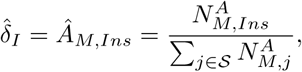

where 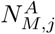 is the number of transitions from state *M* to state *j* ∈ 𝒮 occurring in the entire training set 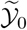. As we want to model *δ_I_* as a function of QV and each read *Y_i_* has a different QV, *QV_i_*, then an insertion probability *δ_I_* (*Y_i_*) now has to be estimated for each read separately.

We assume that the number of insertions 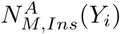 in read *Y_i_* has the binomial distribution

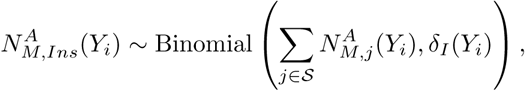

where the insertion probability *δ_I_*(*Y_i_*) is modeled using a generalized linear model (GLM) with logit link function [2] [62],

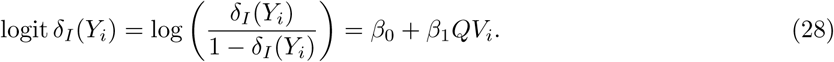

The maximum likelihood estimators of *β*_0_ and *β*_1_ can be obtained using the R function `glm` with `family = binomial()`, thus yielding an estimator 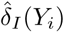 of the insertion probability for each read.

##### Consensus sequence

To estimate the transition and emission probabilities in λ, we need to know the exact true sequence 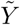 that generated each read *Y* in 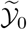 in order to determine the number 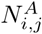 of transitions from state *i* to state *j* and the number 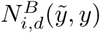 of emissions of the aligned pair of nucleotides 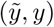. However, we actually do not know the corresponding exact true sequence 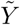 of a read *Y*, but only the bacterium *Z* ∈ 𝓑 it was generated from. Then, we need to consider two cases.

1. When bacterium *Z* has only one reference sequence *r_j_* in the database 𝓡, we assume that the true sequence 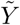 is the same as the reference sequence 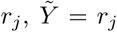. Note that as *r_j_* can belong to several bacteria, we have *Z* ∈ *b*(*r_j_*).
2. When bacterium *Z* has several reference sequences in 𝓡, we need to compute a consensus sequence for the different 16S genes. In our database, sequences from the same bacterium are very similar. When at a given base all sequences have the same nucleotide, the consensus sequence is assigned that nucleotide. For locations at which nucleotides differ between sequences, an “N” is incorporated in the consensus sequence to indicate ambiguity. The function `consensusString` from the R package `msa` [9] is used to perform a multiple sequence alignment using ClustalW (with default parameters) and compute a consensus sequence for each bacterium in our training set 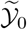. Below is an example of a consensus sequence built from three different reference sequences {*r*_1_, *r*_2_, *r*_3_}.

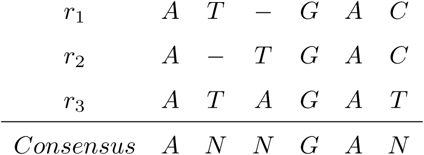

#### 3.3.4 Computation

Computation of a single probability that a read *Y* was generated from a true sequence 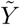 using the Viterbi algorithm involves two nested ‘for’ loops, with the number of iterations being the length of the read times the length of the true sequence, 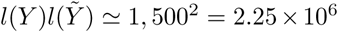. As expected, this computation is slow in R. To reduce computation time, the Viterbi and Viterbi training algorithms were coded using R package `Rcpp`. The computation was also parallelized with sixteen cores using the R function `mclappply` from the package `parallel`. Finally, to avoid underflow problems during the computation, the logarithm of the quantities in Equation (18) was used.

We implemented the Viterbi and Viterbi training algorithms in the R package `gphmm` available on CRAN. Up-to-date code is also available on github at https://github.com/fperraudeau/gphmm.

### 3.4 Estimation of Bacterial Abundances

#### 3.4.1 Maximum likelihood estimator

As mentioned in Section 1.5.3, a natural estimator of the bacterial frequencies π is the maximum likelihood estimator (MLE), defined as

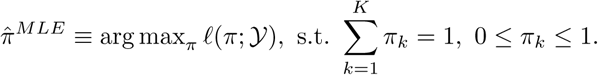

However, due to the log of sums in Equation (5), it is clear that no closed form exists for the MLE of π. Adapted numerical optimization techniques may be used, by noting that the log-likelihood function has the following general form:

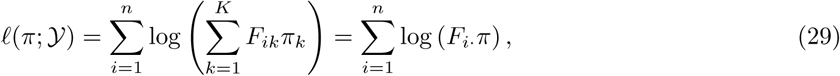

where *F* is an *n* × *K* matrix with 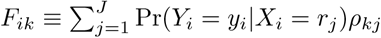 and *i*th row denoted by *F_i._*. Setting partial derivatives over π equal to zero, one has

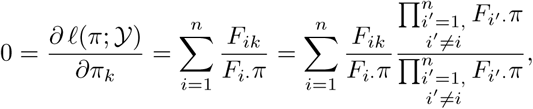

so that

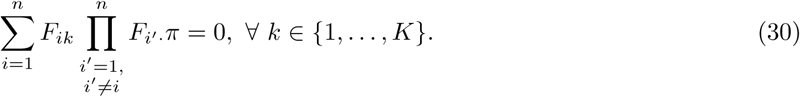

Obtaining the MLE of π therefore involves solving a linear system of *K* equations in *K* − 1 unknowns, summarized by *G*π = 0, where *G* is a *K* × *K* matrix with 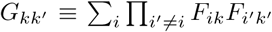. As *det*(*G*) ≠ 0, this system has a non-trivial solution. Numerical optimization techniques can then be used to estimate π. Here, we use the R function `solnp` from the package `Rsolnp` [37].

#### 3.4.2 Penalized estimator

The MLE of π yields many false positives (i.e., 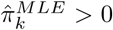 when bacterium *b*_*k*_ is not present in the sample, 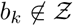). To decrease the number of false positives, we consider estimators based on a penalized log-likelihood function

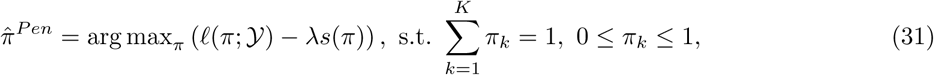

where *s* is a penalty function and λ a tuning parameter controlling the strength of the penalty term (i.e., the shrinking of the estimates towards zero). Note that when 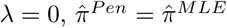.

We focus on two classes of penalty functions:

- Ridge penalty function: 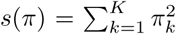. Ridge penalty is intuitive here as we want to decrease the number of false positives, so shrink estimates towards zero. While we know ridge penalized estimation introduces bias, there is a trade-off between bias and variance, so that the root mean square error (RMSE) may overall be reduced.
- Group LASSO [35] penalty function: 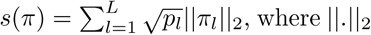 is the *L*_2_ norm, π_*l*_ is the sum of the species frequencies for bacteria from genus *l*, and *p_l_* is the number of reference sequences with genus *l* in our reduced reference database. This penalty should help manage over-representation of certain taxa in our database. For a given taxonomic rank, e.g., genus, certain taxa are vastly over-represented in comparison to other taxa, e.g., *Staphylococcus* is the most represented genus in our database (with 114,146 reference sequences) and is well known to encompass several medically significant pathogens. As the difference in representation has entirely to do with human interests, e.g., studying pathogens, we would like to find a penalty that can perform equally well when assigning to a taxa of 10 versus 100,0 members.

Note that the usual *L*_1_ norm penalty function from the LASSO [86] is not useful here and simply returns the MLE, as, by definition, 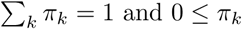.

The penalty parameter λ is selected by Monte-Carlo cross-validation (MCCV), where 90% of the reads are randomly sampled (without replacement) to form a training set and the remaining 10% form a validation set. This process is repeated independently *B* = 100 times, to yield training sets 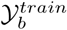 and validation sets 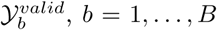. Note that since reads are partitioned independently for each CV fold, the same read can appear in multiple validation sets.

For each penalty parameter λ and each fold *b*, bacterial frequencies π are estimated on the training set 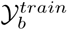 as

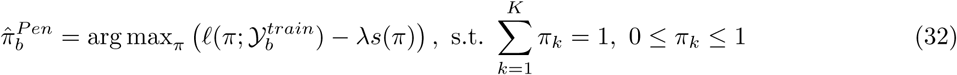

and the log-likelihood evaluated on the validation set 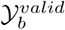

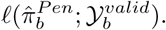

The cross-validated penalty parameter λ is then defined as the maximizer of the average validation set log-likelihood over the *B* different folds

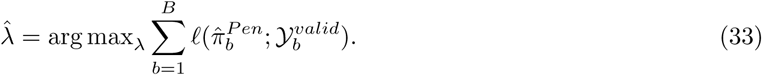

#### 3.4.3 Naive estimator

Additionally, using the same model as described in Section 1.5.3, we propose a naive estimator based on the heuristic that the frequency of bacterium *b*_*k*_ should be proportional to the average probability that the reads in dataset 𝒚 could have been sequenced from bacterium *b*_*k*_, i.e.,

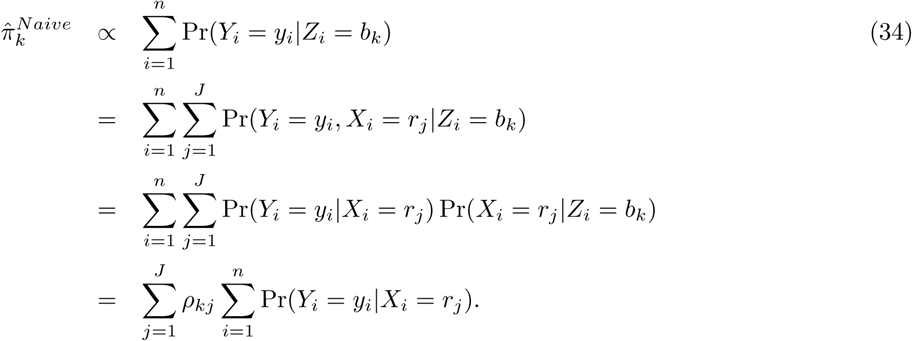

The naive estimators 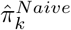 are then normalized to sum to one.

## 4 Results

### 4.1 Simulations

#### 4.1.1 GPHMM read alignment probabilities

As explained in Section 3.2, to reduce the number of computations when calculating the log-likelihood 𝓁(π; 𝒚), we eliminate reference sequences in 𝓡 with small probabilities of having generated the reads in a dataset 𝒚 using the alignment tool Bowtie2. More precisely, reference sequence *r_j_* is eliminated if it has an alignment score lower than the selected minimum alignment score (min-score) for all the reads in 𝒚, i.e.,

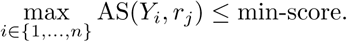

On average, for our 72 simulated datasets 𝒚, 1, 180, 537 reference sequences were eliminated, leaving on average 687 sequences (standard deviation 680) in the reduced versions of the database 𝓡.

Then, for each reference sequence *r_j_* in a reduced database and each read *y_i_* in a simulated dataset 𝒚, the probability Pr(*Y* = *y_i_*|*X* = *r_j_*) that read *y_i_* could have been sequenced from reference sequence *r_j_* is computed using the Viterbi algorithm for a generalized pair hidden Markov model (see Section 3.3).

Figure 20 shows, for two simulated datasets 𝒚, the sum of these probabilities over the *n* reads for each reference sequence *r_j_*, i.e.,

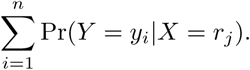

Note that if the reads were sequenced without error, i.e., Pr(*Y* = *y_i_*|*X* = *r_j_*) ∈ {0,1}, this quantity would simply be the number of reads aligned to reference sequence *r_j_*. The two simulated microbial communities in Figure 20 are *Gut 2* and *Vaginal 1* with variability parameter *V* = 100 and *V* = 1, 000 respectively, yielding similar Shannon diversity — Σ_*k*_ π_*k*_ log π_*k*_ (2.4 and 2.6, respectively) and number of distinct simulated species 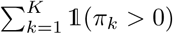 (24 and 27, respectively). Even if the two communities have similar Shannon diversity and number of distinct simulated species (see Figure 9), Figure 20 shows that it is harder to estimate the species frequencies for community *Vaginal 1* than for community *Gut* 2, as the reference sequences in the reduced version of the database are more similar (i.e., the phylogenetics is tighter) for community *Vaginal 1* than for community *Gut 2.* Probably for the same reason, the reference sequences that generated the reads (i.e., simulation frequency is greater than zero for these reference sequences) were included in the reduced version of the database for community *Gut 2* whereas none were included for community *Vaginal 1.* Similar results stand for all our simulated datasets (data not shown), it is harder to match the reference sequences that generated the reads for our simulated *Vaginal* communities than for our simulated *Gut* communities.

**Figure 20:**
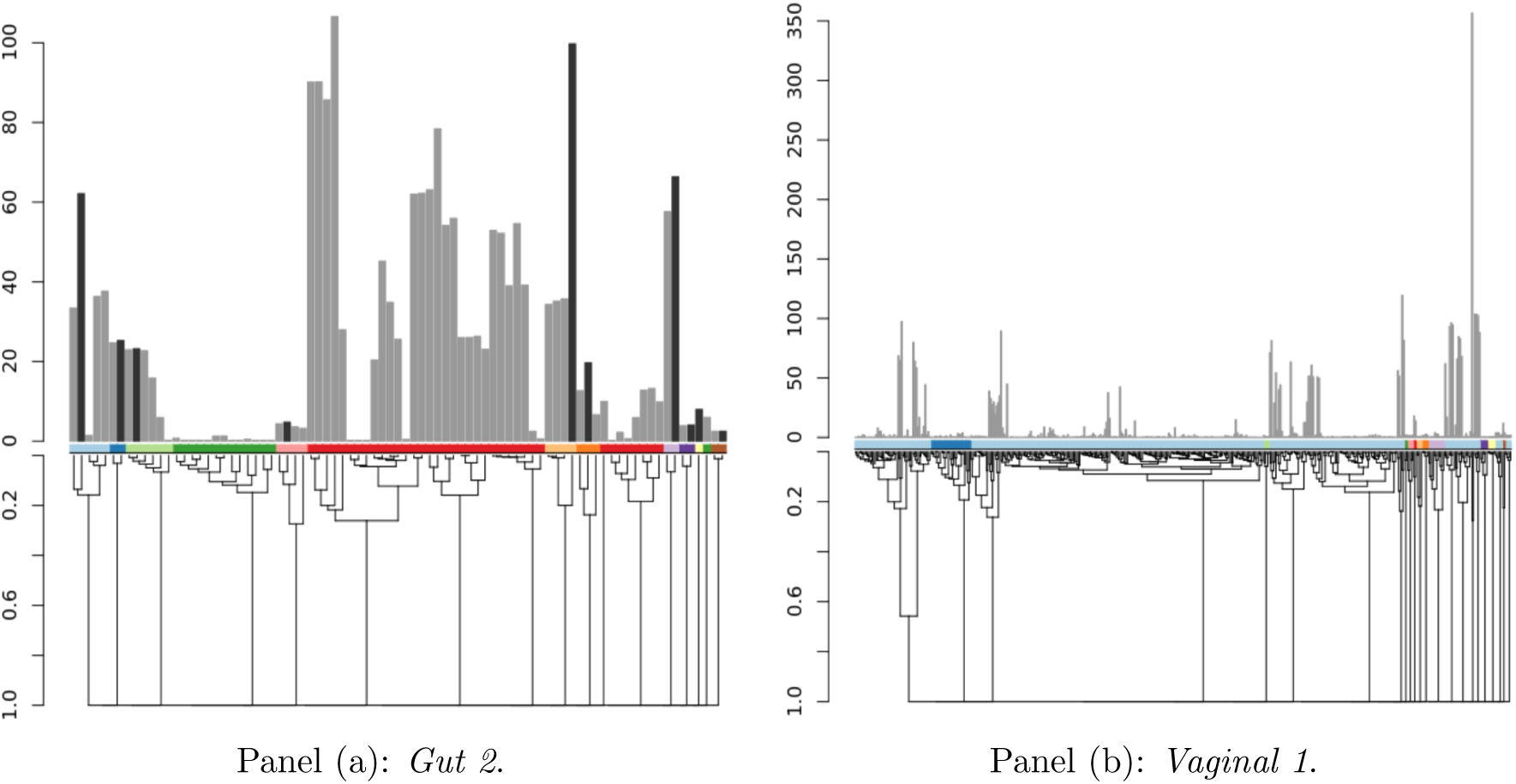
GPHMM read alignment probabilities for reference sequences. Panel (a): Simulated microbial community *Gut 2* with Shannon diversity 2.4 and 24 distinct species. Panel (b): Simulated microbial community *Vaginal 1* with Shannon diversity 2.6 and 27 distinct species. The multinomial sampling distribution was used for both panels. The phylogenetic trees in the bottom panels represent pairwise similarity (based on the Levenshtein distance) between the reference sequences aligning to the reads in a simulated dataset 𝒚. The horizontal colored bars above the phylogenetic trees indicate the genus of each reference sequence, where the same color may correspond to different genera for panels (a) and (b). Each bar in the barplots corresponds to a reference sequence *r_j_* from the phylogenetic tree, its color indicates whether the simulation frequency is greater than zero and its height represents the sum over the reads in 𝒚 of the probabilities that each read could have been sequenced from 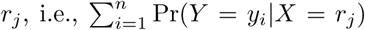. Note that if the reads were sequenced without error, i.e., Pr(*Y* = *y_i_*|*X* = *r_j_*) ∈ {0,1}, the height of the bar would simply be the number of reads aligned to that reference sequence. Note that the y-axes are on different scales for the two panels.

#### 4.1.2 Comparison with SINTAX

The Simple Non-Bayesian TAXonomy (SINTAX) [29] algorithm predicts the taxonomic rank of the reads in a dataset **𝒚** by using k-mer similarity to identify the top hit in a reference database. SINTAX does not require training and, unlike UTAX [25], can be used with full-length 16S reads without running out of memory. Additionally, the SINTAX algorithm provides bootstrap confidence measures for each read and all ranks in the prediction by repeatedly (default 100 iterations) sampling with replacement a set of 32 k-mers (default *k* = 8) from the overall set of k-mers of each read. At each iteration, the reference sequence with the greatest number of k-mers in common with the read is identified and its taxonomy is reported. Then, for each taxonomic rank, the name that occurs most often is identified and its frequency is reported as its bootstrap confidence measure. We compared our estimators to SINTAX results with default parameters and two different bootstrap cutoffs, 0.4 and 0.8. The default bootstrap cutoff is 0.8 and has not as good a performance as a bootstrap cutoff of 0.4.

#### 4.1.3 Performance of bacterial frequency estimators

Let π = (π_*k*_ : *k* = 1,…, *K*) and 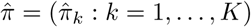, where π_*k*_ denotes the true population frequency of bacterium *b*_*k*_ and 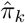 an estimator of this parameter, *k* = 1,…, *K*. The performance of the estimator 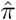 can be assessed using the errors 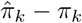 (cf. bias) and the root mean squared error

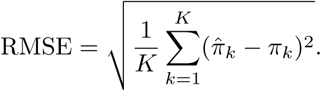

We simulated our read datasets using a variability parameter V inversely related to the Shannon diversity, a metric commonly used to measure the richness and evenness of microbial communities. Indeed, as indicated in the top-right panel of Figure 9, when *V* is large, the Shannon diversity is low, whereas when *V* is small, the Shannon diversity is high. We compared the accuracy and robustness of our estimators as a function of the Shannon diversity instead of the variability parameter *V* as, unlike the artificial variability parameter, the Shannon diversity can be estimated for real datasets.

We also produced receiver operating characteristic (ROC) curves, where the true positive rate (TPR) is plotted against the false positive rate (FPR), with

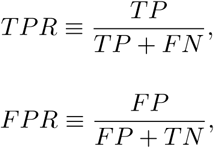

where *TP, FN, FP*, and *TN* stand, respectively, for true positive, false negative, false positive, and true negative, as defined in Figure 23. Each point on an ROC curve corresponds to a different value for the detection threshold e. The greater the area under curve (AUC), the better the estimator.

##### General results

###### Similar robustness for all estimators

To test the robustness of our method, bacterial communities were simulated using three different sampling distributions: the multinomial used in our data generation model and two other distributions, a Poisson distribution and a negative binomial distribution allowing overdispersion. All our estimators showed similar robustness (see Supplementary Figure S1). Not surprisingly, microbial communities simulated from the multinomial and Poisson distributions showed more similar RMSE and sensitivity than the ones simulated from the negative binomial distribution.

###### Abundance estimation is harder for medium Shannon diversity

The Shannon diversity increases with both the richness (i.e., the number of species) and the evenness (i.e., similar relative abundances) of a microbial community. Figure 21 shows that bacterial abundance estimation is harder when the Shannon diversity is neither low nor high. When the Shannon diversity is high, the number of species is high, but the bacterial frequencies are similar. This makes the estimation easier, especially for the ridge and group LASSO estimators which give the same weight to, respectively, all bacteria and bacteria from the same genus. When the Shannon diversity is low, a few bacteria have much higher frequencies than other bacteria, but the number of bacteria is also smaller, which probably eases estimation. On the contrary, when a sample has intermediate Shannon diversity, the imbalance in the bacterial frequencies is not compensated by a small number of bacteria, making the estimation harder.

##### Comparison with SINTAX

###### Accuracy versus sensitivity

Except for our *Vaginal* simulated communities with medium Shannon diversity, SINTAX was superior with respect to RMSE as it had a lower or similar RMSE than our estimators (MLE, Ridge, and group LASSO) (see Figure 21). However, our estimators consistently showed higher sensitivity for a false positive rate greater than 0.05%, meaning that, while SINTAX was unable to detect some bacteria we knew were present in the sample (i.e., simulation frequency π_*k*_ was greater than zero for these bacteria), our estimators showed a smaller number of false negatives 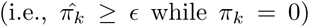, especially for the simulated gut samples. See ROC curves (Figure 22 and Supplementary Figure S2 and), and mean-difference plots (Supplementary Figures S3 to S4).

###### Flexibility

Additionally, our estimators allowed more flexibility than SINTAX estimators to trade false positives against false negatives. When the detection threshold *є* is decreased, the number of true and false positives increases while the number of false negatives decreases. ROC curves (Figure 22 and Supplementary Figure S2) show that the trade-off is smoother for our estimators than for SINTAX estimators.

**Figure 21:**
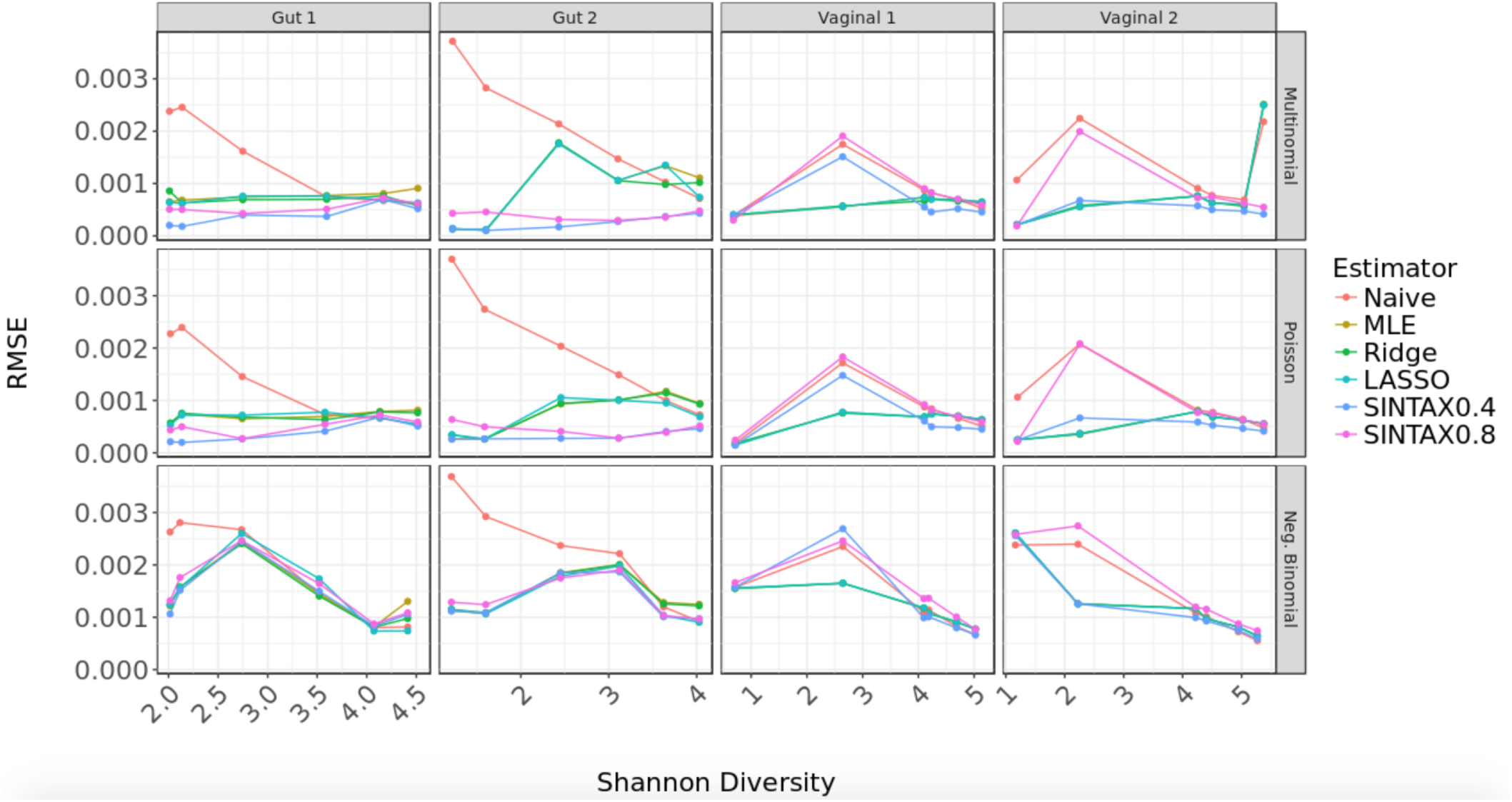
RMSE vs. Shannon diversity. Each panel displays root mean squared error (RMSE) as a function of Shannon diversity (see top-right panel of Figure 9) for a given community and sampling distribution. The colors correspond to our four different estimators, namely, Naive, MLE, ridge, and group LASSO, and to two estimators using SINTAX with bootstrap cutoffs 0.4 and 0.8. For low diversity (Shannon diversity smaller than 3), the MLE tends to have the smallest RMSE. For high diversity (Shannon diversity greater than 3), the MLE, ridge, and group LASSO estimators have similar RMSE. The naive estimator typically has the largest RMSE at all levels of diversity. SINTAX estimators have small RMSE for simulations from the gut communities.

**Figure 22:**
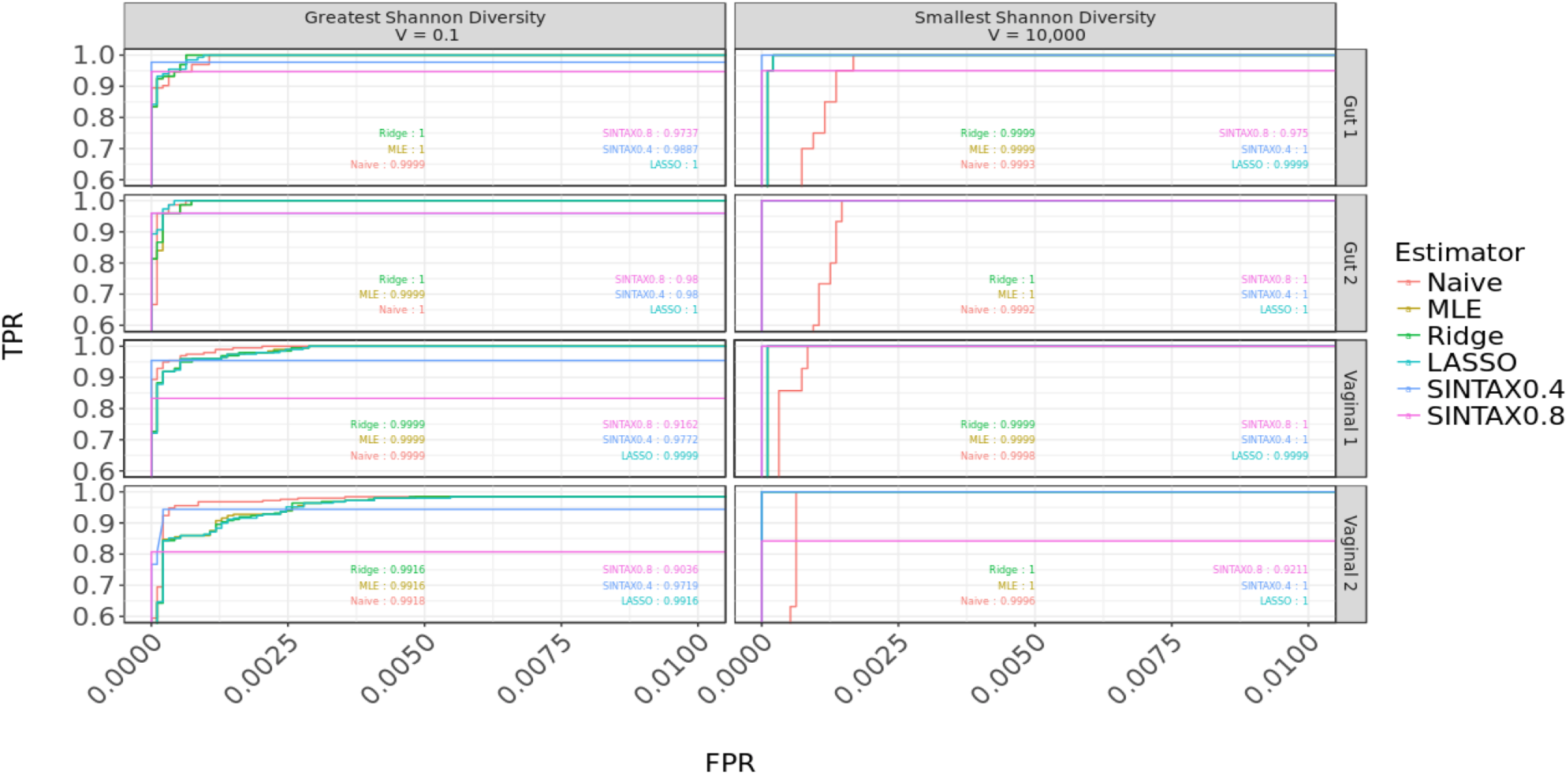
Receiver operating characteristic (ROC) curve. The true positive rate 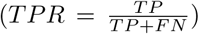 is plotted against the false positive rate 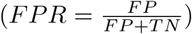 at various levels of detection *є*, where TP, FN, FP, and TN stand, respectively, for true positive, false negative, false positive, and true negative, as defined in Figure 23. Each panel corresponds to a different variability parameter *V* and community (*Gut 1, Gut 2, Vaginal 1,* and *Vaginal 2*) with Poisson sampling distribution. Colors correspond to different estimators.

**Figure 23:**
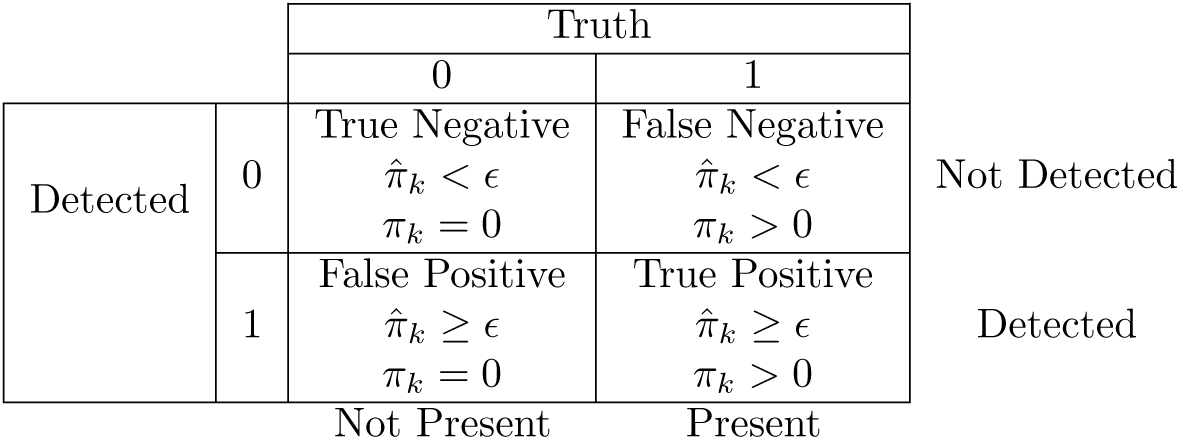
True/false positives and negatives for the different estimators. The table provides the definition of detected/not detected bacteria, present/not present bacteria (“Truth”), and resulting true/false positives and negatives.

**Figure 24:**
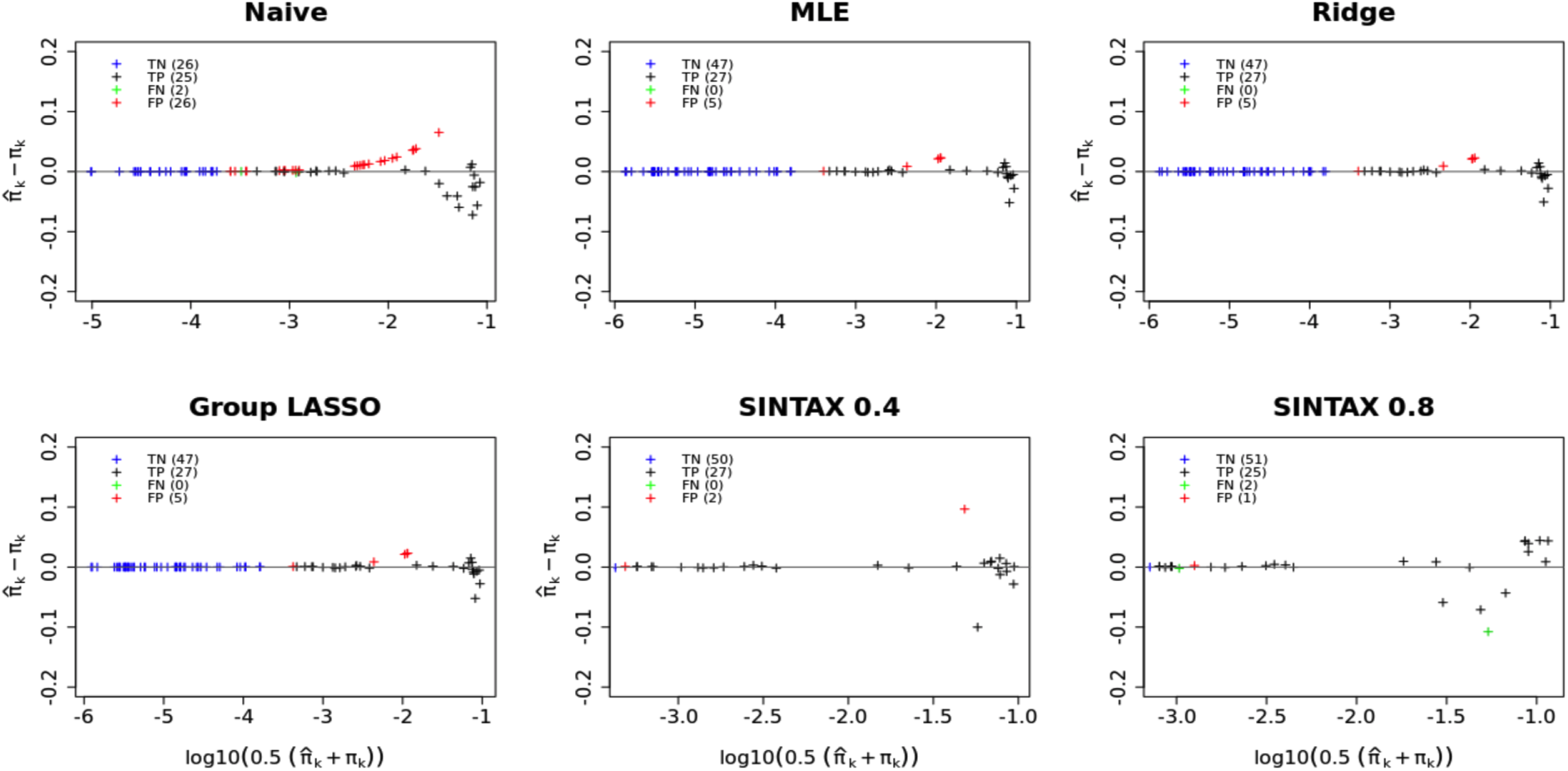
Mean-difference plot of estimated vs. true bacterial frequencies for high Shannon diversity. For simulated dataset *Vag1* with variability parameter *V* = 1000 and Poisson sampling distribution, scatterplots of the differences between estimated bacterial frequencies π̂_*k*_ and true bacterial frequencies π_*k*_ versus the logarithm of the mean of the two values, for the naive, MLE, ridge, group LASSO, and SINTAX estimators. Red points show false positives 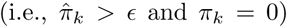, green points false negatives 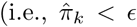 and π_*k*_ > 0), and black points true positives 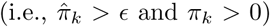 with the number of false positives (FP), false negatives (FN), and true positives (TP) in parenthesis on the upper-left corner (Figure 23). Note that the x-axes are on different scales.

### 4.2 Microbial Mock Community

#### 4.2.1 GPHMM model evaluation

For all the sequencing reads from the mock community, Bowtie2 alignment scores were positively correlated with the length of the alignment whereas GPHMM probabilities were not (see Supplementary Figure S6). It suggests that GPHMM probabilities would be more appropriate for our application than Bowtie2 alignment scores as we want an alignment score or probability to depend on the accuracy of the alignment but not on the length of the reference sequence or the length of the alignment, for alignment lengths varying in a range of the length of 16S gene sequences (1,300 to 1, 600 bp).

For example, reads sequenced from strains *Helicobacter pylori* and *Lactobacillus gasseri* (part of our mock community) had similar read accuracy distributions although the length of their 16S genes (respectively 1,498 bp and 1, 585 bp) is different (see Figure 25). When using Bowtie2 however, the distribution of the alignment scores for strain *Helicobacter pylori* (i.e., the strain with the shortest 16S gene) was shifted to the left compared to the distribution of the Bowtie2 alignment scores for strain *Lactobacillus gasseri* (i.e., the strain with the longest 16S gene). On the contrary, the distributions of the GPHMM probabilities were similar for the two strains showing that for this example length bias was corrected when GPHMM probabilities were used instead of Bowtie2 alignment scores. See Figure 25 and Supplementary Figure S5.

#### 4.2.2 Classification performance of our estimators

We evaluated the performance of our estimators using precision-recall curves and a mock community where by definition the true positives (i.e., the strains present in the sample, that is π_i_ > 0 for the strains in the mock community) are known. At the genus level, RDP classifier had the greatest area under the curve (AUC) and precision at any recall level compared to our methods and SINTAX. See Figure 26 and Supplementary Figure S7. Additionally, all our methods had greater precision than SINTAX at any bootstrap cutoff for a recall between 0.96 and 1. Among SINTAX methods, SINTAX with bootstrap cutoffs greater than 0.8 had better performance than when SINTAX was used with a bootstrap cutoff of 0.5. At the species level, our methods and SINTAX with the bootstrap cutoff of 0.5 had high precision and recall. All of our methods had similar results with MLE having slightly smaller precision than naive, Ridge, and LASSO. As explicitly specified in the FAQ section of the RDP wiki page, RDP classifier was not designed to work at the species level. Thus, we did not compare RDP classifier to the other methods for the species level classification. Overall while at the genus level, RDP Classifier had better performance than our methods and SINTAX, it is not possible to use RDP Classifier to classify bacteria at the species level, and all our methods and SINTAX with a bootstrap cutoff of 0.5 had high precision and recall at the species level.

Note that to evaluate the classification performance of our estimators, we did not use the receiver operating characteristic (ROC) curves which depend on the number of true negatives (i.e., the strains not present in the mock community, that is π_*i*_ = 0 for these strains) which is unknown in this setting. The number of true negatives could be defined as the total number of bacteria present in the universe but absent from the mock community. As the total number of bacteria in the universe is unknown, the true negatives are often set to be the strains present in the database but absent in the mock community. This number is completely subjective to the database used, thus we preferred using the precision-recall curves, independent of the database.

While it was possible to evaluate the classification performances of our method using the mock community, it was impossible to evaluate our performances on quantification as the true bacteria frequencies in the mock community were unknown. The mock community was generated by mixing DNA extracts in equal frequencies (equimolar mixture) before PCR amplification therefore eliminating the DNA extraction bias, but not the PCR amplification bias. It has been studied for decades that PCR amplification efficiencies vary drastically from one bacterium to another introducing bias to estimate bacteria frequencies in microbial communities [1] [3] [68]. To evaluate the performances of our methods on quantification, we would need to create a dataset where true bacterial frequencies are known (see Discussion).

**Figure 25:**
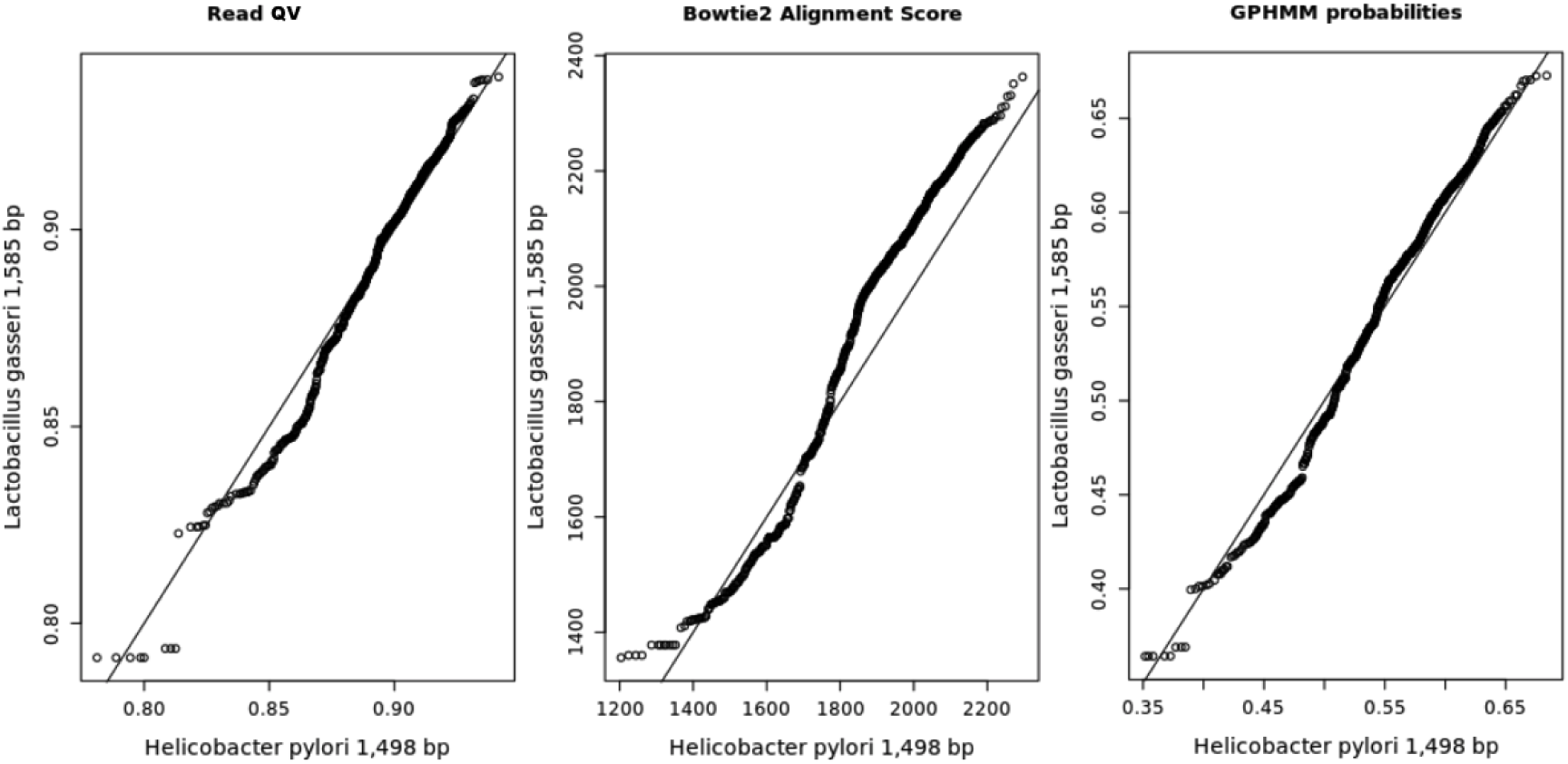
QQplots of expected quality values for the sequencing reads, Bowtie2 alignment scores, GPHMM probabilities for reads sequenced from strains Helicobacter pylori (x-axis) and Lactobacillus gasseri (y-axis). While reads quality values (QV) have similar distributions for strains *Lactobacillus gasseri* and *Helicobacter pylori,* Bowtie2 alignment scores are greater for strain *Lactobacillus gasseri* than for strain *Helicobacter pylori* introducing a length bias. On the contrary, the distribution of GPHMM probabilities are similar for both strains, showing that using GPHMM probabilities can eliminate the length bias. The length of the 16S genes for strains *Helicobacter pylori* and *Lactobacillus gasseri* are respectively 1, 498 and 1, 585 bp.

## 5 Discussion

### 5.1 Limitations to Accurate Quantification

The number of copies and variants of the 16S gene in bacteria genomes varies from one to fifteen [85], but the exact number is usually unknown or uncertain limiting accurate frequencies estimators in microbial samples. The rrnDB database provides estimates of the total number of loci at which a 16S gene could be found in the genome of each bacterium. See section 1.5.2. Although the rrnDB database is a valuable resource, estimates often have a high standard deviation (total standard deviation for the rrnDB database is 2.9) and are not available at the species level. Several single copy housekeeping genes, e.g., those coding for RNA polymerase, concatenated ribosomal proteins, amino-acyl synthetases, or the 60 kDa chaperonin have been proposed as potential phylogenetic markers [89] [43], theoretically avoiding the problems with multiple and variable 16S rRNA copies within bacterial genomes. Protein-coding single copy genes were also successfully used for the assignment of metagenomic sequences, with a better taxonomic resolution than 16S rRNA genes [77]. However, the use of these genes for the analysis of microbial amplicons is limited because the degeneracy of protein sequences makes the design of universal primers difficult. Additionally, the number of 16S rRNA gene sequences in the sequence databases greatly exceeds those of other bacterial genes, which still makes its use preferable by increasing the probability of finding a close hit for taxonomic identification.

**Figure 26:**
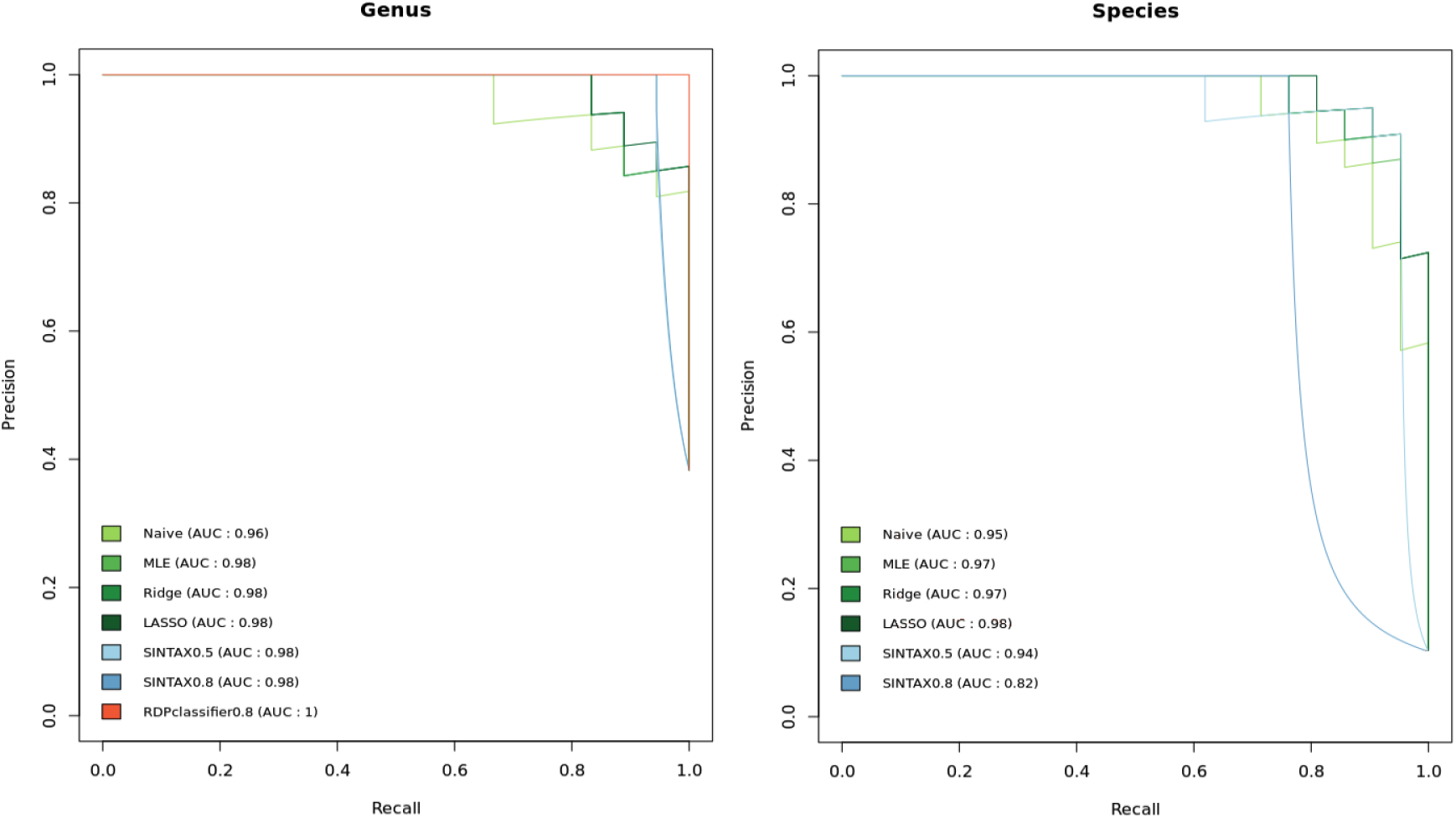
Precision-Recall curve for SINTAX, RDP classifier, and our methods. Left panel shows genus level classification and right panel species level classification. SINTAX is used with cutoff bootstraps of 0.5 and 0.8, and RDP classifier with a cutoff bootstrap of 0.8.

Another limitation to accurate quantification is the fact that DNA extraction efficiency varies from one bacterium to another introducing bias in the quantification estimators, because final sequence read counts may not accurately represent the relative abundance of original DNA fragments. For example, gram-positive bacteria wall have greater strength and rigidity, making it harder to extract the DNA from these bacteria [61]. A solution to estimate DNA extraction efficiencies is to perform an experiment where a known number of cells are lysed and the amount of extracted DNA is measured, using for example fluorometric quantitation. This experiment can be done for a few strains at a time (e.g. pathogens of interest), but it is infeasible to estimate the DNA extraction efficiencies for all the strains in a database. The field of microbiology would benefit from databases where the DNA extraction efficiency is recorded for each strain.

As DNA extraction, PCR amplification efficiencies varies from one bacterium to another affecting quantification accuracy. While this problem has been studied for decades [1] [3] [68], researchers have recently made progress by incorporating random molecular barcodes into PCR primer design. The concept of molecular barcodes is that each original DNA fragment, within the same sample, is attached to a unique sequence barcode which is designed as a string of totally random nucleotides. As a result, sequence reads that have different molecular barcodes represent different original DNA fragments, while reads that have the same barcodes are the result of PCR amplification from the same original DNA fragment. Thus, random molecular barcodes allow to estimate PCR amplification bias by aggregating reads having the same molecular barcode [10] [75].

### 5.2 Limitations Imposed by the Database

In the manuscript, we made the assumption that all bateria in the sample are represented in the annotated reference database. See section 1.5.2. It is obviously a (too) stringent assumption. In [29], Robert C. Edgar accounts for the bacteria not present in the database and estimates two types of false positive error: misclassifications, where a false positive means that a read was assigned to a wrong strain while the true strain is in the database, and over-classifications, where a false positive means that a read was assigned to a strain while the true strain is not in the database. We envision a future report where we would evaluate the over-classification error of our method on different databases.

Another limitation imposed by the use of a database is that bacteria represented in the database are sometimes incorrectly annotated. An accurate method to annotate novel strains would be to use DNA-DNA hybridization (see section 1.1.2) but as this method is time-consuming, labor-intensive, and expensive to perform, fewer and fewer laboratories worldwide perform such assays. Instead, as sequencing cost drastically decreased over the last decade, many studies describing new species are solely based upon their 16S gene sequences. For example, many annotations in the full RDP database are based on the RDP classifier which has a high rate of over-classification errors on full-length sequences (see section 2.2) resulting in incorrect annotations or annotations at low taxonomic levels (e.g., only about 2.6% of the reference sequences are annotated at the species level in the full RDP database). Additionally, some of the reference sequences are probably chimeras (i.e. artifacts made during the PCR process) [21] spuriously increasing the taxonomic diversity of the reference database. To accurately identify bacteria at the species level, it is essential to meticulously choose a complete reference database with correct annotations.

### 5.3 Shotgun Metagenomics

In the past few years, shotgun metagenomics has gained a growing interest in the microbiology field. In shotgun metagenomics, DNA is extracted not only from the 16S gene(s) but from the whole genomes for all the cells in a microbial sample, resulting in a lot of advantages compared to 16S studies. First, there is no need to use specific primers as the whole genomes are sequenced, therefore PCR amplification bias is eliminated. Second, amplicon sequencing typically only provides insight into the taxonomic composition of the microbial community and offers a limited functional resolution [48]. Some methods (e.g., PICRUSt [55]) exists to directly predict the functional profiles of microbial communities using 16S rRNA gene sequences, but is highly dependent on the quality and completeness of the reference database. Third, amplicon sequencing is limited to the analysis of taxa for which taxonomically informative genetic markers are known and can be amplified. Novel or highly diverged bacteria are difficult to study using this approach.

Although shotgun metagenomics indisputably offers a greater potential for identification of strains at the strain level and to perform functional analysis, it presents some challenges [83]. To start with, whole genomes are sequenced instead of just one gene, thus a lot of genomic information need to be sampled from a community. It results in the need of high sequencing depth and an increasing cost to store and analyze the data. Then, while contamination is a challenge general to environmental sequencing studies, the identification and removal of metagenomic sequence contaminants is especially problematic as many of the strains in the microbial samples are unknown [42] [76]. Finally, even if tools have been developed to assemble individual reads into scaffolds or genomes (e.g., Megahit [57], Celera Assembler [66], Omega [40], Spades [8]), classify reads to known genomes (e.g., Kraken [91], Kallisto [78]), bin reads or scaffolds to genome bins (e.g., MaxBin [92], MetaBAT [49]), or even with some additional curation reconstruct closed genomes from metagenomes (e.g. methods described in [5] [11]), these tools have some limitations. Specific exemplary problems include the presence of genes with low evolutionary divergence between organisms or repetitive genomic regions that are larger than a sequencing read [34]. In this context, new statistical and computational tools need to be developed to fully exploit the potential of shotgun metagenomics.

## 6 Acknowledgments

We would like to thank Joey McMurdie and Christian Sieber of Whole Biome for providing valuable feedback on the manuscript.

## Supplementary Figures

**Figure S1:**
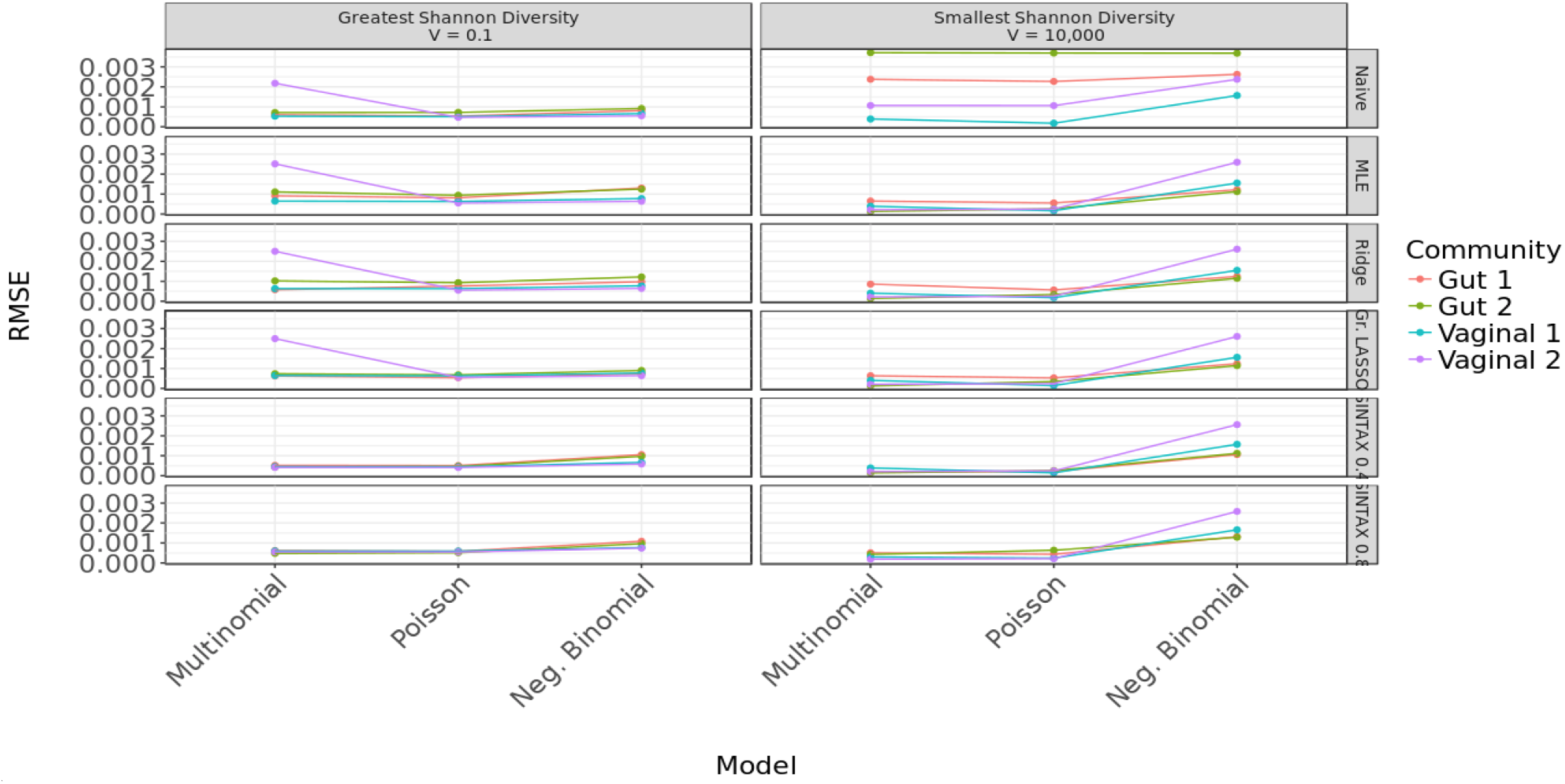
Robustness of estimators. Each panel displays RMSE for each of the three sampling distributions, for a given Shannon diversity and estimator. RMSE is higher for low Shannon diversity (i.e., *V* = 10, 000) than for high Shannon diversity (i.e., *V* = 0.1). The naive estimator is the most robust, as RMSE is similar for the three distributions. The MLE, ridge, group LASSO, and SINTAX estimators are less robust, especially for low Shannon diversity.

**Figure S2:**
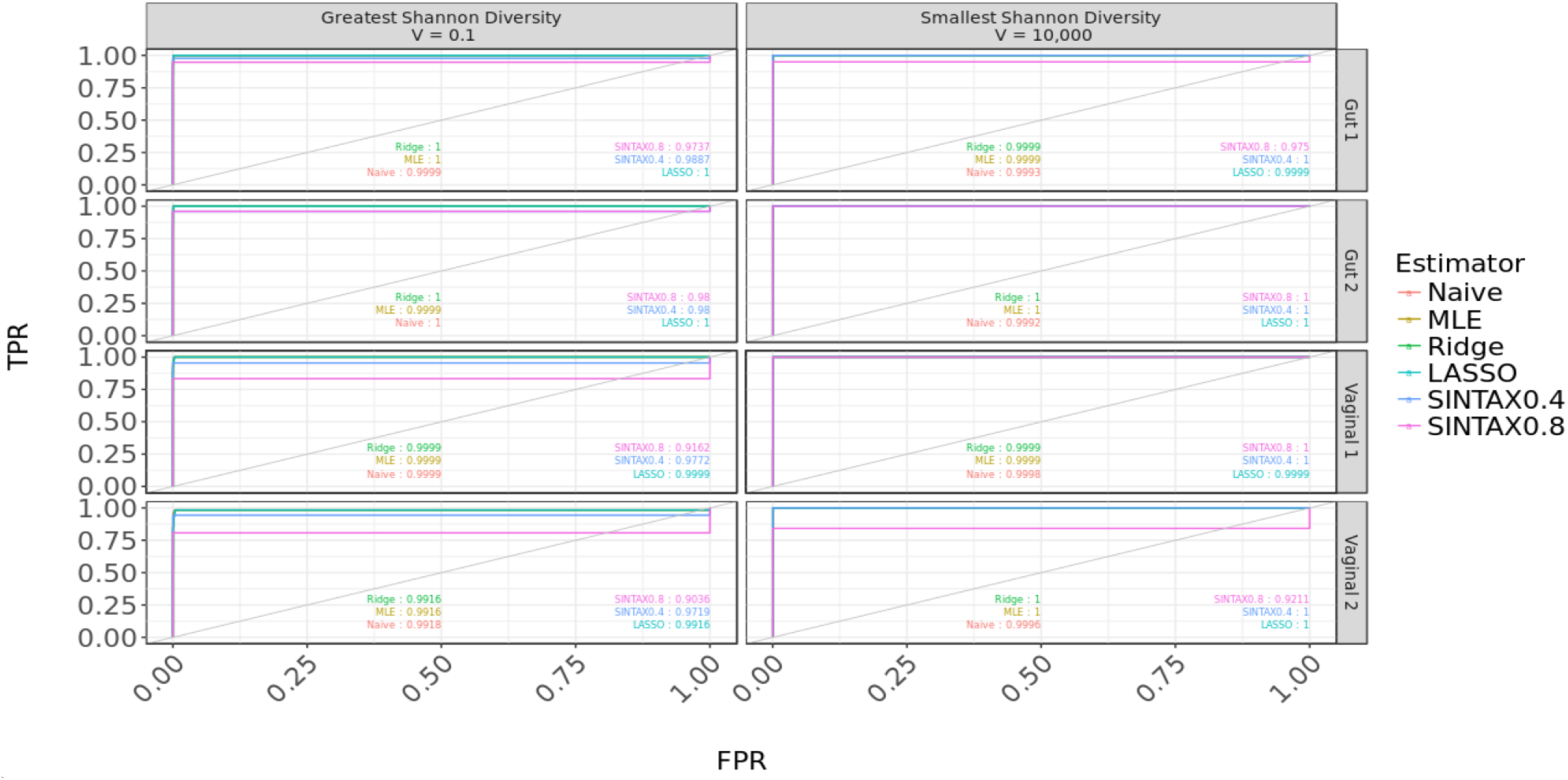
Receiver operating characteristic (ROC) curve. The true positive rate 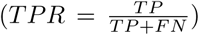 is plotted against the false positive rate 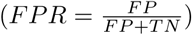 at various levels of detection e, where TP, FN, FP, and TN stand, respectively, for true positive, false negative, false positive, and true negative, as defined in Figure 23. Each panel corresponds to a different variability parameter *V* and community (*Gut 1, Gut 2, Vaginal 1*, and *Vaginal 2*) with Poisson sampling distribution. Colors correspond to different estimators.

**Figure S3:**
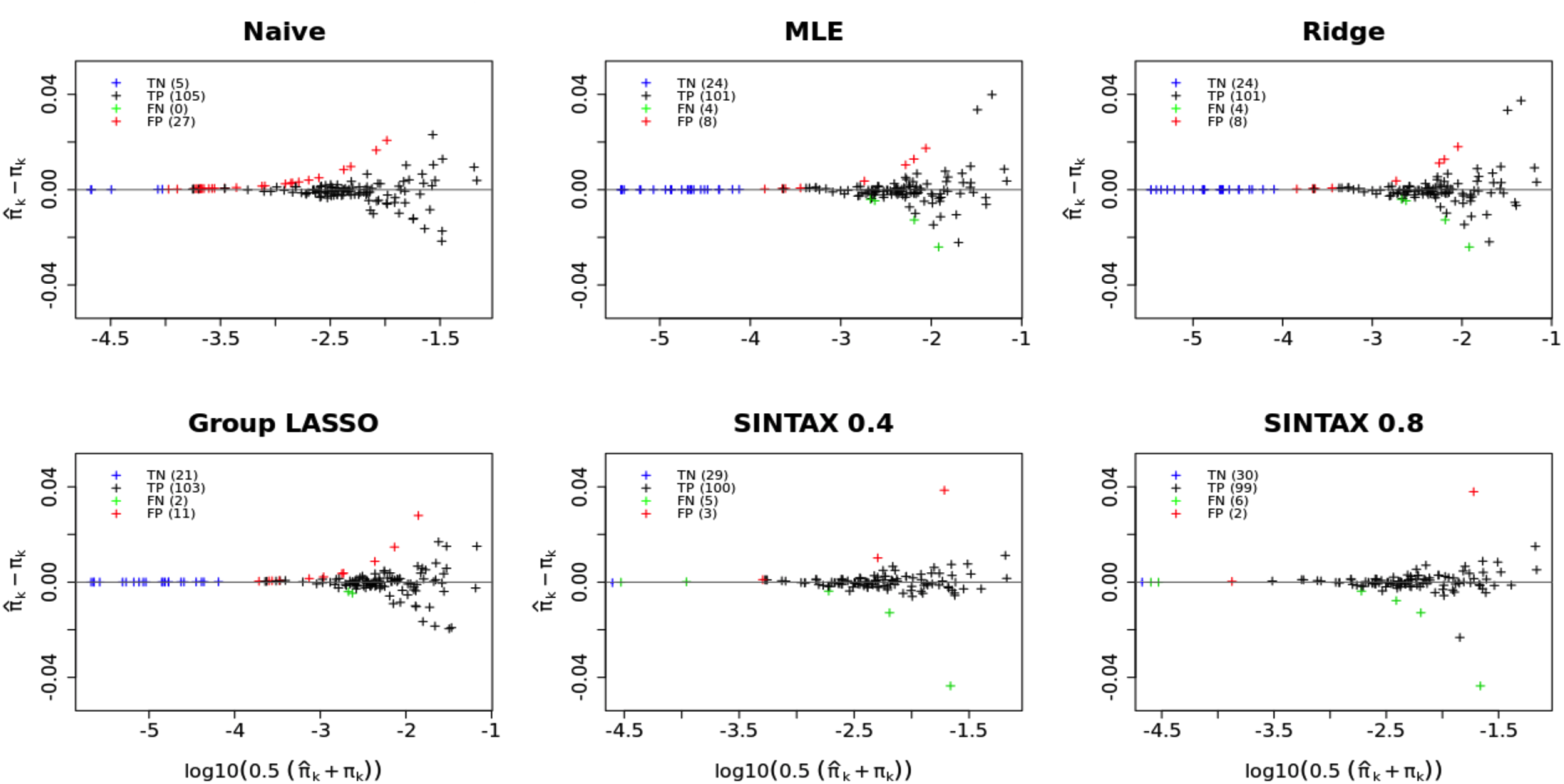
Mean-difference plot of estimated vs. true bacterial frequencies for high Shannon diversity. For simulated dataset *Gut1* with variability parameter *V* = 10 and Poisson sampling distribution, scatterplots of the differences between estimated bacterial frequencies 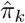 and true bacterial frequencies π_*k*_ versus the logarithm of the mean of the two values, for the naive, MLE, ridge, group LASSO, and SINTAX estimators. Red points show false positives 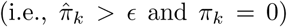, green points false negatives 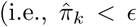 and π_*k*_ > 0), and black points true positives 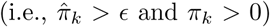 with the number of false positives (FP), false negatives (FN), and true positives (TP) in parenthesis on the upper-left corner (Figure 23). Note that the x-axes are on different scales.

**Figure S4:**
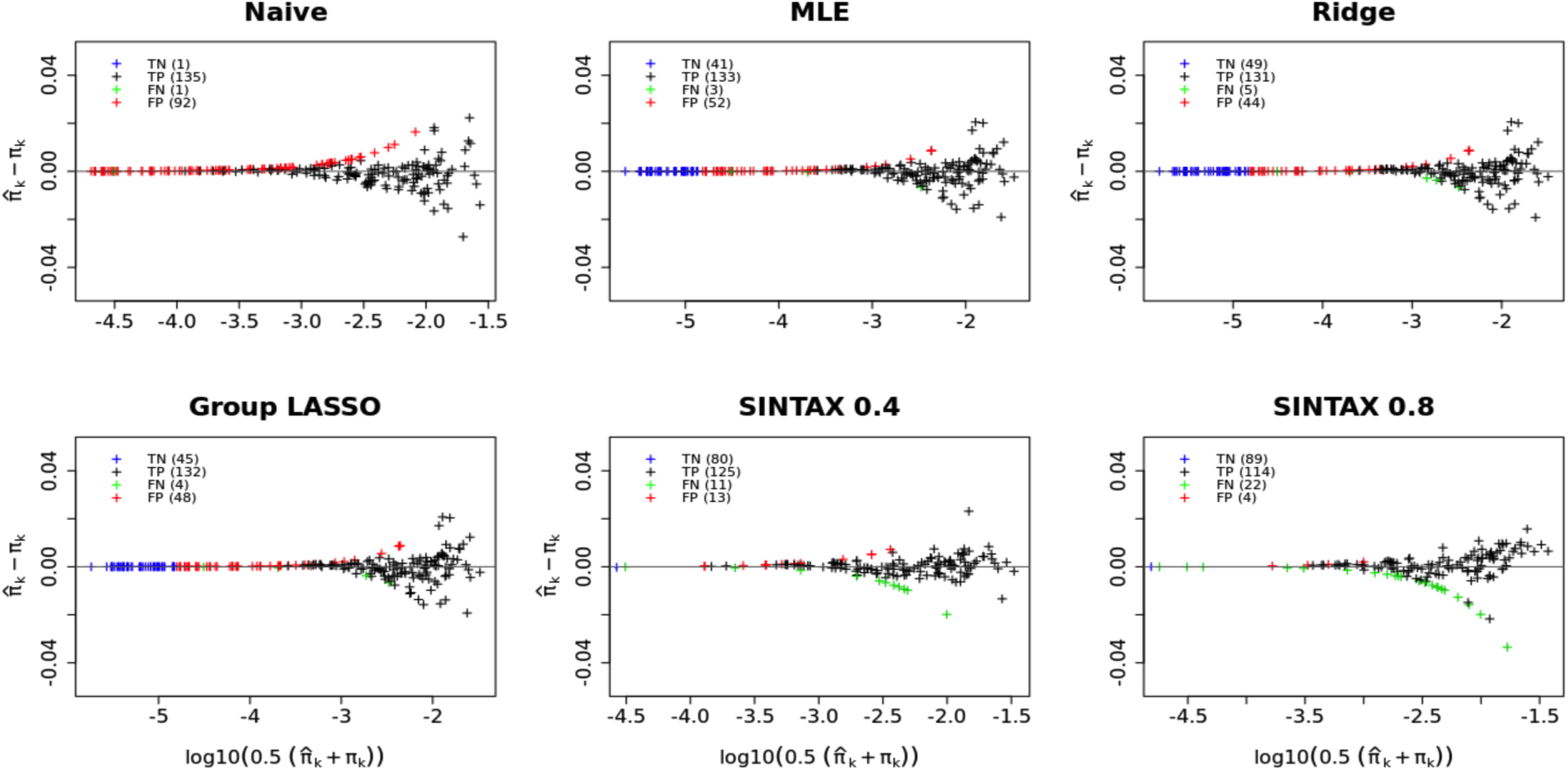
Mean-difference plot of estimated vs. true bacterial frequencies for high Shannon diversity. For simulated dataset *Vag2* with variability parameter *V* = 10 and Poisson sampling distribution, scatterplots of the differences between estimated bacterial frequencies 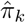 and true bacterial frequencies π_*k*_ versus the logarithm of the mean of the two values, for the naive, MLE, ridge, group LASSO, and SINTAX estimators. Red points show false positives 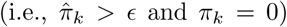, green points false negatives 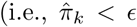 and π_*k*_ > 0), and black points true positives 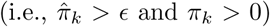 with the number of false positives (FP), false negatives (FN), and true positives (TP) in parenthesis on the upper-left corner (Figure 23). Note that the x-axes are on different scales.

**Figure S5:**
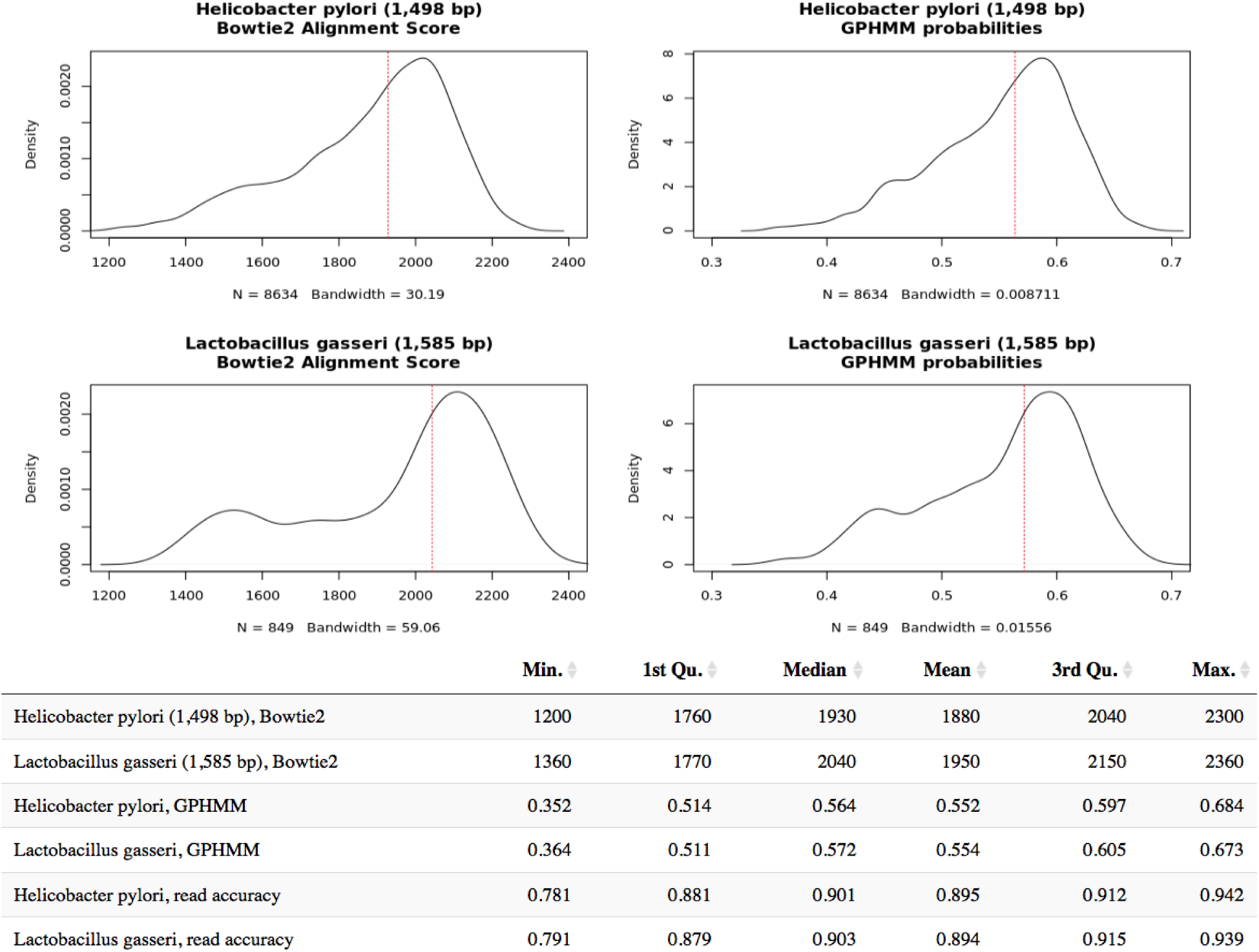
Comparison of Bowtie2 alignment scores and GPHMM probabilities for Helicobacter pylori and Lactobacillus gasseri. Four upper density plots show the distributions of from left to right and top to bottom, Bowtie2 alignment scores for Helicobacter pylori, GPHMM probabilities for Helicobacter pylori, Bowtie2 alignment scores for Lactobacillus gasseri, and GPHMM probabilities for Lactobacillus gasseri. The table summarizes the statistical properties of the four distributions in the upper panel.

**Figure S6:**
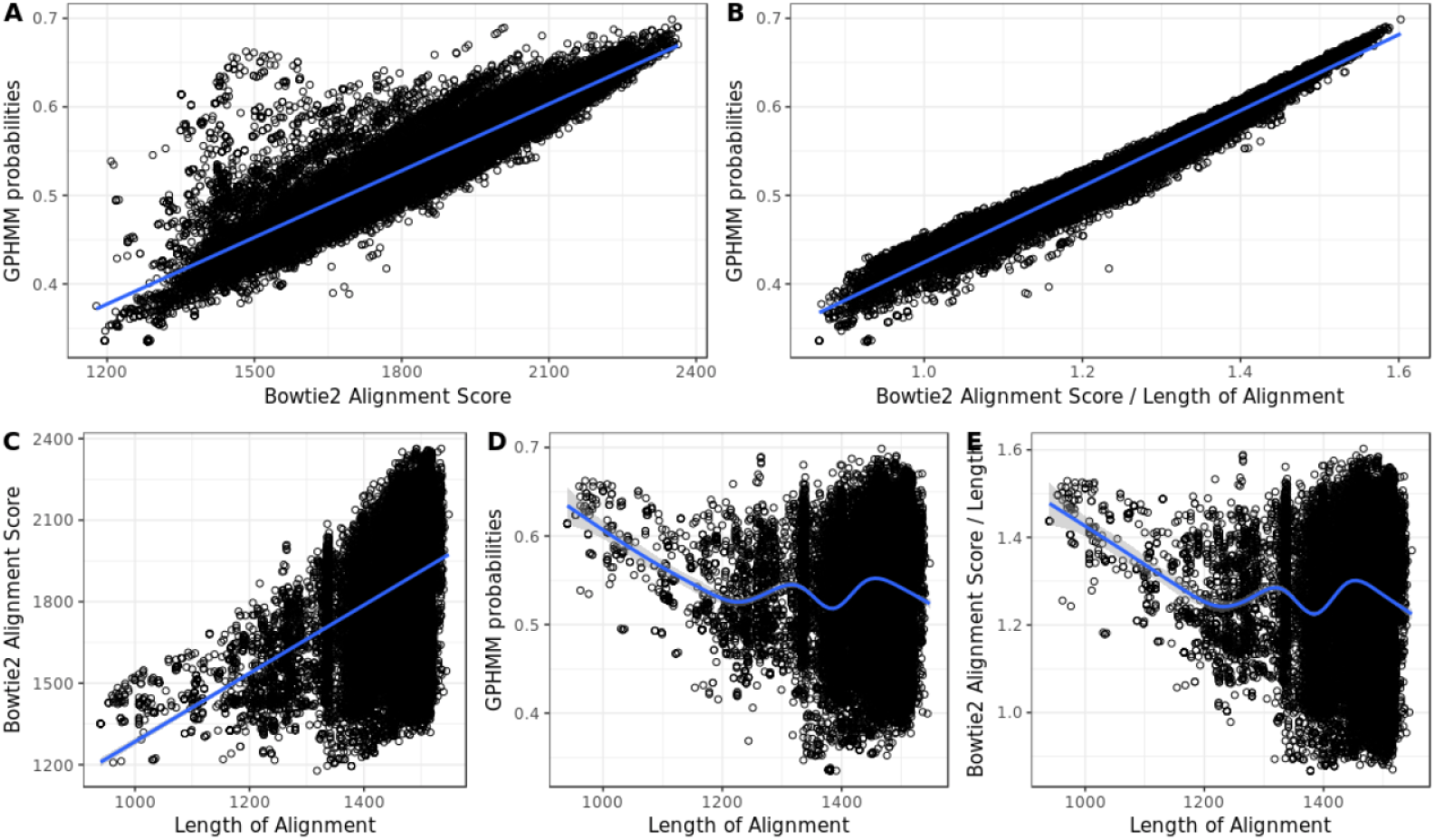
Relationship between Bowtie2 alignment score, GPHMM probability and length of the alignment for all the reads sequenced from the mock community dataset. (A) Bowtie2 alignment scores against GPHMM probabilities, (B) Bowtie2 alignment scores divided by the length of the alignment against GPHMM probabilities, (c) Bowtie2 alignment scores against length of alignment, (D) GPHMM probabilities against length of alignment, and (E) Bowtie2 alignment scores divided by length of alignment against length of alignment.

**Figure S7:**
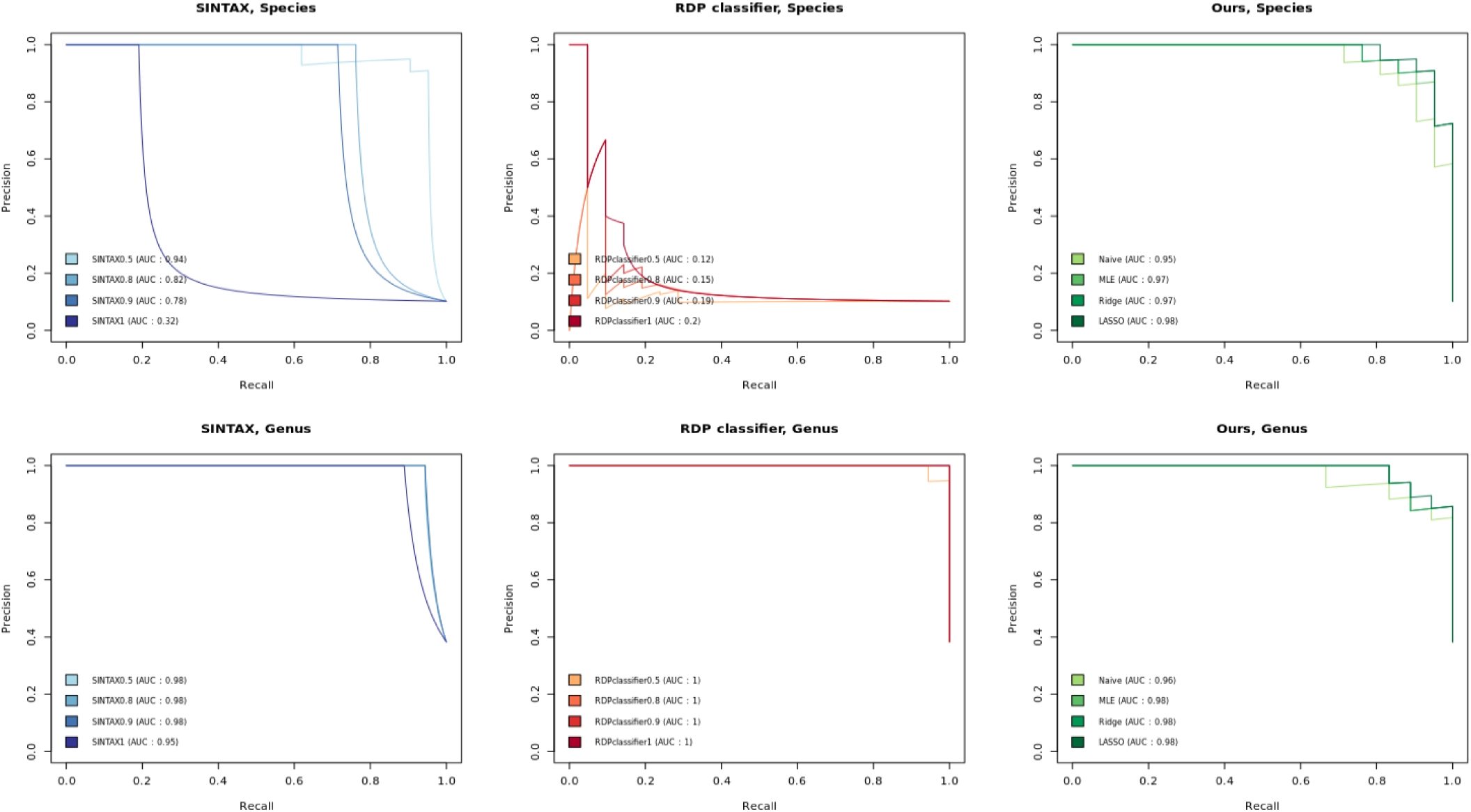
Precision-Recall curve for SINTAX, RDP classifier, and our methods. SINTAX and RDP classifier are shown for bootstrap cutoff of 0.5, 0.8, 0.9, and 1. Plots in the first row show species level classification and plots in the second raw genus level classification.

## Footnotes

1 The DNA of one organism is labeled, then mixed with the unlabeled DNA of another organism to be compared against. The mixture is incubated to allow DNA strands to dissociate and reassemble, forming hybrid double-stranded DNA. Hybridized sequences with a high degree of similarity will bind more firmly and require more energy (i.e., higher temperature) to be separated.

